# Generalists link peaks in the shifting adaptive landscape of Australia’s dragon lizards

**DOI:** 10.1101/2025.08.27.672751

**Authors:** Ian G. Brennan, Natalie Cooper, Joanna Sumner, Sarin Tiatragul, Leonardo G. Tedeschi, Elizabeth S. Broady, Jane Melville, J. Scott Keogh

## Abstract

The adaptive landscape is a flexible concept in biology which helps map entities onto an eco-evolutionary topography. A common practice is clustering species to identify adaptive peaks but the pathways that allow movement into new adaptive zones remain poorly understood. Here we integrate new sequence-capture (>5,000 nuclear markers) and multivariate morphological data to investigate the adaptive landscape of Amphibolurinae dragon lizards (Agamidae). Our well-supported, time-calibrated phylogeny includes over 175 species, including iconic taxa such as the frilled-neck lizard, thorny devil, and bearded dragon. We show that arrival in Australia was followed by rapid expansion of morphological diversity facilitated by generalist habitat use. Ancestral dragons likely had broad niche breadths and intermediate morphologies that provided avenues to subsequent specialization. These findings reveal an evolutionary pathway for adaptive diversification in a major continental radiation.

## Introduction

The adaptive landscape is an indispensable tool for considering and visualizing eco-evolutionary spaces. In adapting Wright’s (1932) genotypic fitness landscape to a phenotypic space, Simpson (1944) provided a shared vocabulary and conceptual bridge between micro-and macroevolution (Arnold et al. 2001). At local scales (species or populations), landscapes help us to map form to fitness. However, at broader scales (species, genera, families), landscapes (often called morphospaces or functional trait spaces) reveal the density and position of peaks, and how clades or communities fit in multidimensional space. This can help in determining the cost of moving between phenotypes or mapping ecology to morphology. Importantly, phenotypic landscapes are dynamic and can vary through time in their characteristics (Marshall 2014). This may dictate the persistence and evolution of the species they represent. Consequently, we can use data from those species to infer parameters of, and changes to, the landscape.

Despite how it is often visualized, the adaptive landscape was never limited to only two or three dimensions. Instead, landscapes can be multidimensional, which intersects with Hutchinson’s (1957) niche hypervolume. In multidimensional space we can assess the volume and density of individual peaks, probe the depths of multidimensional valleys, or detect holes in the landscape hypervolume (Blonder 2016). Higher dimensionality may also provide paths between peaks that might not be identifiable with fewer dimensions. The complex topography of the landscape, including holes, dictates viable paths from existing morphologies into new ones. As such, the ideas of adaptive radiation and ecological opportunity have long been entwined with Simpson’s view of movement across the landscape (Schluter 2000; Simpson 1944; Yoder et al. 2010). However, we still do not understand how new peaks form in disparate regions of the landscape. One hypothesis is that the center of morphological space acts as a hub/bridge to more distant adaptive zones (Dennis et al. 2011; Futuyma & Moreno, 1988). This idea builds the expectation that ecological specialists require extreme morphologies that evolve from ancestors with broad niche breadths and morphologies that are nearer to the morphospace center. With advances in computational power, development of more complex comparative methods, and larger phenotypic datasets, we can begin to unravel historical changes to the landscape and the evolution of new forms (Harmon et al. 2005). Importantly, we can ask which axes have shaped the landscape at different points in time and how these changes are linked with shifts in ecology, habitat, and/or geographic range.

While biologists have been visualizing the adaptive landscape and determining the relative heights of peaks since Simpson, comparatively little empirical attention has focused on how lineages populate new peaks and the landscape changes through time (but see Marshall 2014; Burin et al. 2023). This is partly due to the necessity of a high dimensional dataset, well-resolved phylogeny, and an appropriate organismal group. Here we investigate patterns of adaptive landscape change and the role of ecology and expanding ranges in the morphological evolution of Australo-Papuan agamid lizards. Amphibolurine agamids consist of ∼130 species with varied and diverse ecologies, habitat preferences, and highly imbalanced biogeographic richness. The majority of species (∼110 spp.) are endemic to Australia, where their distributions span the continent’s driest deserts and wettest rainforests. Their morphologies reflect these diverse habitats, ranging from spiny forest dragons, squat desert pebble mimics, and include iconic species such as the frilled-neck lizard (*Chlamydosaurus*) and thorny devil (*Moloch*). To match, they show a range of ecologies from cryptic arboreal species to sand dune burrowers. These varied ecomorphologies provide an ideal opportunity to study the tempo and mode of evolution and the processes driving phenotypic change.

Here we present and test the idea that specialist peaks in the adaptive phenotypic landscape are linked through ancestors that were morphologically intermediate habitat generalists. We began by first reconstructing the phylogenetic relationships of Australo-Papuan dragon lizards using a phylogenomic dataset. To investigate morphological diversity we collected an extensive phenotypic dataset that summarizes broad axes of variation across the head, body, limbs, and tail. We use phylogenetic comparative methods to assess the tempo and mode of trait evolution and from this present a picture of the evolution of amphibolurine dragon lizards. Across the dragon body, many traits show temporally and phylogenetically heterogeneous evolutionary histories. Those traits with evidence of varied evolutionary processes often exhibit large shifts that result in new morphologies. These morphological jumps are better described by biased directional changes which suggest selection, rather than increased evolutionary rates. Lastly, we identify that the diversity of forms of Australian dragons likely evolved from ancestors with broad niches that filled space in the adaptive morphological landscape and acted as intermediates connecting an otherwise patchy landscape.

## Materials and Methods

### Molecular Data Collection and Processing

Molecular sampling used the Squamate Conserved Loci (SqCL) kit (Singhal et al. 2017), which comprises ∼5,000 ultra-conserved elements, ∼400 anchored hybrid enrichment loci, and ∼40 legacy genes commonly used in squamate molecular phylogenetics. Sample collection and laboratory preparation were completed as part of the Australian Amphibian and Reptile Genomics initiative (AusARG) following the protocol outlined in Tiatragul et al. (2023). Raw sequence reads are available from the Bioplatforms Australia Data Portal (https://data.bioplatforms.com/organization/ausarg). We processed raw sequence data using the *pipesnake* workflow (Brennan et al. 2024a). A reproducible list of all software and versions used is included in the Supplement.

### Phylogenetic Analyses

We estimated individual genealogies for our sequence-capture data (n = 5441) under maximum-likelihood in IQ-TREE2 (Minh et al. 2020) allowing the program to assign the best fitting substitution model using ModelFinder (Kalyaanamoorthy et al. 2017), then performed 1,000 ultrafast bootstraps (Minh et al. 2013). We then estimated the species tree using the coalescent-consistent methods wASTRAL-hybrid and ASTRAL-IV (Zhang & Mirarab 2022), using IQ-TREE2 gene trees as input. We used two ASTRAL methods to take advantage of the way wASTRAL-hybrid weights branch lengths and support values in estimating the species tree, and how ASTRAL-IV allows multiple individuals per species and estimates both internal and terminal branch lengths in substitutions-per-site (as opposed to coalescent units for internal branch lengths and static terminal branch lengths in wASTRAL). To quantify topological signal from individual gene trees we calculated gene concordance factors using IQ-TREE2 and focused on intergeneric relationships using a single representative for each genus.

### Divergence Dating

To estimate divergence times among taxa we applied a series of fossil and secondary calibrations in MCMCTree (Rannala & Yang 2007). Our methodology is described at length in the Supplement. Briefly, we partitioned AHE loci by rate and removed third codon positions. We estimated approximate likelihoods and ran four replicate analyses which we compared for convergence before combining to summarize divergence times. We ran a prior-only analysis to determine the effective priors for comparison against posterior age estimates (Fig.S1).

### Morphological Data Collection

We collected 19 linear measurements for 128 amphibolurine lizard species from 482 museum samples (x̅ = 5 per spp., min. = 1, max. = 8). Linear measurements were collected by one author (IGB) using Mitutoyo digital calipers (product 500-763-20). These measurements aimed to capture the gross morphology of the lizard body plan and are distributed across the head, body, limbs, and tail (Fig.S2). Definitions of trait measurements and adjustments are summarized in Table S3. Data processing and analyses used R (R Core Team 2024).

### The Evolution of Habitat Use

To address the concepts of generalism and specialization, we focused on niche breadth. Specialization can occur on varied axes, with a given taxon being a generalist on one (e.g. habitat type), but a specialist on another (e.g. diet) (Poisot et al. 2011). We focus on the fundamental Eltonian niche of species morphologies and the realized niche through substrate usage (Devictor et al. 2010). We first discretized surface features and microhabitats used by dragon lizards into six categories which have different functional demands: aquatic, rock, terrestrial, low structure, high structure, and arboreal (see Supplement for details). Many species regularly use more than one substrate and so we scored all species for all traits. We then modelled the evolution of substrate usage by tracking acquisition and loss of character states allowing polymorphic conditions (e.g. a species found on rocks and the ground would be “R+G”). Using this matrix we fitted a series of Markov models using *phytools* (Revell 2024) to compare theoretical pathways for the evolution of substrate usage, niche breadth, and specialization in dragons. These included stepwise (increasing or decreasing breadth by one state), unordered, and ordered transitions (terrestrial <-> low structure <-> high structure). Importantly, we applied derivatives in which state acquisition and loss are allowed separate transition rates (see Supplement for more details) to all eligible models. To estimate ancestral states we used a model averaging approach that accounts for uncertainty by incorporating model weights.

### The Evolution of Morphological Diversity

To identify the mode of individual morphological traits we fit a series of models ranging from an unbiased random walk to rate-variable and trended. The most basic model—Brownian Motion (BM)—allows phenotypes to evolve through incremental random change. The most complex—*BayesTraits V4*’s fabric model (Pagel et al. 2022)—also follows a process of gradual change, but allows rates to shift at discrete points (changes in *evolvability*, *v*) on the tree and allows traits to change rapidly along individual branches through directed evolution (*trends*, β). Using *BayesTraits V4* we fit these models and their derivatives (*fabric* with no *v* parameter, *fabric* with no β parameter) and estimated their marginal likelihoods using a stepping-stone sampler (500 stones each for 5,000 generations).

*BayesTraits V4* implements a reversible-jump MCMC sampler, which we specified to run for 100 million generations with 10 million generations of burn-in. We summarized the output to check for convergence with custom scripts, and verified that all parameters had ESS > 200. To verify model fit and consistency among runs we ran four separate runs for each trait. Individual model outputs were summarized using the *FabricPostProcessor* software and independent runs were combined using *MergeFabric*.

To track trait values at internal nodes we logged values from the MCMC chain using the *AddTag* and *AddMRCA* features in *BayesTraits* by generating custom control files using functions defined in the Supplement. We also used this script to generate priors on the root state and rate by fitting a single rate BM model using *GEIGER* (Pennell et al. 2014). We summarized posterior estimates of ancestral trait values under varied models using custom scripts included in the Supplement. To track the evolution of dragon morphospace through time we extrapolated trait values linearly along branches at 0.5 million year intervals given start and end values at nodes and constant estimated evolutionary rates. We then reduced observed and estimated ancestral values to three principal component axes and fit multidimensional hypervolumes using *hypervolume* (Blonder et al. 2018) to extract the volume. To determine how the evolution of the empirical hypervolume might depart from expectations we simulated 100 morphological datasets under Brownian Motion and carried out the same exercise. To understand morphological distances between observed and estimated (ancestral) taxa/optima we measured Euclidean distances. We visualized functional morphological spaces using the package *funspace* (Pavanetto & Puglielli 2024).

To identify adaptive peaks across the multidimensional landscape (i.e., morphological regimes) we used an OU-based method that accounts for correlation among traits (*PhylogeneticEM*; Bastide et al. 2018). We used two datasets as input. The first included all 19 non-overlapping morphological traits. Due to the strong correlation of some traits (Fig.S3) we generated a second dataset by reducing dimensionality through principal components analysis (PCA) and choosing the top six principal component axes which accounted for >90% of the cumulative variance, where each component explained >2%. We fit the models across a number of shift values (K = 0-15) and 30 alpha values (ɑ = 0-5) to both datasets. For consistency, we fit several additional schemes to test the reliability of our results (see Supplement). After identifying the preferred number of optima (either 5 or 12; see Results; also called regimes) we used these as discrete characters for *RandomForest* (Liaw & Wiener, 2022) analysis to categorize ancestral taxa according to contemporary optima. Assigning species to ecological guilds is fraught with subjectivity so we relied on our morphological regimes to identify clusters of species that map loosely to existing ecological categorizations (e.g. arboreal, terrestrial) following discussion in Melville & Wilson (2019). RandomForest confusion (error) rates for almost all contemporary optima were sufficiently low (<10%), so we predicted the morphological regime of ancestral taxa (nodes) based on trait values estimated by the best fitting model for each trait. We further explored the possibility of bias in our inferences and model sensitivity through simulations (see Supplement).

### Linking Morphology to Niche Breadth

Ecomorphology predicts a correlation between the ecology and morphology of organisms (Bock 1994; Ricklefs & Miles 1994) and underpins the idea of adaptation. To test the idea that habitat usage in dragons is dictated by morphological position, we used phylogenetically corrected Generalized Linear Mixed Models in *glmmTMB* (Brooks et al. 2017) following the practice of Williams et al. (2026). We fit two model types with differing predictors using the sum of substrate states (breadth) as the response. In the first, we estimated the multivariate distance to the morphological midpoint as the Mahalanobis distance. In the second we fit the first six Principal Component axes (accounting for >90% of variation) and included both linear and quadratic terms to account for expected hump-shaped responses as PC scores become more extreme. Because our response variable consists of positive integers starting at 1, we specified a zero-truncated Poisson distribution.

## Results

### Molecular Data Collection and Processing

Sequence capture using the SqCL kit resulted in assembly of 5441 targets. New sampling recovered a median of 5118 loci, with a concatenated and aligned sequence length of >6 Mbp (Fig.S4). See Supplement for additional sequence statistics.

### Phylogenetic Analyses

Coalescent-consistent topologies estimated by wASTRAL-hybrid and ASTRAL-IV are well resolved with high support across most branches, including good resolution among genera and major clades (Fig.1; Fig.S5). Internal branches separating the Australian genera are short and consecutive rapid splits show high levels of incomplete lineage sorting resulting in low gene concordance factors (Fig.S6). This is notable in the positions of the *Moloch*/*Chelosania* and *Lophosaurus* clades and placement of *Tropicagama*. Additional areas of conflict are limited to interspecific placements of the *Ctenophorus fordi* group, the *Pogona minor* group, and the *Diporiphora pallida* group. In agreement with a recent study (Bruton et al. 2025), we find *Ctenophorus* (*Cryptagama*) *aurita*—a highly morphologically derived lineage—nested within *Ctenophorus*. Our phylogeny also supports the assignments of several frequently confused arid and savannah species, upholding the genera *Amphibolurus, Gowidon, Lophognathus,* and *Tropicagama*. We also provide strong evidence that unites *Rankinia*—the world’s highest latitude dragon—with *Pogona*. Beyond Amphibolurinae, we also provide a strongly supported estimate of subfamilial relationships in Agamidae. We recover a sister relationship between the Agaminae and Draconinae, which together are the sister to the Amphibolurinae. The next closest relatives are the Hydrosaurinae, then Uromastycinae, and lastly Leiolepidinae (Fig.1)

**Figure 1.**
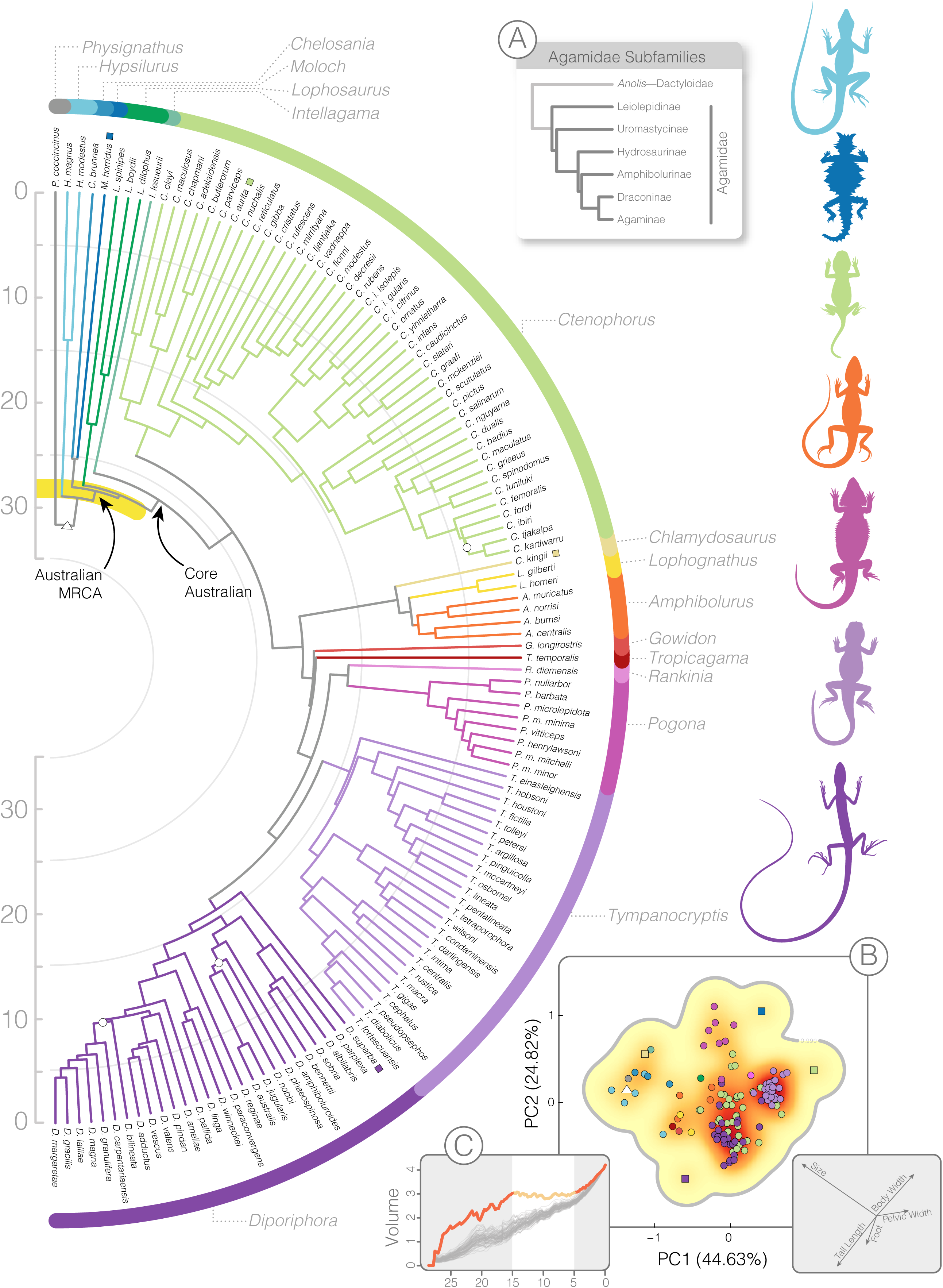
Species-level timetree of the Amphibolurinae, color-coded by genus. The ASTRAL species tree was time-calibrated with MCMCTree. A yellow band highlights a series of rapid splits early in the history of this group. White circles on the tree denote local posterior probabilities <0.95. (A) Relationships among agamid subfamilies and color-coded dragon drawings at right (not to scale) highlight some of the morphological diversity of this group. (B) The spread of species in two-dimensional morphospace (also colored by genus) and primary axes of variation. *Diporiphora* (dark purple) and *Tympanocryptis* (light purple) highlight dense clusters of morphospace, contrasting with morphological outliers indicated by squares in the morphospace and on the tree (*D. superba*, *M. horridus*, *C. aurita*). The estimated root morphotype is indicated by a triangle. (C) The evolution of the morphological hypervolume through time (orange) relative to null expectations (grey) shows periods of rapid expansion (25-15; 5-0 mya) and a period of relative stasis (15-5 mya).

### Divergence Dating

Amphibolurines appear in the late Cretaceous (∼75 mya), separating from their sister the Agaminae/Draconinae. The crown split among amphibolurines does not occur until the Eocene/Oligocene boundary ∼34 mya. Cladogenetic events occur rapidly at the base of the amphibolurine tree, with all major splits among Australian groups occurring between 30-25 mya. With the exception of the clade comprising *Chlamydosaurus, Lophognathus,* and *Amphibolurus*, all extant genera were established by the mid-Miocene ∼15 mya. Rapid speciation events at the base of the amphibolurine tree obscure some relationships. This is likely the result of quick successive splits that occurred in a narrow temporal window, <600 ky (Fig.1).

### Evolution of Niche Breadth

Our discrete trait modelling of the evolution of substrate usage identified best fitting models (rock.order.jump, dAIC = 0.0; two.order.jump, dAIC = 1.5; two.order, dAIC = 2.0) which followed an ordered transition among habitat states (Fig.S13). Preferred models included two transition rates: a habitat acquisition rate (0.0033-0.0037) and a habitat loss rate that was roughly six-to-nine times faster (0.0225-0.0271). These also allowed limited transitions between habitat specialists (breadth = 1), with the exception of transitions from ground to rock-living *Ctenophorus*.

### Phenotypic Evolution

Of 19 morphological traits, 10 were best fit by the rate-variable and trended model from *fabric*, including the summary trait encapsulating size (Table S4). The remaining nine traits were best fit by a random walk model (Brownian Motion). Traits with greater variance and disparity were more likely to fit the *fabric* model (Fig.S7). Ancestral trait values for the 10 traits best fit by *fabric* were extracted and summarized from the mcmc output files. Ancestral trait values for the remaining nine traits were estimated under Brownian Motion. This provided a complete trait matrix for living and ancestral species which we used to quantify ancestral morphologies. Morphospace showed a heterogeneous pattern of expansion with a steep increase from 30-20 ma, before a period of little expansion from 15-5 ma, and another increase in volume from 5 ma to the present (Fig.S8). Early (>20 ma) and middle (>15 ma) morphospace expansions departed from Brownian expectations.

Multi-optima OU models identified either five or 12 morphological regimes. We refer to these groups by their inferred regime identities (R1, R2… R5) or the clade to which they belong (*Tympanocryptis, Moloch*), and consider how these morphological assignments map to ecology and habitat use in the Discussion. Our multi-optima solutions were nested with the same five optima identified in the more conservative model and in the liberal 12 optima model. These optima correspond to arboreal taxa at the base of the tree (R1: *Hypsilurus, Lophosaurus, Chelosania*), two terrestrial clades (R2: *Ctenophorus*; R5: *Tympanocryptis*), an Australian semiarboreal group (R3: *Amphibolurus*, *Gowidon*, *Intellagama*, *Tropicagama*, *Lophognathus, et al.*), and the unique *Moloch* (R4) (Fig.3; Fig.S9). More liberal models added optima which largely corresponded to genera (e.g. *Diporiphora*, *Pogona* + *Rankinia*) and outlier species (*Chlamydosaurus kingii*, *Cryptagama aurita*, *Diporiphora superba*) and optimally identified 12 peaks (Fig.S10). These clusters were largely visible when looking at raw morphological distances among taxa (Fig.S11).

In both optima scenarios (k = 5, k = 12), RandomForest predictions suggested an arboreal-type ancestor for the Amphibolurinae with some uncertainty (k = 5, 66% assignment to arboreal; k = 12, 58%). The core Australian amphibolurine ancestor (most recent common ancestor [MRCA] of *Intellagama* and relatives) is estimated as regime R3 (semiarboreal generalist), however, its immediate ancestor is estimated as regime R1 (arboreal). The amphibolurine MRCA is estimated to have trait values most similar to *Chlamydosaurus*, *Intellagama*, and *Physignathus* (Fig.S11,S12). Simulations show our approach has reasonable accuracy (80%) and no identifiable bias in assigning ancestral morphologies to contemporary regimes, though accuracy varied across nodes and declined with increasing node age.

### Morphology and Niche Breadth

Phylogenetic GLMMs indicate significant effects of distance to the morphological center and PCs 1 and 4 on predicting substrate usage breadth (Fig.5). Niche breadth declines with increasing distance from the morphological center (β = −4.11, SE = 1.22, Z = −3.35, P < 0.001). PC1 (β = −1.09, SE = 0.42, Z = −2.55, P = 0.010) shows a negative quadratic function where substrate breadth is highest at intermediate PC scores. PC4 (β = 2.17, SE = 0.86, Z = 2.53 P = 0.11) shows a positive linear trend with substrate breadth. PCs 2, 3, 5, 6 do not show significant effects (see Supplement for statistics).

## Discussion

Organismal forms in nature are not distributed evenly across the possible phenotypic space. Instead, observed phenotypes often cluster in discrete regions of morphospace, a result of genetic, functional, and/or developmental constraints (Deline et al. 2018). These clumps of morphologically similar organisms identify peaks in the adaptive landscape. Through time, the adaptive landscape may change as conditions vary and species go extinct, persist, and proliferate. The invasion of a new geographic region provides us the opportunity to observe how the adaptive landscape evolves, and to identify the tempo and mode with which new regions of the landscape become populated. Our findings suggest that dispersal of dragon lizards into Australia facilitated a marked expansion in the morphological landscape. We argue this exploration was driven first by increasing substrate use. Later ecological transitions were aided by ancestrally intermediate phenotypes and focused evolution towards specialized ecomorphologies.

### Phylogenomics of the Amphibolurinae

Dragon lizards (Agamidae) are a charismatic and diverse group of ∼600 species found across Africa, Asia, and Oceania. Our new sequence-capture dataset extends our phylogenetic understanding of the primarily Australo-Papuan subfamily Amphibolurinae (Macey et al. 2000; Melville et al. 2001, Hugall et al. 2008; Shoo et al. 2008; Melville et al. 2018; Chaplin et al. 2020; Fenker et al. 2024). Amphibolurines likely originated in Late Cretaceous Asia before dispersing into Australo-Papua (Scarpetta et al. 2026). This biogeographic pattern is consistent with other Australian squamate groups (Keogh 1998; Esquerre et al. 2020; Brennan et al. 2021), though the long amphibolurine stem lineage is unique. The subsequent rapid Miocene diversification of the group, followed by a later slowdown is consistent with theory around adaptive radiations and dispersal into new regions (Harmon et al. 2010; Schluter 2000; Simpson 1944; Yoder et al. 2010).

### Mapping Morphology to Ecology

The morphological landscape of amphibolurine dragons is dispersed and patchy, with peaks that largely correspond to recognized genera. Obvious phenotypic peaks relate to strongly arboreal (*Hypsilurus*, *Lophosaurus*) and terrestrial species (*Tympanocryptis*, some *Ctenophorus*) (Fig.1B). However, intermediate forms that use a variety of substrate types (*Amphibolurus, Gowidon, Lophognathus, Tropicagama, Pogona*) fill the morphological space. Evidence from GLMMs link these morphologically intermediate taxa to wider habitat usage and broader niches than more morphologically extreme relatives. This pattern, paired with evidence from discrete evolutionary modelling indicates specialist dragon niches evolve from generalist ancestors. This provides a pathway to unite disparate peaks of the landscape and facilitate transitions to new ecologies.

In morphological clustering, taxa with broader niche breadths are either lumped under a single broad adaptive peak (when k = 5 optima; Fig.S9) or two peaks (when k = 12 optima; Fig.S10). This is important because these intermediate morphologies appear common (∼50% of species richness), are distributed across the entire continent in all habitat types, and provide a theoretical link between more extreme morphological forms. Niche breadth modelling supports this by favoring pathways of narrowing substrate usage (loss of habitat states) through faster transition rates. Transitions also overwhelmingly occur through habitat generalism (use of multiple habitat states) rather than transitions from one specialist to another (Fig.S13). The idea of generalists as a bridge between morphotypes is not exclusively a reflection of how we generate morphospaces. In dragon lizards this process is informed by the branching pattern of the phylogeny. Specialist forms often descend from morphologically intermediate ancestors, and in some cases are nested deeply within generalist clades. For example the stout rock-dwelling *Diporiphora bennetti* and gracile hummock grass living *Diporiphora adductus* likely both evolved from a more typical ancestral amphibolurine morphology. While we identify both gains and losses in habitat usage, increasing specialisation through losses occur at more than six times the rate of gains. Additionally, transitions in habitat usage typically follow an ordered gain/loss of a single state, traversing through polymorphic (multi-habitat states). A notable exception are the specialist rock-dwelling *Ctenophorus* which have independently evolved twice from terrestrial ancestors.

Linking morphology to ecology is not always perfect. Here, however, GLMMs provide some evidence that morphology is predictive of substrate usage. Along PC1 (47.9% of variance) intermediate morphologies show greatest niche breadth (Fig.5). This PC displays strong loadings for size, as well as tail length, and body width. These traits seem to partition across the arboreal and terrestrial axes. However, the linear relationship of PC4 (4.4%) with breadth highlights an interesting interaction with PC1. Species with proportionally shorter limbs and longer heads (PC4) show a greater propensity for niche breadth irrespective of their size (Fig.5). In contrast, species may also exhibit many-to-one mapping where varied morphologies enable the same ecology (Wainwright et al. 2005; Moen 2019). For example, taxa such as *Lophosaurus*, *Chelosania*, and *Diporiphora superba* are all obligately arboreal and highly specialized dragons, however, these taxa map to either two or three discrete adaptive zones (Fig.3) and use arboreal spaces differently. In this situation, convergence on an arboreal lifestyle is met with only incomplete morphological similarity (Grossnickle et al. 2024). This may be due to a number of reasons including phylogenetic inertia resisting selection (Blomberg & Garland 2002). This idea is relevant when considering the ecology of the ancestral amphibolurine. Our analyses suggest this ancestor was morphologically similar to the arboreal *Lophosaurus* and *Hypsilurus*, but also to *Physignathus* and *Intellagama* — the most ecologically diverse species in the group. We flag this to note the imperfect mapping of morphology to ecology and uncertainty in estimating ancestral states.

### The Evolving Adaptive Landscape

The transition from wet closed-canopy rainforests in Asia and New Guinea towards new and different environments in Australia presents a prime case for ecological opportunity (Collar et al. 2010; Gray et al. 2019a). Among the newly available habitats, dragons encountered sparse open forests, grasslands, sandy and stony deserts, and rocky escarpments, which are all but absent across Malesia. By expanding their geographical range, dragon lizards also extended the adaptive landscape to enable new forms that took advantage of these habitats (Yoder et al. 2010). This extension provided opportunity for new peaks located in distant regions of morphospace. To traverse that space however, required deterministic trends in trait evolution that suggest selection on individual or suites of traits (Figs.3,4). In dragon lizards, this may have been aided by an ancestrally intermediate morphology (Fig.2) that changed rapidly and dramatically, resulting in a multivariate shape change dozens of times greater than expected (Fig.4 highlight 5). The resulting form likely had shorter limbs and tail, a wider body, and shorter snout as it spent less time in the trees and more time on the ground (Fig.5). From generalist forms amphibolurines were able to further explore morphological extremes. This includes repeated transitions towards small-bodied terrestrial forms (*Ctenophorus aurita*, *Rankinia*, *Tympanocryptis*), and moves back towards elongate arboreal forms (*Diporiphora superba*, *Chlamydosaurus*) (Fig.5). Along with this increasing distance from the morphological center, niche breadth narrowed creating increasingly specialized species (Fig.5). In this way, generalists negate the need to cross wide fitness valleys in the adaptive landscape. Instead, species can explore across ridges or broad fitness peaks to reach distant morphospaces. This provides more evidence that transitions across the landscape from specialist to specialist are uncommon and likely require an intermediate form (Fig.2) (Dennis et al. 2011).

**Figure 2.**
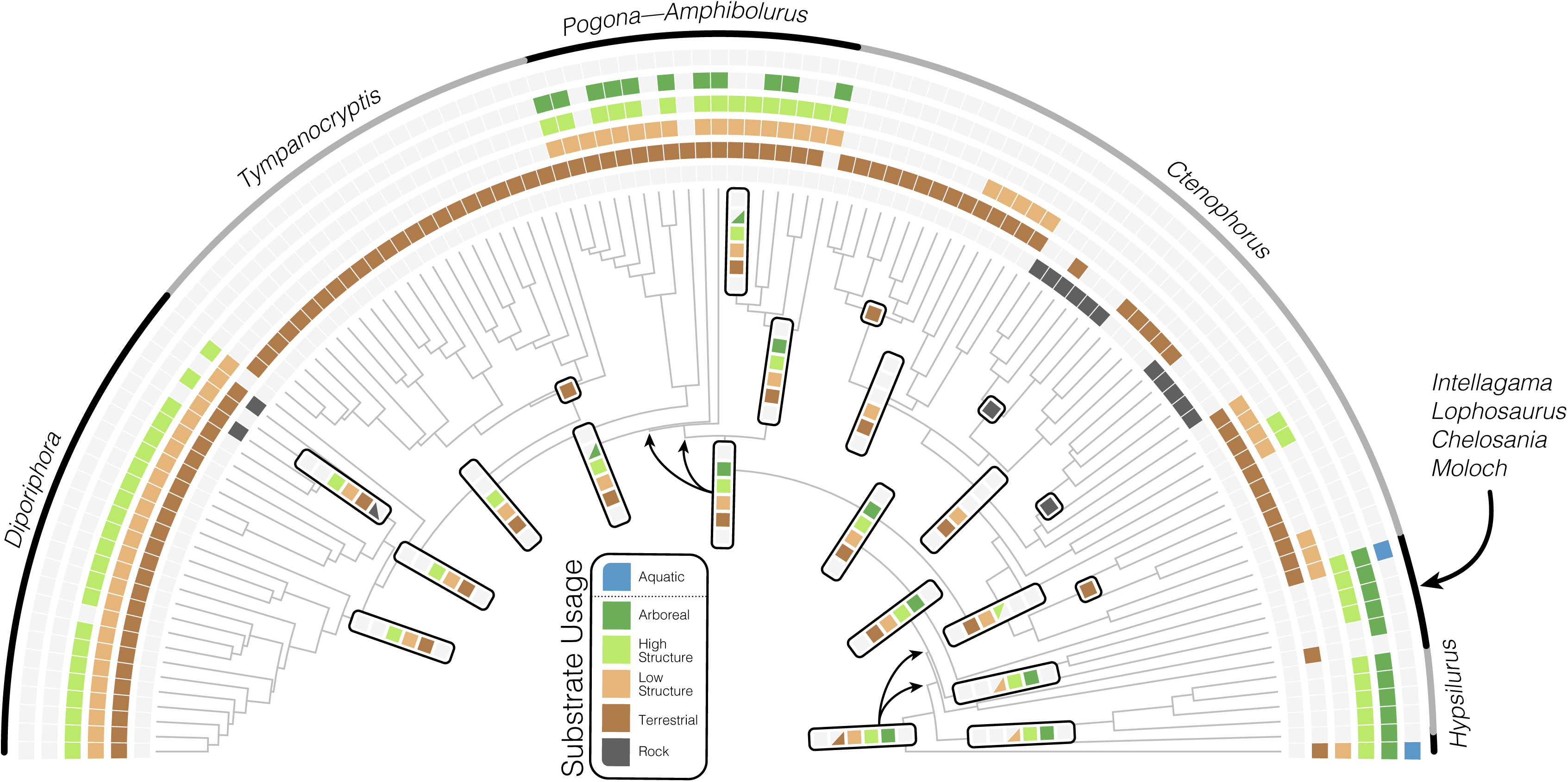
The evolution of habitat usage favors transitions from niche generalists (high niche breadth) towards specialists (low niche breadth). Six rings plotted outside the tree indicate substrate usage for each species, with colored squares indicating usage and grey squares indicating absence. Tabs at internal nodes indicate substrate usage for ancestral taxa modelled under two-rate ordered polymorphic models and estimated using a model averaging approach. Half squares (triangles) reflect state uncertainty in reconstructions (see Fig.S16). Single colored squares highlight transitions to highly specialised forms. An ancestral amphibolurine likely expanded its substrate usage to spend more time on the ground after arriving in Australia.

**Figure 3.**
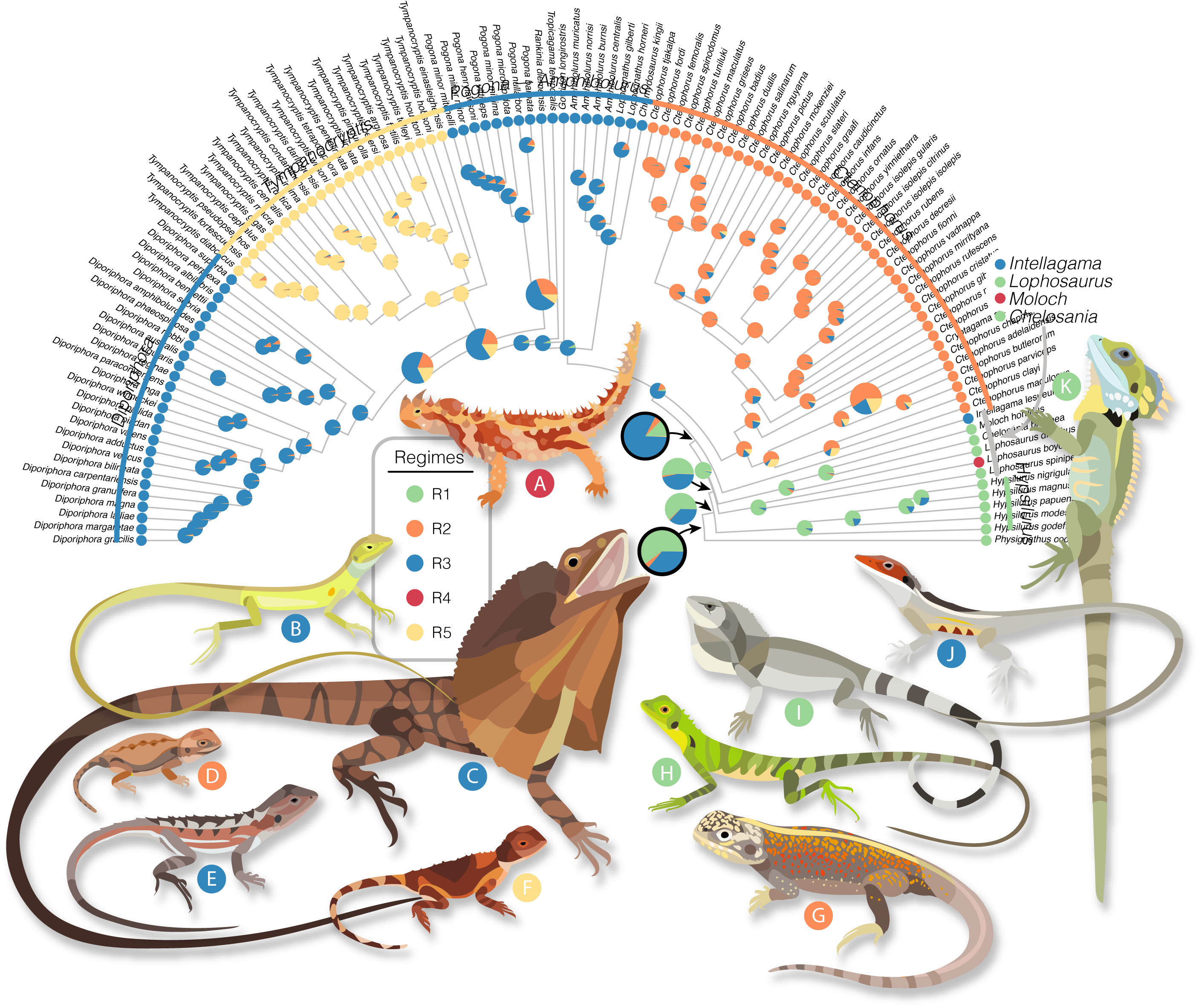
The adaptive morphological zones of amphibolurine dragons highlight a diversity of forms. Colored regimes were identified under a multi-optima OU model in *PhyloEM* and applied to ancestral taxa using *randomForests*. The tree shows the preferred five regime model estimated from all 19 size corrected morphological traits and likelihoods for ancestors as estimated by *randomForest*. Two regimes (blue, orange) map to obligately terrestrial groups. The red regime highlights primarily arboreal species and the morphologically similar *Physignathus*. Yellow indicates a diverse clade of primarily generalist species. The most recent common ancestor (MRCA) of amphibolurines is estimated as a *Hypsilurus/Lophosaurus-*like arboreal dragon and the common ancestor of core Australian taxa (excluding *Moloch, Chelosania*, and *Lophosaurus*) is estimated as a semiarboreal generalist lizard (both indicated with black outlined pie). Illustrated species are (A) *Moloch horridus,* (B) *Diporiphora superba,* (C) *Chlamydosaurus kingii,* (D) *Ctenophorus aurita,* (E) *Rankinia diemensis,* (F) *Tympanocryptis pseudopsephos,* (G) *Ctenophorus reticulatus,* (H) *Hypsilurus modestus,* (I) *Chelosania brunnea,* (J) *Gowidon longirostris,* (K) *Lophosaurus boydii*.

**Figure 4.**
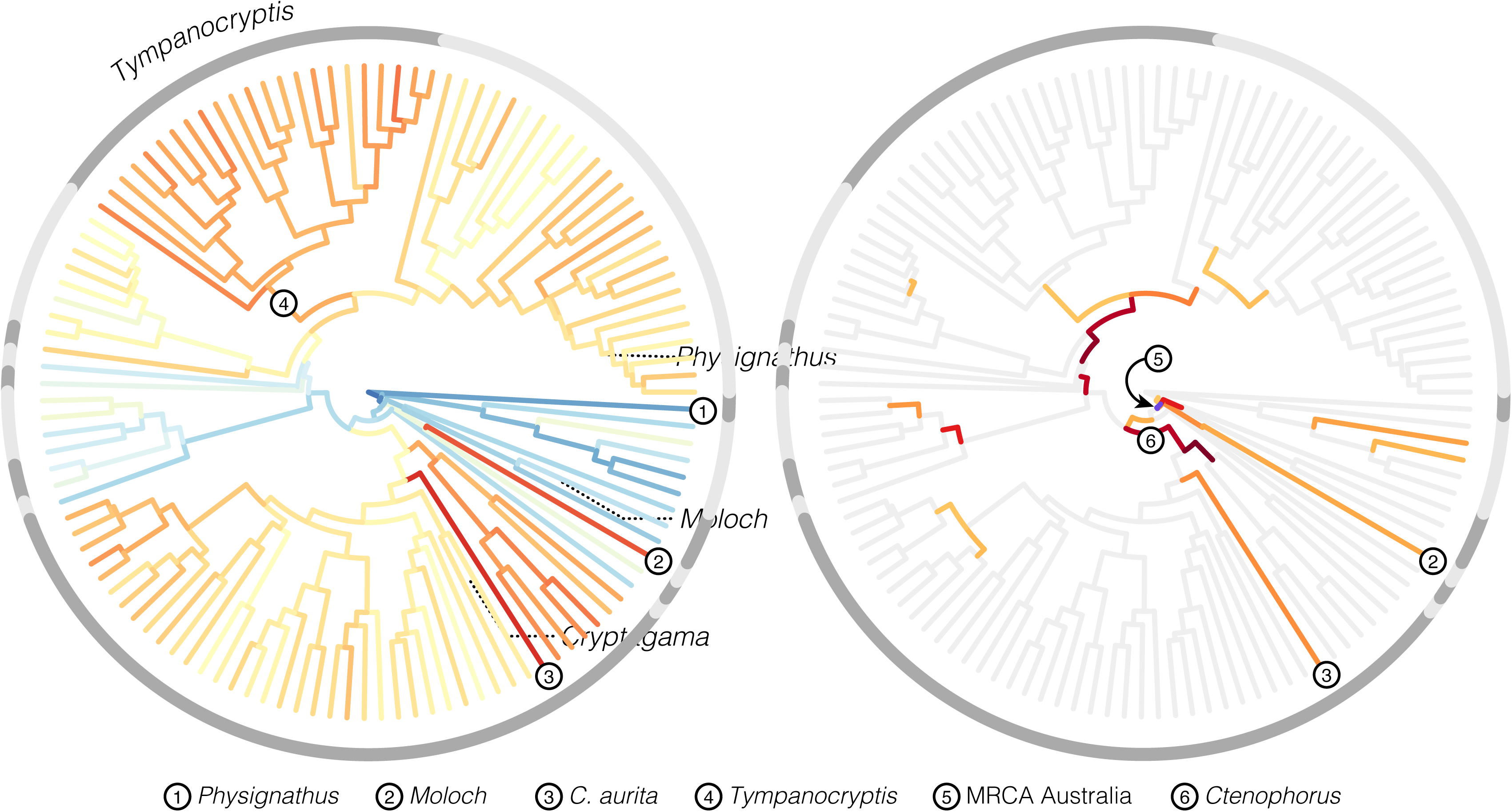
Major shifts in morphologies occur as directional evolutionary changes that are greater than expected. Left, morphological innovation across the dragon tree, with warm colors indicating greater multivariate Euclidean distance from the amphibolurine most recent common ancestor, and cooler colors indicating greater similarity. Grey shading in outer circles delineates genera. Right, the position and strength of directional changes that exceed multivariate shape change under the null evolutionary model. Light grey branches follow expectations under the null Brownian model, and colored branches indicate greater than expected change. Circle number 5 indicates the branch leading to the Australian radiation of dragon lizards and denotes a 26-fold increase over the expected amount of morphological change. This very short branch accounts for a shift in the evolution of amphibolurine lizards away from a more arboreal ancestor towards a more generalist morphology.

**Figure 5.**
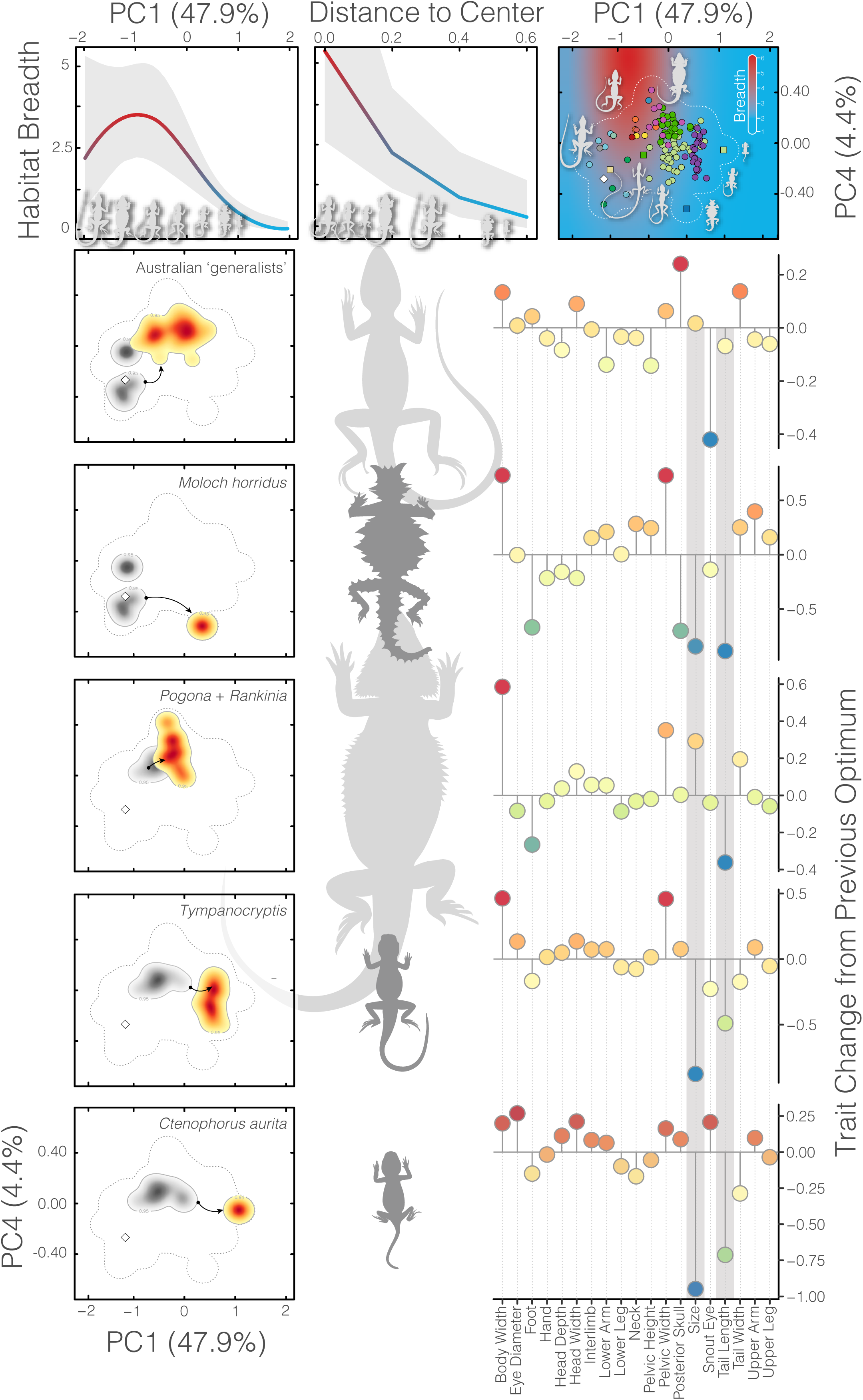
Niche breadth decreases as morphologies become more extreme through changes in suites of traits. Plots in the top row show results of GLMMs. (Left) niche breadth is greatest at intermediate PC1 scores. (Center) niche breadth decreases with increasing distance from the morphological center. (Right) visualization of predicted breadth values (background colors) plotted against PCs 1 and 4. Overlaid species points are colored by genus following Figure 1. Below, plots in the left column visualize the shift from the immediately ancestral optimum (grey shading) to a new optimum (warm shading) represented in two dimensional space. Lizards in the center column illustrate the morphological extremes of new optima. Plots at right indicate how individual traits changed in evolving from the previous optima to current. Grey bars highlight the two most variable traits: size (a composite of the geometric mean of all trait measures) and tail length.

In our investigation across the dragon lizard body plan, we identify varied modes of evolution among and within traits. The evolution of dragon morphology involved both a gradual diffusion process away from stable peaks and infrequent shifts in the evolutionary rate or trait mean. Interestingly, to explain the observed morphological landscape, fewer than 10% of branches need multivariate trait change that exceeds Brownian expectations. This is visible as discrete clumps across the morphospace. These high density areas indicate slow evolutionary rates or constrained evolution (i.e. Ornstein Uhlenbeck), as has been previously identified in *Diporiphora* (Smith et al. 2010). Stable peaks populated by many closely-related species highlight diffusion as the most frequent pathway for change among species. In comparison, rare evolutionary shifts are concentrated around changes among adaptive zones. These shifts are nested and temporally staggered resulting in a dynamic adaptive landscape that does not reflect the archetypical tempo or mode of an adaptive radiation. This complex mosaic of evolutionary modes helps to reconcile Simpson’s idea of quantum evolutionary change (few, extreme changes) with Darwin’s ideas of gradualism (frequent, small changes) (Simpson 1944; Charlesworth et al. 1982).

The morphological evolution of dragon lizards has been a heterogeneous process, varying temporally, phylogenetically, and across traits. Broadly, dragons have gone through periods of early (30-25 my) and late expansion (5-0 my) in the morphological hypervolume, bookending a period of little innovation during much of the Miocene (Fig.S8). Admittedly, this pattern lacks information provided by extinct species. However, if our expectation that ancestrally intermediate morphologies yield young derived morphologies is true, then we would not expect extinction to considerably affect our conclusions about the accumulation of morphological volume. This does establish further questions about how biased extinction of morphologically extreme species may drive the pattern that we see. Looking across all species, we identify that the primary axes for variation are through change in absolute size and tail length (Fig.S7) and that these have a significant correlation with substrate usage. This is immediately apparent when comparing large tree dragons (*Hypsilurus magnus*: SVL >200 mm, tail length >600 mm) to pebble-mimicking dragons (*Ctenophorus aurita*: SVL <40 mm, tail length <35 mm; *Tympanocryptis cephalus* group: SVL <60 mm, tail length <90 mm). Elaboration in size and tail length beyond other traits is a pattern seen in other squamate clades (Brennan et al. 2024b; Meiri 2010), but dramatic changes are not limited to these axes. Instead, amphibolurines show exceptional variation in many traits and greater than expected diversity relative to other iguanians (Gray et al. 2019a; Gray et al. 2019b). Most trait trends appear associated with habitat type (Thompson & Withers 2005; Collar et al. 2010), e.g. wide bodies in terrestrial groups, but others are harder to quantify and likely associated with defense or communication, like the frill of *Chlamydosaurus*. Harder yet to explain is the novelty of the enigmatic thorny devil, *Moloch*. They likely pre-date the Australian deserts in which they wander (Pepper & Keogh, 2021). They exclusively eat ants. And they have undergone such a profound evolutionary modification that they are nearly unrecognizable as agamids. In expanding into this new adaptive zone, the thorny devil has stretched the morphological space and become profoundly specialized. Trends towards increasing specialization in agamids show how the adaptive landscape itself can change and grow, and how species with broad niches can act as bridges to stretch borders and alter the landscape topography in response to new opportunities.

## Acknowledgements

This project represents an output of the Australian Amphibian and Reptile Genomics initiative (AusARG), with data generation generously funded by BioPlatforms Australia and salary support through an Australian Research Council Discovery Grant (DP210100820 to JSK). We appreciate the provision of computing and data resources provided by the Australian BioCommons Leadership Share (ABLeS) program and Seqera Tower service. These programs are co-funded by Bioplatforms Australia (enabled by NCRIS), the National Computational Infrastructure and Pawsey Supercomputing Centre. Special thanks to Sophie Mazard and Ziad al Bkhetan. We thank our many Australian partner institutions (Australian Museum, Museum and Art Gallery of the Northern Territory, Queensland Museum, Museums Victoria, South Australian Museum, Western Australian Museum) and their associated curators and collections managers, who made this work possible through generous tissue loans and collections access. Sincere thanks to Stephen Zozaya for insight into species ecology and habitat use. Lastly we thank two anonymous reviewers and Senior Editor Akira Mori for valuable comments on this work. We declare no conflicts of interest.

