## Supplement for "Generalists link peaks in the shifting adaptive landscape of Australia’s dragon lizards"

#### Document S1. Figures S1–S14 and Tables S1–S4

**Supplementary Figures** **17**

**Software** **39**

**Supplementary Tables** **40**

**Supplemental References** **52**

### Supplementary Information

#### Molecular Taxon Sampling

We assembled a sequence-capture dataset across 309 Amphibolurinae representing 120 of 136 currently recognized species, with a focus on Australian taxa (115 of 117 spp.). This sampling covers all 16 genera and many recognized subspecies (Table S1). We included outgroup representatives from all agamid subfamilies and used tuatara (Sphenodon punctatus) and chicken (Gallus gallus) to root the tree (368 total samples).

The following Amphibolurinae taxa were unsampled in our molecular efforts:
*Diporiphora convergens** STORR, 1974
*Hypsilurus binotatus* (MEYER, 1874)
*Hypsilurus bruijnii* (PETERS & DORIA, 1878)
*Hypsilurus capreolatus* KRAUS & MYERS, 2012
*Hypsilurus geelvinkianus* (PETERS & DORIA, 1878)
*Hypsilurus godeffroyi* (PETERS, 1867)
*Hypsilurus hikidanus* MANTHEY & DENZER, 2006
*Hypsilurus longi* (MACLEAY, 1877)
*Hypsilurus macrolepis* PETERS, 1872
*Hypsilurus modestus* (MEYER, 1874)
*Hypsilurus nigrigularis* (MEYER, 1874)
*Hypsilurus ornatus* MANTHEY & DENZER, 2006
*Hypsilurus papuensis* (MACLEAY, 1877)
*Hypsilurus schultzewestrumi* (URBAN, 1999)
*Hypsilurus tenuicephalus* MANTHEY & DENZER, 2006
*Tympanocryptis uniformis** MITCHELL, 1948

*These species were described from a single specimen and have not been confirmed via genetic data (Melville & Wilson, 2019).

#### Molecular Data

Sample collection and laboratory preparation, including extraction, shearing, size selection, library generation, and hybridization capture were completed as part of the Australian Amphibian and Reptile Genomics initiative (AusARG) under the Phylogenomics Working Group following the protocol outlined in Tiatragul et al. (2023). Raw sequence data for newly sampled taxa are available from the BioPlatforms Australia Data Portal. Assembled target sequences organized by sample (pseudo-reference genomes; PRGs) are available from https://github.com/iangbrennan/Amphibolurinae.

We used the *pipesnake* v1.2 workflow (Brennan et al. 2024) https://github.com/AusARG/pipesnake to handle, process, assemble, and align our molecular data, estimate gene trees and a species tree. All software employed for individual tasks are indicated in the Supplementary file software_versions.yml. *pipesnake* is a highly reproducible and flexible workflow that provides consistent results.

In brief, the pipeline concatenates individual read files, then removes duplicate reads via BBMAP (Bushnell 2014), trims adapters and barcodes with TRIMMOMATIC (Bolger et al. 2014), identifies read pairs with PEAR (Zhang et al. 2014), removes off-target reads with BBMAP, assembles reads to contigs with SPAdes (Prjibelski et al. 2020), maps contigs to targets with BLAT (Kent 2002) and extracts the best hit for each target, carries out preliminary alignments with MAFFT (Katoh & Standley 2013), before refining the alignments with GBLOCKS (Talavera & Castresana 2007), estimates locus trees with IQ-TREE2 (Minh et al. 2020), and a species tree with weighted ASTRAL hybrid (Zhang & Mirarab 2022).

#### Additional Sampling

To take advantage of all available morphological and phylogenetic information we incorporated three additional New Guinean *Hypsilurus* species into our dated tree used for trait evolution analyses. *Hypsilurus godeffroyi*, *H. papuensis*, and *H. nigrigularis* were added to our MCMCTree output using bind.tip in phytools (Revell 2024), based on the topology and ages estimated from a recent phylogenomic investigation of Squamata (Title & Singhal et al. 2024).

#### Phylogenomics of the Amphibolurinae

Dragon lizards (Agamidae) are a charismatic and diverse group of ~600 species found across Africa, Asia, and Oceania. They are diurnal and often brightly colored and territorial, making them visible and familiar. The Amphibolurinae subfamily extends out of Asia into Australia and New Guinea, where they reach ecomorphological extremes including burrowing species that live along the salt-crusted edges of desert lake-beds and semiaquatic species that patrol the eastern watercourses. While species such as the bearded dragons and frilled-neck lizard are well-known, phylogenetic understanding of the group has been limited in genetic scope and focused within genera (Macey et al. 2000; Melville et al. 2001, Hugall et al. 2008; Shoo et al. 2008; Melville et al. 2018; Chaplin et al. 2020; Fenker et al. 2024). Our new sequence-capture dataset provides a well-supported estimate of the relationships among all 17 amphibolurine genera and more than 88% of described species.

Relationships among some (Streicher et al. 2016; Burbrink et al. 2020; Title et al. 2024) or all (Welt & Raxworthy 2022; Scarpetta et al. 2025) agamid subfamilies have been estimated from different phylogenomic datasets, but have rarely been discussed. Our topology differs from that of Welt & Raxworthy (2022) and Scarpetta et al. (2025) in the placement of the Leiolepidinae and Uromastycinae at the base of the tree, and the stepwise (rather than sister) relationship between the Hydrosaurinae and Amphibolurinae. The topology we present has biogeographic implications, suggesting an Asian and not African origin for the group. This is further supported by fossil evidence of the Priscagamidae which were stem-acrodonts of east Asia, and the ~99 myo *Protodraco* from Myanmar (Wagner et al. 2021) which is potentially the oldest known acrodont. The debate over an Asian or African origin of agamids is mirrored in other Australian squamate groups such as pythons (Esquerre et al. 2020), monitor lizards (Brennan et al. 2021), and elapid snakes (Keogh 1998). One possible explanation for differences in the subfamily topology is the expanded molecular dataset presented here. Reanalyzing a UCE-only dataset however, returns a topology identical to our SqCL estimate, suggesting other factors (e.g. alignment parameters, substitution models) are likely causing the differences in topology.

While the Amphibolurinae as a group date back to the Cretaceous, their living diversity is less than half that age. The split between the sole Asian species Physignathus and the remaining Australo-Papuan diversity occurred around the Eocene/Oligocene transition, and was closely followed by rapid diversification over the next ten million years (Fig.1). This is consistent with paleontological evidence from Australia which suggests fossils assignable to Intellagama are relatively common in the early-to-middle Miocene deposits of the Riversleigh World Heritage Area (Covacevich et al. 1990). The rapid divergence of Australian species from their forest-dwelling relatives in New Guinea resulted in a series of very rapid branching events which can prove challenging for phylogenetic methods. These deep and rapid splits occur over just a few hundred thousand years and highlight a period of dramatic ecomorphological change. Following this period of radiation, the diversification of the group is far slower, more homogeneous, and dictated by three major groups: Ctenophorus (41 spp.), Diporiphora (28 spp.), and Tympanocryptis (26 spp.). This pattern of speciation slow-down is consistent with theory around adaptive radiations and dispersal into new regions (Harmon et al. 2010; Schluter 2000; Simpson 1944; Yoder et al. 2010).

#### Divergence Dating

Divergence Dating To estimate divergence times among taxa we applied a series of fossil and secondary calibrations in MCMCTree (Rannala & Yang 2007) as outlined in the Supplement (Table S2). We started by trimming our ASTRAL tree down to a single representative for each ingroup species or subspecies for input. We reduced our molecular data to exonic markers in the AHE loci, then estimated raw genetic distances from alignments to use as a proxy for evolutionary rate. We removed the fastest 5% and slowest 5% of loci to avoid issues with extreme rate heterogeneity, removed third codon positions from all loci, then partitioned the remaining loci into three partitions using AMAS (Borowiec 2016).

For each partition we ran MCMCTree with usedata = 3 to get the approximate likelihoods and branch lengths using baseml (dos Reis & Yang 2011), then concatenated the out.BV files. We then ran four replicate MCMCTree analyses on the gradient and Hessian (in.BV file; usedata = 2), each for 20k burn-in generations before collecting 20k samples at a sampling frequency of 100 generations (2,020,000 total generations). We compared mcmc files for stationarity and convergence (ESS of all parameters > 200), combined them using logCombiner, and used this combined mcmc file to summarize divergence times on our tree (print = -1 in .ctl file). To validate our priors we ran an additional analysis (usedata = 0) to run explicitly from the prior calibrations and determine our effective priors for comparison against our posterior age estimates (Fig.S1).

##### Node Priors and Agamids in the Fossil Record

Acrodontan lizards are known from the fossil record of Europe, Australia, and across Asia, with arguably the oldest described lineage originating in Myanmar (Wagner et al. 2021). Unfortunately, the oldest fossil taxa (*Protodraco*, *Gueragama*) are of uncertain phylogenetic placement and remaining fossils (*Uromastyx europaeus*, *Barbaturex*, *Tinosaurus*) are considerably younger than recent estimates of subfamily splits in agamids (Title et al. 2024; Burbrink et al. 2020). Agamid fossils are known from Australia (Covacevich 1990; Ramm

2025), however most are comparatively young and unlikely to be of value as calibrations. The exceptions are mid-Miocene agamid fossils from Riversleigh World Heritage Area assigned to *Sulcatidens* and '*Physignathus*'. The latter were ascribed to *Physignathus* when this genus applied to both *P. cocincinus* and *I. lesueurii*, and leaves the current placement of these fossils uncertain. They are clearly Australian in origin, and while similar to *Intellagama*, they also share affinities with *Physignathus*, *Hypsilurus*, and *Chelosania* (M. Hutchinson, *pers. comm.*). These fossils were found across a number of Riversleigh sites, all of which have been dated to 16-18 myo by Woodhead et al. (2016): Camel Sputum (17.8), Wayne's Wok (17.8), Inabayance (17.8), Upper (17.8), and RSO (16.6). To use this fossil we applied it as a minimum on the crown of Australian Amphibolurinae, with a soft upper bound of 50 mya informed by estimates of the Amphibolurinae—Draconinae/Agaminae split as estimated by Burbrink et al. (2020) and Title et al. (2024). We applied a secondary calibration on the divergence between Amphibolurinae and Agaminae (Phrynocephalus) as a uniform prior with soft bounds (50-80 mya) following Burbrink et al. 2020 and Title et al. 2024, and another uniform prior with soft bounds (238-255 mya) on the crown of Lepidosauria representing the fossil *Sophineta*, arguably the oldest stem squamate. To assess the appropriateness of this calibration strategy we downsampled our species tree to a single representative of each genus and ran preliminary analyses in MCMCTree.

#### Optima Model Fitting

To explore the reliability of our results concerning the number and species assignment of optima we fit a number of alternative schemes. In addition to the base scheme of PCs 1-6 (>90% variance) and K=0-15, we also fit PCs 1-9 (>95% variance) and all 19 raw trait axes. To assess a more conservative view we also reduced the number of possible shifts, therefore limiting the number of optima to 5 (K=0-4). The results are highly consistent regardless of input data type (PCs vs. raw traits) or trait number (6, 9, 19), in identifying a core set of 4 shifts that are recovered in every scenario. Those correspond to (1) a baseline shift at the base of the Australian radiation of Amphibolurinae, and three shifts that correspond to primarily terrestrial taxa in (2) the edge leading to *Moloch*, (3) *Ctenophorus* + *Cryptagama*, and (4) *Tympanocryptis.* Additional shifts identified by favored (PCs 1-6) and less conservative (K>4) analyses identify optima that largely correspond to genera, including (5) *Lophosaurus*, (6) *Pogona* + *Rankinia*, (7) *Diporiphora*, and (8) the large-bodied clade of *Ctenophorus.* The remaining shifts identify outlying individual species that show unique morphologies, (9) *Diporiphora superba*, (10) *Chlamydosaurus kingii*, and (11) *Ctenophorus aurita.* The final regime belongs to the taxa at the base of the tree comprising *Intellagama* and *Hypsilurus.*

##### Incomplete Convergence

*Diporiphora superba* is an extremely gracile species with a long tail, small head, and long slender limbs that it uses to move through shrubs. Lophosaurus species similarly have long tails and limbs, but have large heads and deep, laterally compressed bodies and generally perch vertically on tree trunks. Chelosania also show a narrowing of the body but have short sturdy limbs and a relatively short tail for a species that is almost never found outside of the canopy where they often use horizontal branches. This may be due to a number of reasons including phylogenetic inertia resisting selection (Blomberg & Garland 2002). This idea is relevant when considering the ecology of the ancestral amphibolurine. Our analyses suggest this ancestor was likely morphologically similar to the arboreal *Lophosaurus* and *Hypsilurus*, but also to Physignathus and Intellagama—two species that are considered semiaquatic (Fig.S12). These last two species are similar enough that they were considered congeneric for over 150 years and are often called "water dragons". Despite their preferred habitat, they fit most morphological descriptors of an arboreal dragon. We flag this to note the imperfect mapping of morphology to ecology and uncertainty in estimating ancestral states.

##### Calculating Morphological Distances

To estimate multivariate morphological distances we calculated euclidean distance between pairs of taxa. For Figure 4 this required identifying transitions among optima and measuring distances between ancestral taxa or between an ancestor and extant taxon (see function node.to.node in script Morphological_Analysis/07_funspace.R).

#### Modelling Niche Breadth

To characterize species niches we discretized microhabitat and structure usage into six categories. We then fit species to these classifications following the ecology and biology of the species in Melville & Wilson (2019). We scored a species as 1 for a microhabitat trait if that feature is regularly used and 0 if not used. While most species will use elevated structures for brief perching and basking (even *Tympanocryptis* species will climb grass tussocks or stones) we only scored species as 1 if the microhabitat is a regular part of their daily activity. We differentiate low and high structure, trees, and rock outcrops due to different physical requirements resulting from shear forces. For example, narrow branches and twigs in low and high structure allow for ample grasping and are often oriented horizontally. In contrast, trees and rock outcrops provide broad or vertical surfaces that may require different morphologies (longer limbs, claws) to access them. Evidence for this partitioning among substrate types has been documented for Australian dragons (Stuart-Fox & Owens, 2004; Collar et al. 2010).

- **Aquatic**: water courses including streams and rivers
- **Rock**: exposed rock surfaces and outcrops with little-to-no vegetation
- **Ground**: all terrestrial surfaces (sand, stony plain, compact soil, cracking clays, grassland) that are not elevated
- **Low Structure**: fallen logs, stones, grasses <30 cm high
- **High Structure**: taller vegetation such as shrubs >30 cm high
- **Tree**: tall vegetation with broad vertical surfaces

This substrate usage scheme allowed us to categorize species into three character coding schemes. The first (**Polymorphic**) used the coding scheme as-is and modelled the acquisition of loss of individual character states using polymorphism-aware Markov model. In the second (**Niche Breadth**), we summed substrate scores to develop a measure of species niche breadth. Low scores indicate limited breadth and are indicative of substrate specialists, while high scores indicate generalist tendencies. For ease of reference we call this the Niche Breadth (NB) coding. While this coding system distinguishes species with wide and narrow breadths, it lumps all specialist taxa into a single coding, which obscures the ability to test transitions among specialist states. To address this, we created a second coding scheme (**Generalist-Specialist**) which classified taxa with only one or two substrates as specialists in their primary state, and all taxa with three or more substrates as generalists. This allowed us to specifically test if transitions are more likely between generalist and specialist states than they are between specialist states.

##### Polymorphic State Models:

To model the evolution of substrate usage, we used the *fitpolyMk* model of *phytools*. We started by dropping the 'aquatic' character state from the substrate character coding because it is rare (2 species) and increases the model state space significantly without providing valuable insight into habitat evolution. The discrete substrate states were translated into numeric characters (rock–>0, terrestrial–>1, low structure–>2, high structure–>3, tree–>4). We then established a set of models as follows:

- **eqr.unorder**: “equal rate unordred”. A single rate unordered polymorphism model, allowing transitions between character states through polymorphic states (e.g. to transition from 0 to 4 occurs via 0–>0+2–>2).
- **eqr.ordered**: “equal rate ordered”. A single rate ordered polymorphism model, allowing transitions between adjacent character states only (e.g. to transition from 0 to 2 occurs via 0–>0+1–>0+1+2–>1+2–>2).
- **two.unorder**: “two rate unordered”. A model where the acquisition of a state and the loss of a state occur at different rates. This is an unordered “transient” model in *phytools* terminology.
- **two.ordered**: “two rate ordered”. A two rate ordered model where the acquisition of a state and the loss of a state occur at different rates. This is an ordered “transient” model in *phytools* terminology.
- **eqr.unorder.jump**: “equal rate unordered jump”. A single rate unordered model that allows transitions directly between different character states without forcing them to go through a polymorphic state (e.g. a transition from 0 to 2 can occur via 0<–>2). We refer to these transitions directly between states as “jumps”.
- **two.unorder.jump**: “two rate unordered jump”. A two rate unordered model that allows transitions directly between different character states without forcing them to go through a polymorphic state (as in eqr.unorder.jump), but acquisition and loss of a state occur at different rates.
- **eqr.order.jump**: “equal rate ordered jump”. A single rate ordered model that allows transitions directly between adjacent character states (e.g. 0<–>1).
- **two.order.jump**: “two rate ordered jump”. A two rate ordered model that allows transitions directly between adjacent character states (e.g. 0<–>1), but acquisition and loss of a state occur at different rates.
- **rock.order.jump**: “two rate rock ordered jump”. A two rate ordered model as above, but the only “jump” transition allowed is between ground and rock substrates (as observed).

The ordered models above work on the assumption that state transitions occur between adjacent states. In our case, this suggests that substrate type is ordered vertically (ground <–> low structure <–> high structure . . . ). This is founded on the observation that species frequently use more than one adjacent structure state, but very rarely use discontinuous states (e.g. ground, tree).

**Polymorphic Model Results** Model fitting results for the Polymorphic models. Ancestral state estimations were generated using weighted model averages across all fitted models.

```
##                     log(L) d.f.      AIC       weight     delta
## object           -171.0163    1 344.0326 2.384198e-11 47.699665
## eqr.unorder.jump -170.5469    1 343.0937 3.812646e-11 46.760744
## two.unorder      -159.4092    2 322.8184 9.637439e-07 26.485399
## eqr.ordered      -156.6576    1 315.3153 4.104329e-05 18.982285
## two.unorder.jump -154.6862    2 313.3725 1.084217e-04 17.039483
## eqr.order.jump   -154.7588    1 311.5176 2.740899e-04 15.184627
```

```
## two.ordered      -147.1767    2 298.3533 1.979163e-01  2.020341
## two.order.jump   -146.9109    2 297.8218 2.581675e-01  1.488812
## rock.order.jump  -146.1665    2 296.3330 5.434917e-01  0.000000
```

**Niche Breadth Models:**

We generalized niche breadth by summing trait usage. Values of 1-6 indicate highly specialized (low values) to absolute generalist species (high values).

- **ER**: equal rates of transition among all 6 breadth states
- **ARD**: separate transition rates among all breadth states.
- **STP**: stepwise transition in the acquisition or loss of breadth states, with independent rates for each transition.
- **STP-ER**: stepwise transition in the acquisition or loss of breadth states, with equal rates for each transition.
- **NJ-ER**: a loose stepwise model where transitions are allowed only between n+2 states (e.g. 1→2, 1→3). All transition rates are equal.
- **NJ**: a loose stepwise model where transitions are allowed only between n+2 states (e.g. 1→2, 1→3). Transitions between increasing specialization (e.g. 3→2, 2→1) and increasing generalism (e.g. 2→3, 1→2) are allowed separate rates.
- **INC**: a two rate model where increasing specialization (e.g. 3→2, 2→1) and increasing generalism (e.g. 2→3, 1→2) have different rates.
- **SPC**: a one-directional model where transitions may only happen from generalism towards specialization and not in reverse.

**Niche Breadth Model Results** Model fitting results for the Niche Breadth models. Ancestral state estimations were generated using weighted model averages across all fitted models.

```
##               log(L) d.f.      AIC       weight     delta
## fit.ARD   -80.73266   20 201.4653 8.453649e-05 17.467514
## fit.STP   -92.11783    8 200.2357 1.563377e-04 16.237844
## fit.STPER -97.78807    1 197.5761 5.909735e-04 13.578338
## fit.INC   -94.55858    2 193.1172 5.493203e-03  9.119347
## fit.ER    -93.52645    1 189.0529 4.191502e-02  5.055082
## fit.NJ    -91.09626    2 186.1925 1.751821e-01  2.194718
## fit.NJER  -91.73389    1 185.4678 2.516897e-01  1.469976
## fit.SPC   -90.99891    1 183.9978 5.248881e-01  0.000000
```

Discrete trait models of niche breadth favored models in which transitions occur through neighboring (n±1) or near-neighboring (n±2) states (Fig.2, Fig.S13). AIC indicated support for a two-rate (NJ) model with a faster transition rate towards decreasing niche breadth (increasing specialization) (AICw = 0.54), however a likelihood ratio test is unable to reject ($p = 0.39$) a simpler single-rate (NJ-ER) model (AICw = 0.29). Model fitting of transitions among specialist (1 substrate) and generalist (3+ substrates) habitat usage indicated support for a model (GENSPC) where transition rates are faster between generalist and specialist states, and slower—but not absent—among specialist states (AICw = 0.50). However, a likelihood ratio test is unable to reject (p = 0.10) a simpler single-rate model (ER; AICw = 0.36) in which transitions between the generalist and specialist states occur at equal rates.

**Generalist-Specialist Models:**

To investigate transitions among specialist and generalist states we classified taxa as either specialist (one or two substrates used) or generalist (>=3 substrates used) category.

- **ER**: equal rates of transition among all 6 breadth states
- **ARD**: separate transition rates among all breadth states
- **SYM**: symmetric transition rates between each pair of states
- **GENSPC**: two rate model where transitions between specialist states (rate 1) and between specialist and generalist states (rate 2 ) differ.

**Generalist Specialist Model Results** Model fitting results for the Generalist-Specialist models. Ancestral state estimations were generated using weighted model averages across all fitted models.

```
##                log(L) d.f.      AIC     weight     delta
## fit.GEN     -69.84953    1 141.6991 0.00103916 12.778534
## fit.ARD     -56.88436   12 137.7687 0.00741559  8.848191
## fit.ER      -65.29414    1 132.5883 0.09886964  3.667756
## fit.GENSPC2 -64.26816    2 132.5363 0.10147246  3.615785
## fit.SYM     -59.73774    6 131.4755 0.17246634  2.554956
## fit.ROCK    -63.46027    1 128.9205 0.61873681  0.000000
```

#### Model Bias and Sensitivity

To assess the reliability of our modelling inferences we designed a simulation exercise to (1) determine the relative accuracy of our machine-learning approach in assigning ancestral regimes based on morphological data, and (2) investigate the potential for bias resulting from the interaction of our tree, data, and models. /Scripts/Morphological_Analysis/10_randomForests_Sensitivity.R.

To establish reasonable starting parameters for simulations we began by fitting a multi-OU model to each morphological trait along our SIMMAP tree of mapped amphibolurine regimes (k=5) using mvMORPH. These model fits provided empirical estimates of theta for each optimum, and single alpha and sigma values. We then fit an equal-rates model to our empirical regime data using phytools and used the transition rate to simulate the evolution of a discrete character with five states under an equal-rates model along our amphibolurine tree. To simulate the 19 continuous morphological traits we used the newly generated SIMMAP tree and the empirical multi-OU parameter values. We then used RandomForest to classify the simulated morphological data and regimes. Our next step was to estimate trait values at all internal nodes of the tree for each trait, and predict the regime of each internal node based on the RandomForest classifications. We carried out this procedure 500 times.

Our last step was to summarize the machine-learning classification outputs and compare them against the known simulated data. To identify the relative accuracy of the method, we compared the known character state of an internal node against the highest estimated classification (if >=0.5; else a node was listed as 'equivocal'). We collated this information across every node for all 500 simulations (59,000 comparisons), accuracy was calculated as # correct / 59,000. We also assessed accuracy on a node-by-node basis with the expectation that older nodes may show reduced accuracy relative to shallow nodes (Fig.S14). To determine if there was a bias towards reconstructing some regimes, we tabulated the number of times a regime was reconstructed at nodes of particular interest (MRCA Amphibolurinae, node120; MRCA of Australian clade, node131). We compared the frequencies across groups, and to the known simulated state frequencies. Results did not depart from expectations, suggesting limited bias in inferred ancestral states.

### GLM Model Fitting

To link habitat usage with morphology we used a series of Generalized Linear Mixed Models (GLMMs). The expectation is that habitat usage (as the sum of habitat types used) will increase with decreasing distance to the center of the morphospace, and that intermediate morphologies will exhibit greatest habitat usage breadth. The idea is that morphological centrism supports niche generalism. We investigate this by measuring the distance between each taxon and the morphological center, and by looking at the distribution of PC scores. We undertake this exercise with two sets of models outlined below. We tested several model families (gaussian, poisson, zero-truncated poisson, truncated Conway-Maxwell poisson) and assessed family/model fit and adequacy via three common tests (K-S, Dispersion, Outlier) in *DHARMa*. Our data best fit the assumptions of the zero-truncated poisson family model and so we applied this to both exercises. P-values are the result of Wald-Z test.

##### Breadth as a Function of Distance to Morphological Center

Model Input and Variables:

- **bdist**: [dataframe] holding the predictor and response variables with a single entry [row] per species.
- **breadth**: [column] the breadth score for a species based on summing all habitat types used.
- **mah**: [column] the mahalanobis distance of each species calculated to the geometric center of the morphospace.
- **agam.mat**: [matrix] the phylogenetic variance-covariance matrix of the time-calibrated amphibolurine tree, under the assumption of Brownian Motion.
- **propto()**: [function] provides the covariance structure of the data. Within the propto call, we provide a known distance matrix (**agam.mat**) as the variance-covariance among species. The `0 + species` part forces the model to calculate a distinct variance profile for each species, suppressing a global random intercept. This is conditioned on `g` which provides a global evaluation of the covariance matrix across all species. We follow the design of Williams et al. 2026.

Intercept-only fit

```
## glmmTMB(formula = breadth ~ 1, data = bdist, family = truncated_poisson(link = "log")
##     REML = F, ziformula = ~0, dispformula = ~1)

##              Estimate Std..Error  z.value     Pr...z..
## (Intercept) 0.606119 0.05461007 11.09903 1.268092e-28
```

Non-Phylogenetic fit

```
## glmmTMB(formula = breadth ~ mah, data = bdist, family = truncated_poisson(link = "log
##     REML = F, ziformula = ~0, dispformula = ~1)
```

```
##               Estimate Std..Error   z.value     Pr...z..
## (Intercept)  0.9771836 0.08638727 11.311661 1.148932e-29
## mah         -4.1487789 0.86611215 -4.790117 1.666837e-06
```

Phylogenetic fit

```
## glmmTMB(formula = breadth ~ mah + propto(0 + species | g, agam.mat),
##     data = bdist, family = truncated_poisson(link = "log"), REML = F,
##     ziformula = ~0, dispformula = ~1)
```

```
##              Estimate Std..Error   z.value     Pr...z..
## (Intercept)  1.618747  0.3518083  4.601220 4.200244e-06
## mah         -4.115205  1.2268068 -3.354404 7.953628e-04
```

AIC Comparison

```
##                  phy     nonphy intercept
## AIC         583.4738  657.64749  682.3656
## BIC         593.8653  664.57515  685.8294
## logLik     -288.7369 -326.82375 -340.1828
## -2*log(L)   577.4738  653.64749  680.3656
## df.resid    233.0000  234.00000  235.0000
## delta.AIC     0.0000   74.17371   98.8918
```

##### Breadth as a Function of Principal Component Scores

Model Input and Variables:

- **lps**: [dataframe] holding the predictor and response variables with a single entry [row] per species.
- **breadth**: [column] the breadth score for a species based on summing all habitat types used.
- **PC1 ... PC6**: [column] principal component scores for each species, calculated from 19 size-corrected linear morphological measurements.
- **agam.mat2**: [matrix] the phylogenetic variance-covariance matrix of the time-calibrated amphibolurine tree, under the assumption of Brownian Motion.
- **propto()**: [function] provides the covariance structure of the data. Within the propto call, we provide a known distance matrix (**agam.mat2**) as the variance-covariance among species. The `0 + species` part forces the model to calculate a distinct variance profile for each species, suppressing a global random intercept. This is conditioned on `g` which provides a global evaluation of the covariance matrix across all species. We follow the design of Williams et al. 2026.

Intercept-only fit

```
## glmmTMB(formula = breadth ~ 1, data = lps, family = truncated_poisson(link = "log"),
##     REML = F, ziformula = ~0, dispformula = ~1)
```

```
##              Estimate Std..Error  z.value     Pr...z..
## (Intercept) 0.4800403 0.08337969 5.757281 8.547979e-09
```

Non-Phylogenetic fit

```
## glmmTMB(formula = breadth ~ PC1 + I(PC1^2) + PC2 + I(PC2^2) +
##     PC3 + I(PC3^2) + PC4 + I(PC4^2) + PC5 + I(PC5^2) + PC6 +
##     I(PC6^2), data = lps, family = truncated_poisson(link = "log"),
##     REML = F, ziformula = ~0, dispformula = ~1)
```

```
##                Estimate Std..Error     z.value     Pr...z..
## (Intercept)  0.37469383  0.2084427  1.79758674 7.224252e-02
## PC1         -1.85902690  0.4013296 -4.63217005 3.618527e-06
## I(PC1^2)    -1.09265612  0.4275991 -2.55532861 1.060876e-02
## PC2         -0.01185736  0.2745108 -0.04319449 9.655465e-01
## I(PC2^2)     0.62689137  0.4414977  1.41991983 1.556310e-01
## PC3         -0.36935312  0.5009547 -0.73729842 4.609409e-01
## I(PC3^2)    -1.60916234  1.4957002 -1.07585884 2.819904e-01
## PC4          2.17071758  0.8587999  2.52761745 1.148394e-02
## I(PC4^2)    -2.88829150  2.6162875 -1.10396563 2.696080e-01
## PC5         -1.25882606  0.8843203 -1.42349553 1.545926e-01
## I(PC5^2)    -3.12981826  4.4624393 -0.70136938 4.830725e-01
## PC6         -0.36946034  0.7448716 -0.49600543 6.198906e-01
## I(PC6^2)     1.80211459  3.3137419  0.54383070 5.865580e-01
```

Phylogenetic fit

```
## glmmTMB(formula = breadth ~ PC1 + I(PC1^2) + PC2 + I(PC2^2) +
##     PC3 + I(PC3^2) + PC4 + I(PC4^2) + PC5 + I(PC5^2) + PC6 +
##     I(PC6^2) + propto(0 + species | g, agam.mat2), data = lps,
##     family = truncated_poisson(link = "log"), REML = F, ziformula = ~0,
##     dispformula = ~1)
```

```
##                Estimate Std..Error     z.value     Pr...z..
## (Intercept)  0.37469372  0.2084427  1.79758606 7.224263e-02
## PC1         -1.85902779  0.4013297 -4.63217151 3.618502e-06
## I(PC1^2)    -1.09265662  0.4275991 -2.55532949 1.060873e-02
## PC2         -0.01185772  0.2745109 -0.04319581 9.655454e-01
## I(PC2^2)     0.62689164  0.4414978  1.41992016 1.556309e-01
## PC3         -0.36935333  0.5009548 -0.73729871 4.609407e-01
## I(PC3^2)    -1.60916192  1.4957002 -1.07585861 2.819905e-01
## PC4          2.17071625  0.8587999  2.52761597 1.148399e-02
## I(PC4^2)    -2.88829029  2.6162876 -1.10396514 2.696083e-01
## PC5         -1.25882692  0.8843203 -1.42349654 1.545923e-01
## I(PC5^2)    -3.12979480  4.4624385 -0.70136424 4.830757e-01
## PC6         -0.36946047  0.7448718 -0.49600544 6.198906e-01
## I(PC6^2)     1.80208971  3.3137453  0.54382264 5.865635e-01
```

AIC Comparison

```
##                  phy    nonphy  intercept
## AIC         266.8775  264.8775  323.20422
## BIC         305.7853  301.0061  325.98334
## logLik     -119.4388 -119.4388 -160.60211
## -2*log(L)   238.8775  238.8775  321.20422
## df.resid    105.0000  106.0000  118.00000
## delta.AIC     2.0000    0.0000   58.32668
```

The AIC comparison table above indicates that the non-phylogenetic model provides a marginally better fit to the data than the model with a phylogenetic random-effect. All parameter estimates (beta, SE, Z, p-value) between the phylogenetic and non-phylogenetic models are identical, and so do not change the results. Conceptually, the equivalent fit of the two models suggests no effect of the phylogenetic covariance structure. Regardless, we retain the phylogenetic model on philosophical grounds (the non-independence of taxa).

#### Visualizing Trajectories

To help visualize evolutionary trajectories relative to the morphological center, we created a branch-by-branch summary of morphological trends. To do this we measured the angle of morphological change between each node, their direct ancestral node, and the morphological center/midpoint (see Fig.S15C). We classified trajectories with angles $<90°$ as moving towards the morphological center, and classified trajectories with angles $\geq 90°$ as moving away from the morphological center. For example, an acute angle of 5° is evidence of a strong move towards the hub of the morphospace, while an obtuse angle of 175° is evidence of a strong move away from the hub. The color and saturation of each branch indicates the direction and value of the angle of change (Fig.S15D). Early branches of the tree predominantly show movement towards the center of morphospace. To emphasize this relationship, we ran a linear model to fit angle-of-change as a function of node height. Results are presented below.

```
##
## Call:
## lm(formula = angle ~ parent.height, data = angles.df)
##
## Residuals:
##     Min      1Q  Median      3Q     Max
## -51.336 -14.983   1.545  12.691  55.036
##
## Coefficients:
##               Estimate Std. Error t value Pr(>|t|)
## (Intercept)    96.5633     2.6779  36.060  < 2e-16 ***
## parent.height  -0.8496     0.2157  -3.939 0.000108 ***
## ---
## Signif. codes:  0 '***' 0.001 '**' 0.01 '*' 0.05 '.' 0.1 ' ' 1
##
## Residual standard error: 20.11 on 234 degrees of freedom
## Multiple R-squared:  0.06218,    Adjusted R-squared:  0.05817
## F-statistic: 15.51 on 1 and 234 DF,  p-value: 0.0001081
```

### Supplementary Figures

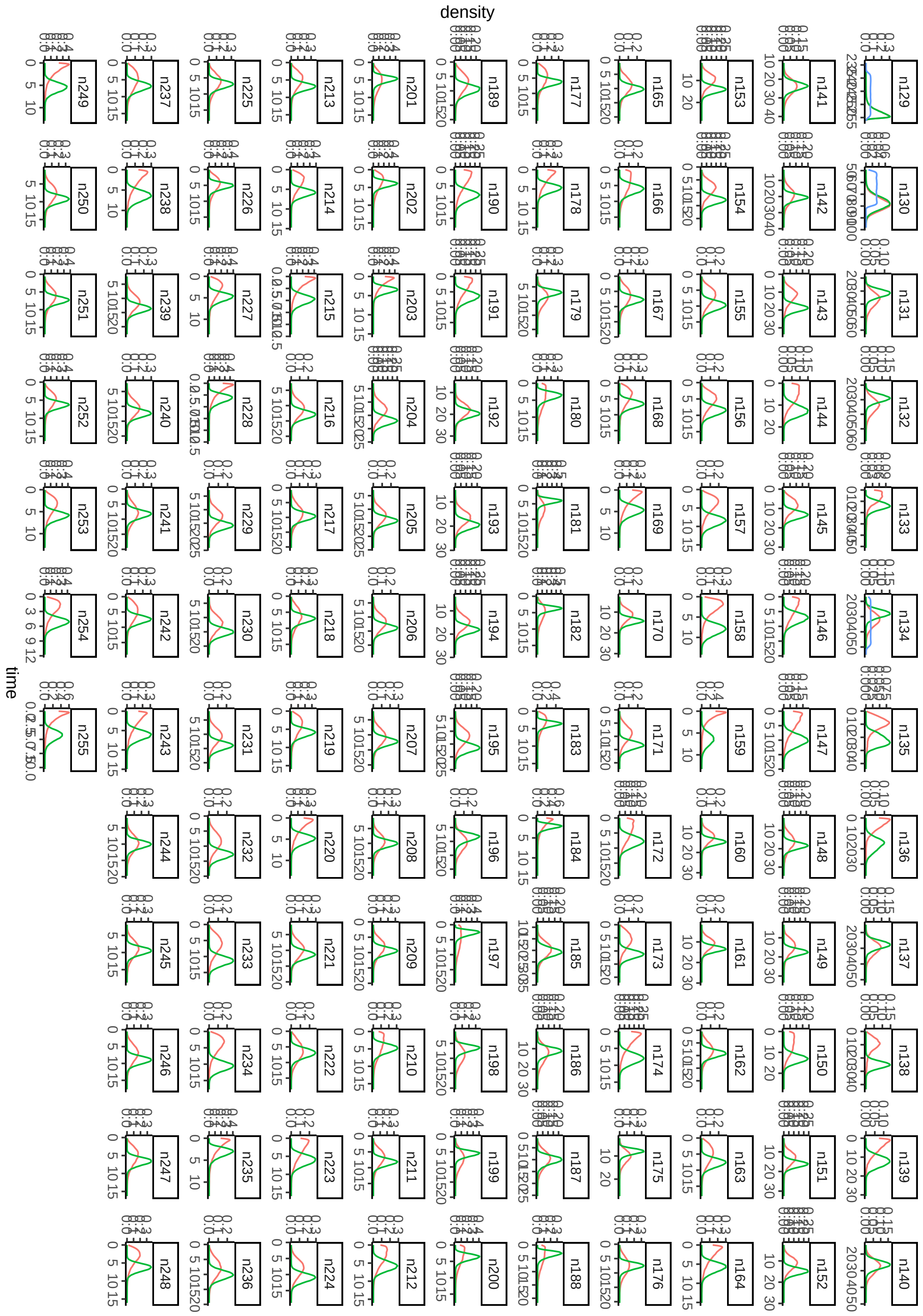

**Figure S1.** Plots of applied priors (blue), effective priors (pink), and posteriors (green) for each node in the tree indicate reasonable behavior. Node numbers correspond to the phylogeny plotted below, plotted with ape, where n129 is the root node. The effective prior (pink) shows how the interaction of multiple priors can shape the expected ages for a given node. This can be seen when comparing input (blue) and effective (pink) priors on any calibrated node (e.g. n129, n130, n134).

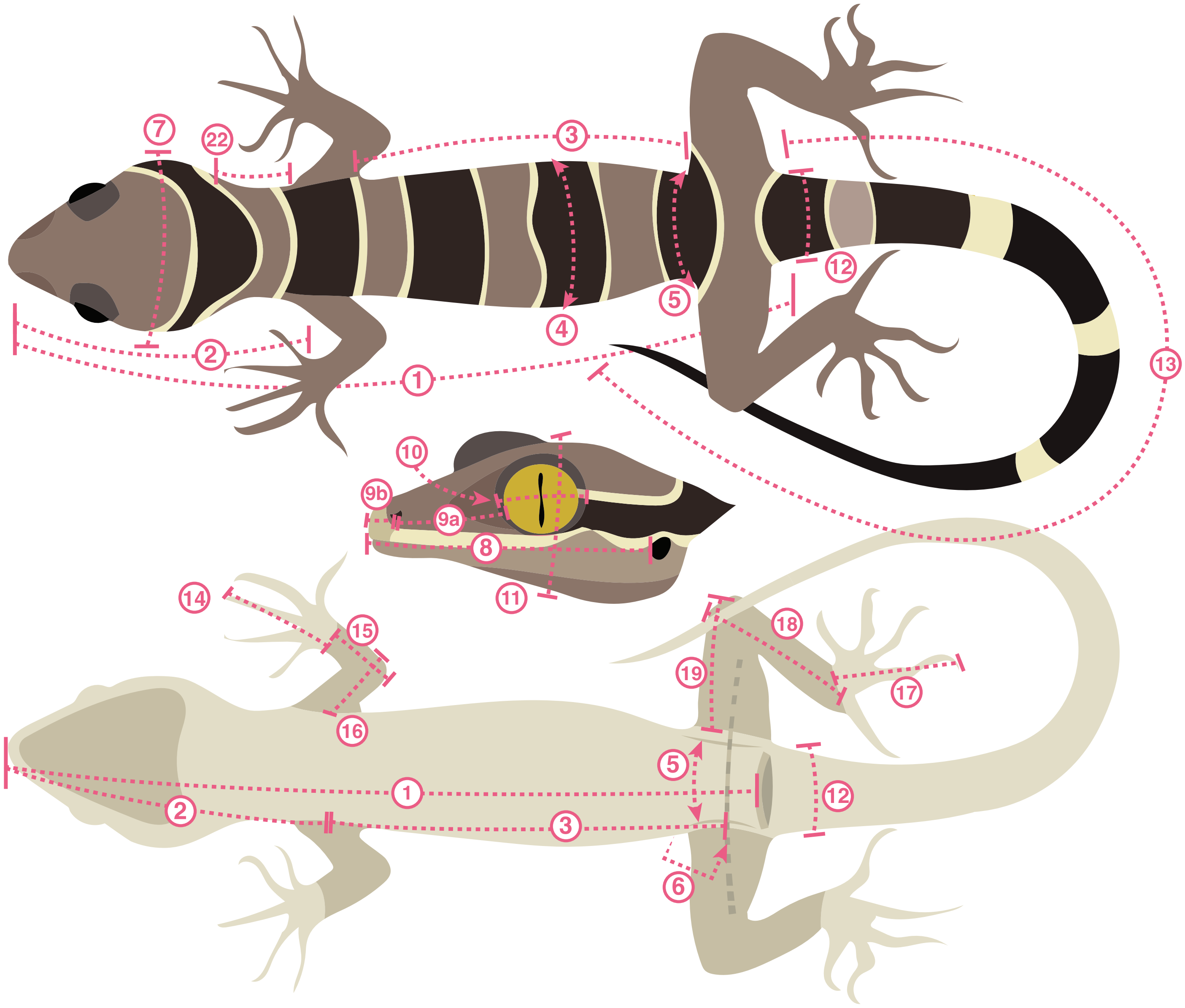

**Figure S2.** Schematic of morphological measurements taken and outlined in Table S3.

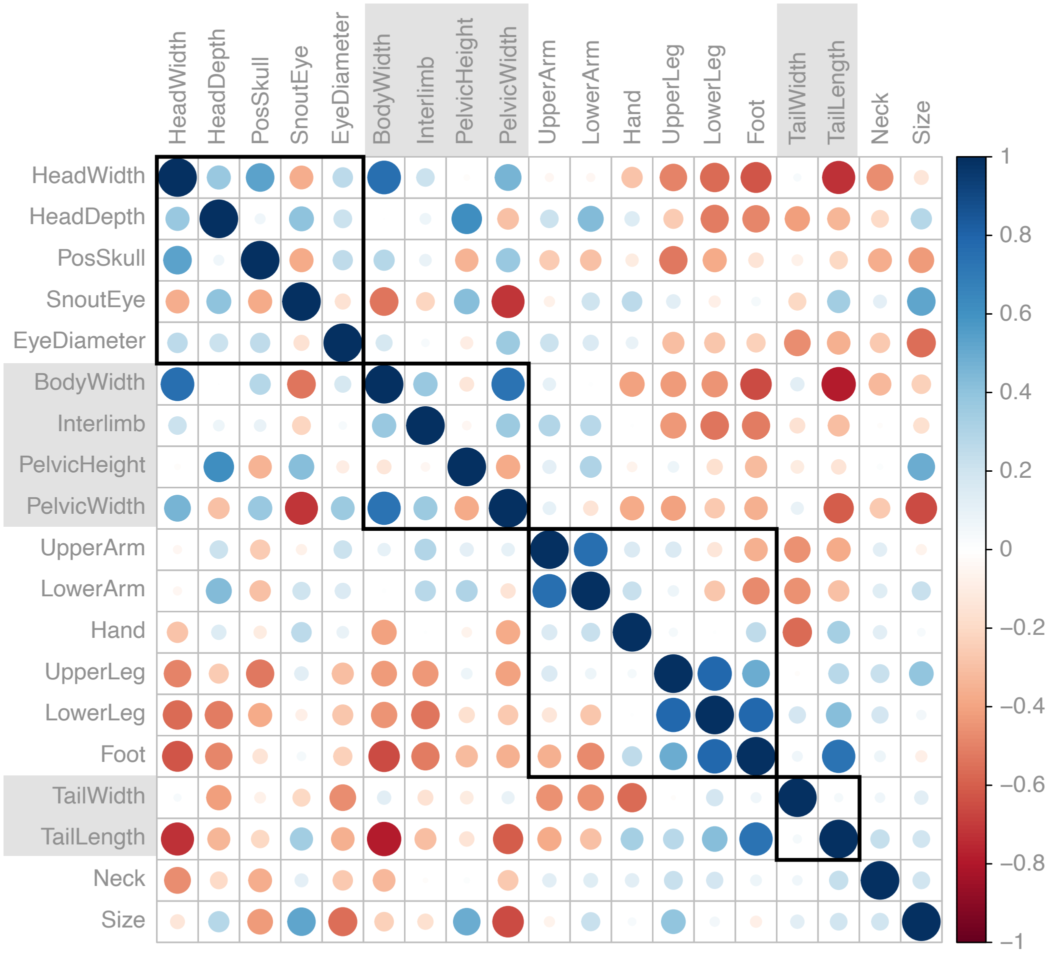

**Figure S3.** Plot of size-corrected trait correlations with traits grouped by the morphological module they correspond to (either head, body, limb, or tail). Blues indicate positive correlations, red indicate negative correlations, with saturation indicating increasing intensity.

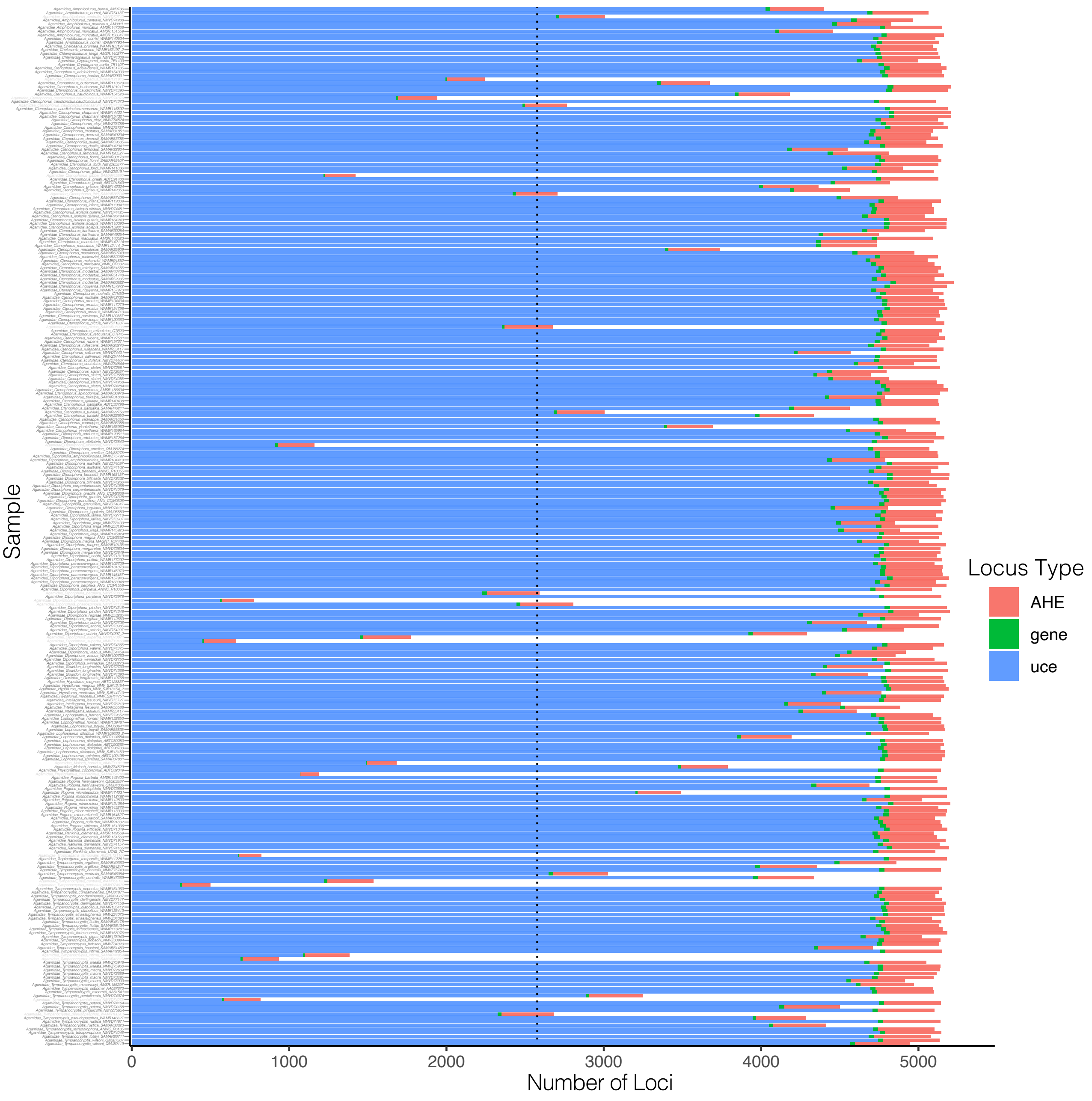

**Figure S4.** Locus type and number summarized by sample for newly sequenced samples. Dotted line indicates 50% of targeted sequences recovered. Samples in grey text at left were not included in final analyses.

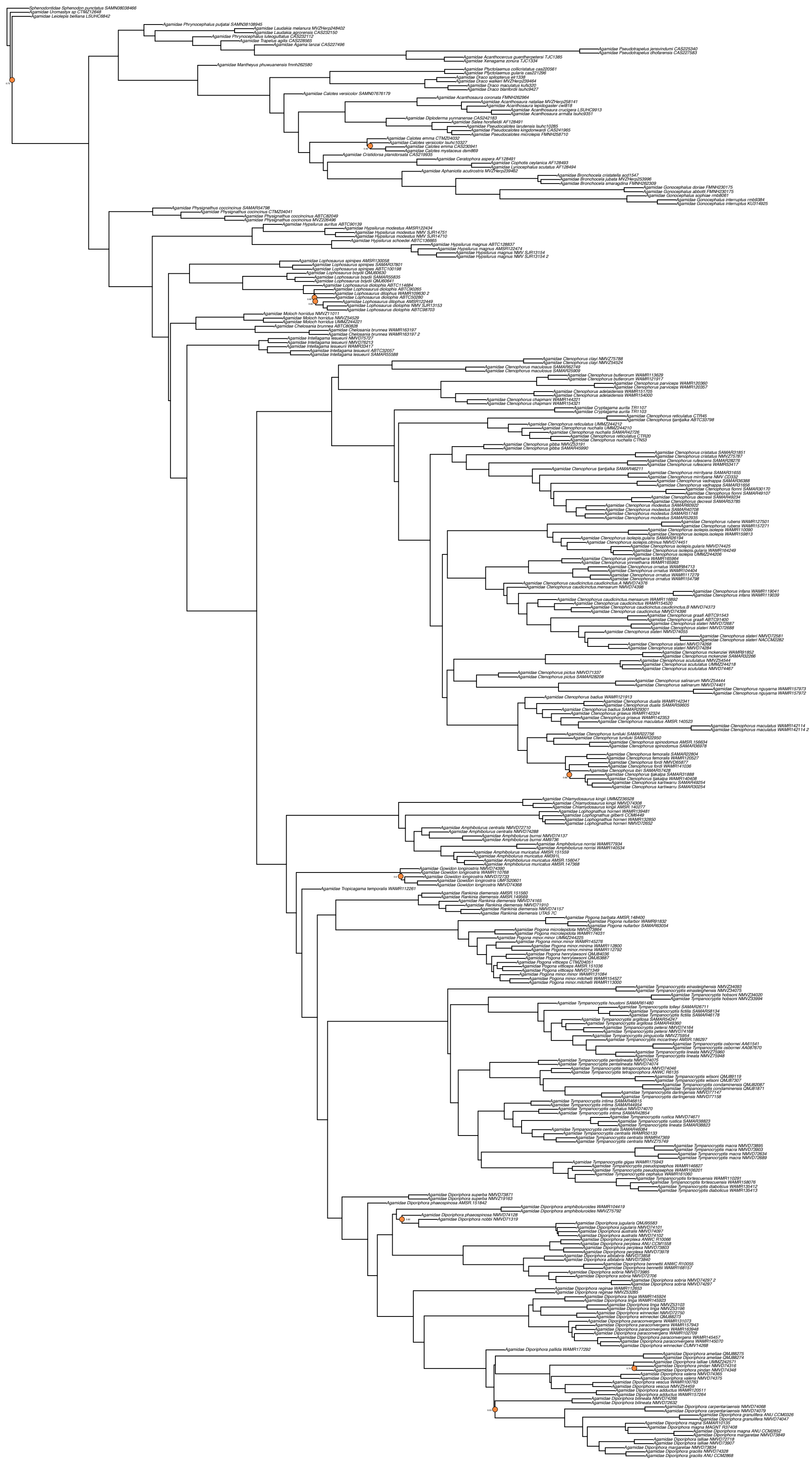

**Figure S5.** Species tree from weighted ASTRAL-hybrid including all samples sequenced and agamid outgroups. Local posterior probabilities are 1 for all nodes except those indicated by an orange circle.

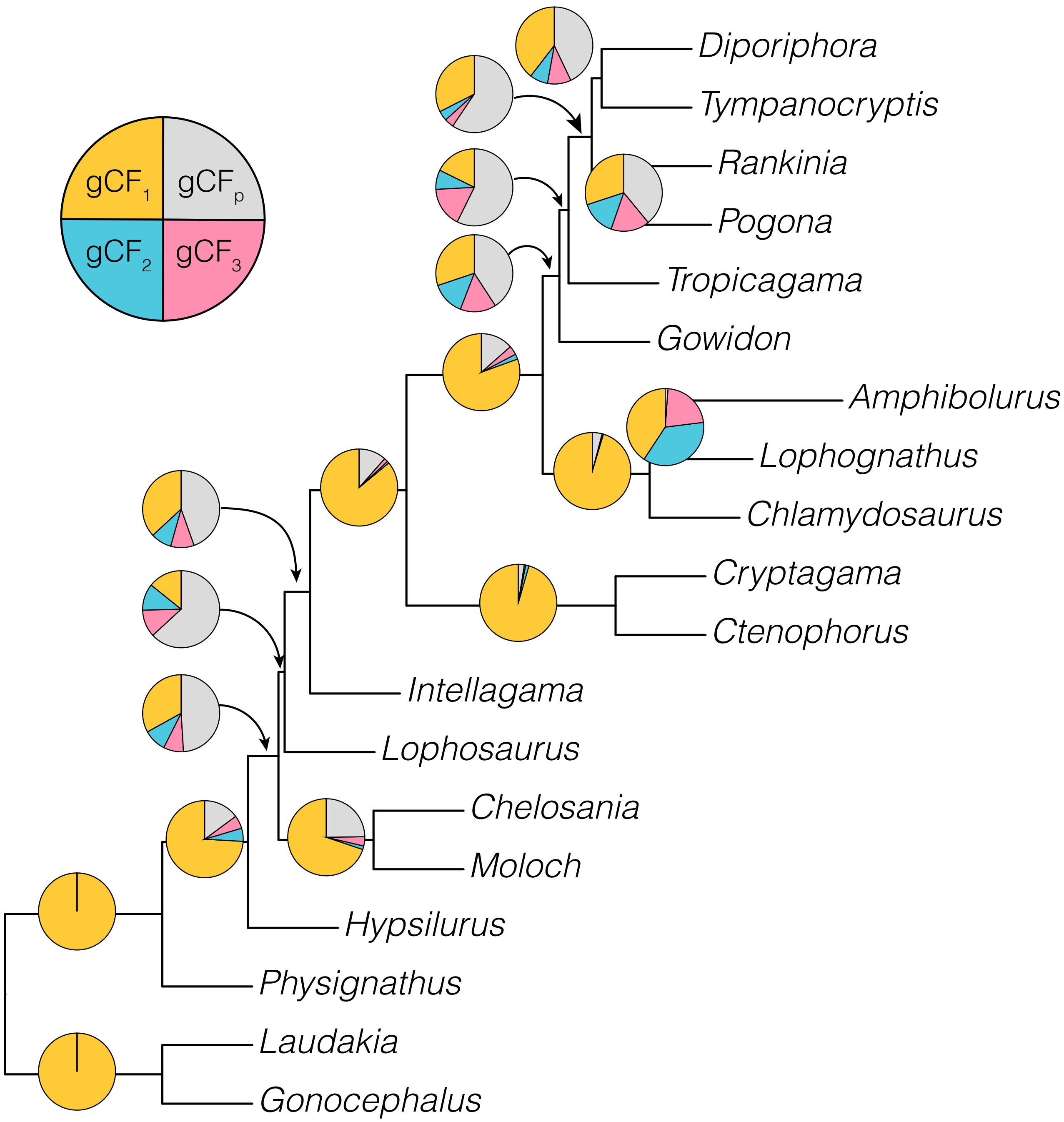

**Figure S6.** Phylogeny of the Amphibolurinae showing intergeneric relationships and support across individual loci. Gene concordance factors (gCF) show the proportion of gene trees which support a given bifurcation. Pie charts on branches indicate the percent of loci which support the presented bifurcation ($gCF_1$, orange), one of each of the two other most common resolutions ($gCF_2$, blue; $gCF_3$, pink) or all other possibilities ($gCF_p$, grey). Areas of low concordance (e.g. placement of *Lophosaurus*) indicate topological uncertainty, likely as a result of high levels of incomplete lineage sorting.

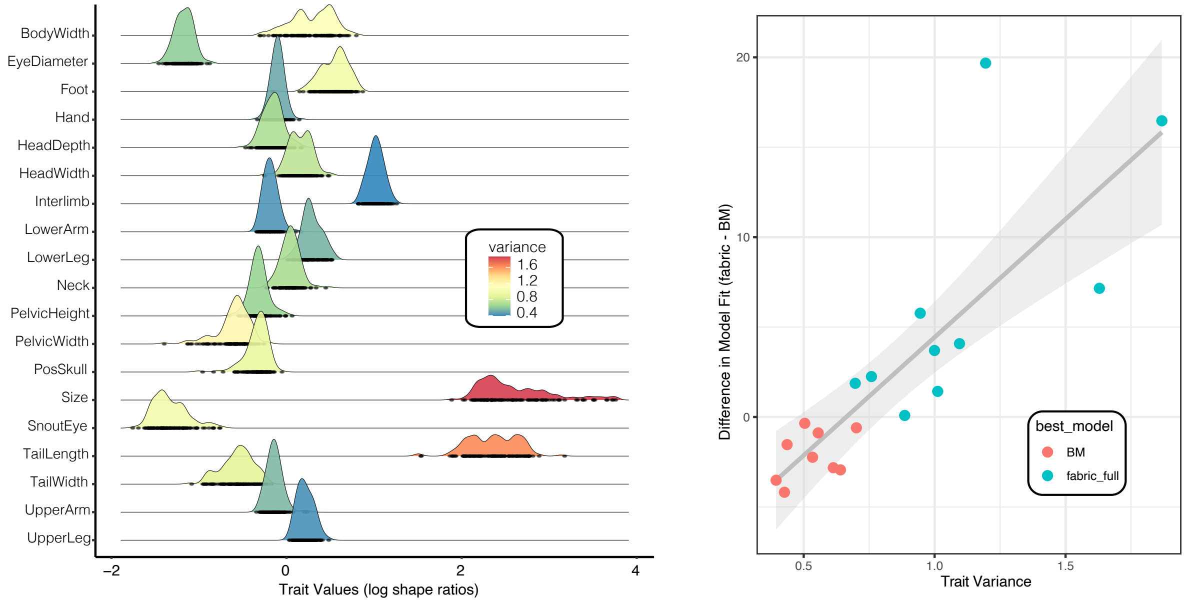

**Figure S7.** Individual traits show different levels of variance among amphibolurine species, with the greatest variances occurring in size (geometric mean) and tail length. Some traits, such as hand length and upper arm length show vanishingly small variation once corrected for absolute size. (Right) Traits with higher variances are more likely to fit the fabric model, which better partitions large changes in trait values.

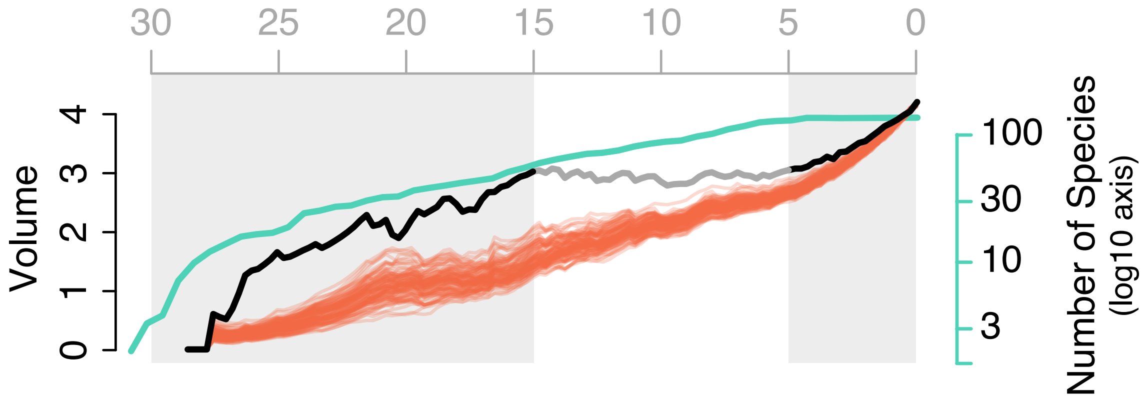

**Figure S8.** The Amphibolurinae hypervolume expands early and late in their evolution, with a period of niche packing in the Miocene. Diversification and morphological expansion are coupled in the early radiation of dragon lizards, but are largely decoupled from the Miocene onwards. Green line represents the accumulation curve for species, black line the hypervolume through time, and red lines are 500 null expectations.

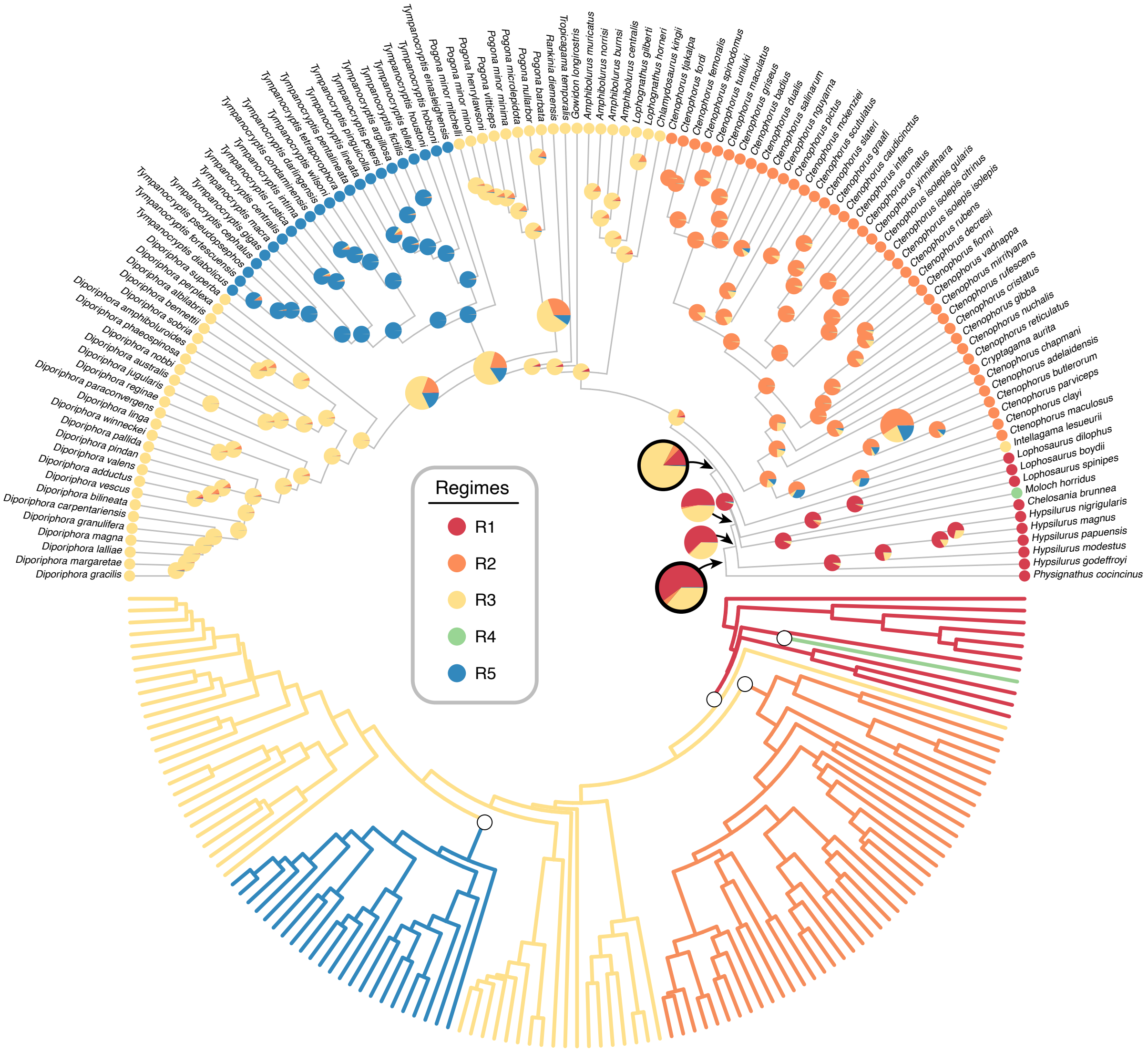

**Figure S9.** Adaptive peaks corresponding to morphological regimes identified under a multi-optima OU model in *PhyloEM* and applied to ancestral taxa using randomForests. (Bottom) Tree shows the preferred 5 regime model estimated from all 19 size corrected morphological traits, with regime shifts denoted by small white circles. (Top) Tree shows the preferred 5 regimes including likelihoods for ancestors as estimated by randomForest. The amphibolurine MRCA is estimated as a *Hypsilurus*/*Lophosaurus*-like arboreal dragon and the Australian MRCA is estimated as a semiarboreal generalist lizard (both indicated with black outline to pie).

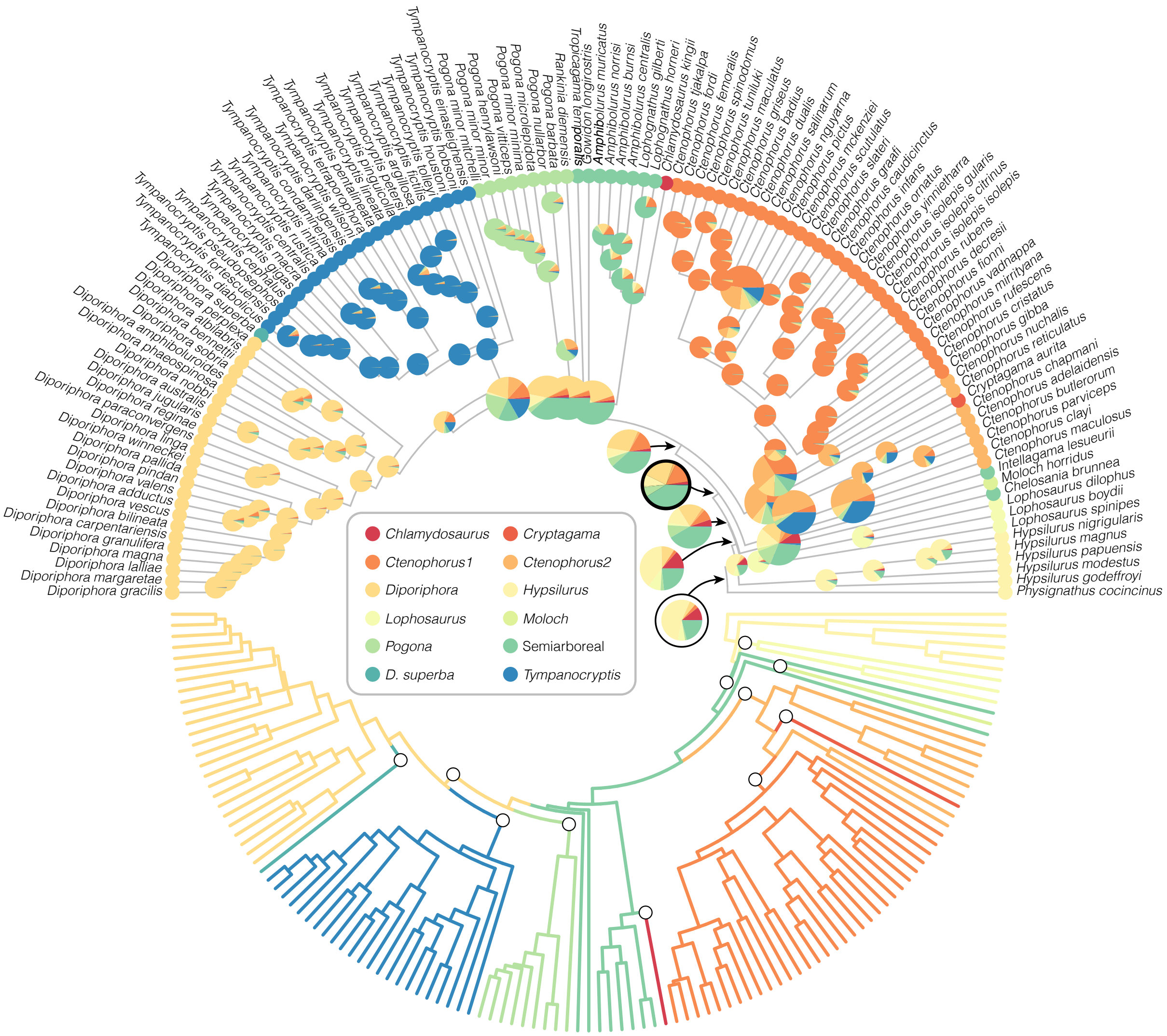

**Figure S10.** Adaptive peaks corresponding to morphological regimes identified under a multi-optima OU model in *PhyloEM* and applied to ancestral taxa using randomForests. (Bottom) Tree shows the preferred 12 regime model estimated from the first 6 PC axes of the morphological data, with regime shifts denoted by small white circles. (Top) Tree shows the preferred 12 regimes including likelihoods for ancestors as estimated by randomForest. The amphibolurine MRCA is estimated as a *Hypsilurus*-like arboreal dragon and the Australian MRCA is estimated as a semiarboreal lizard (both indicated with black outline to pie).

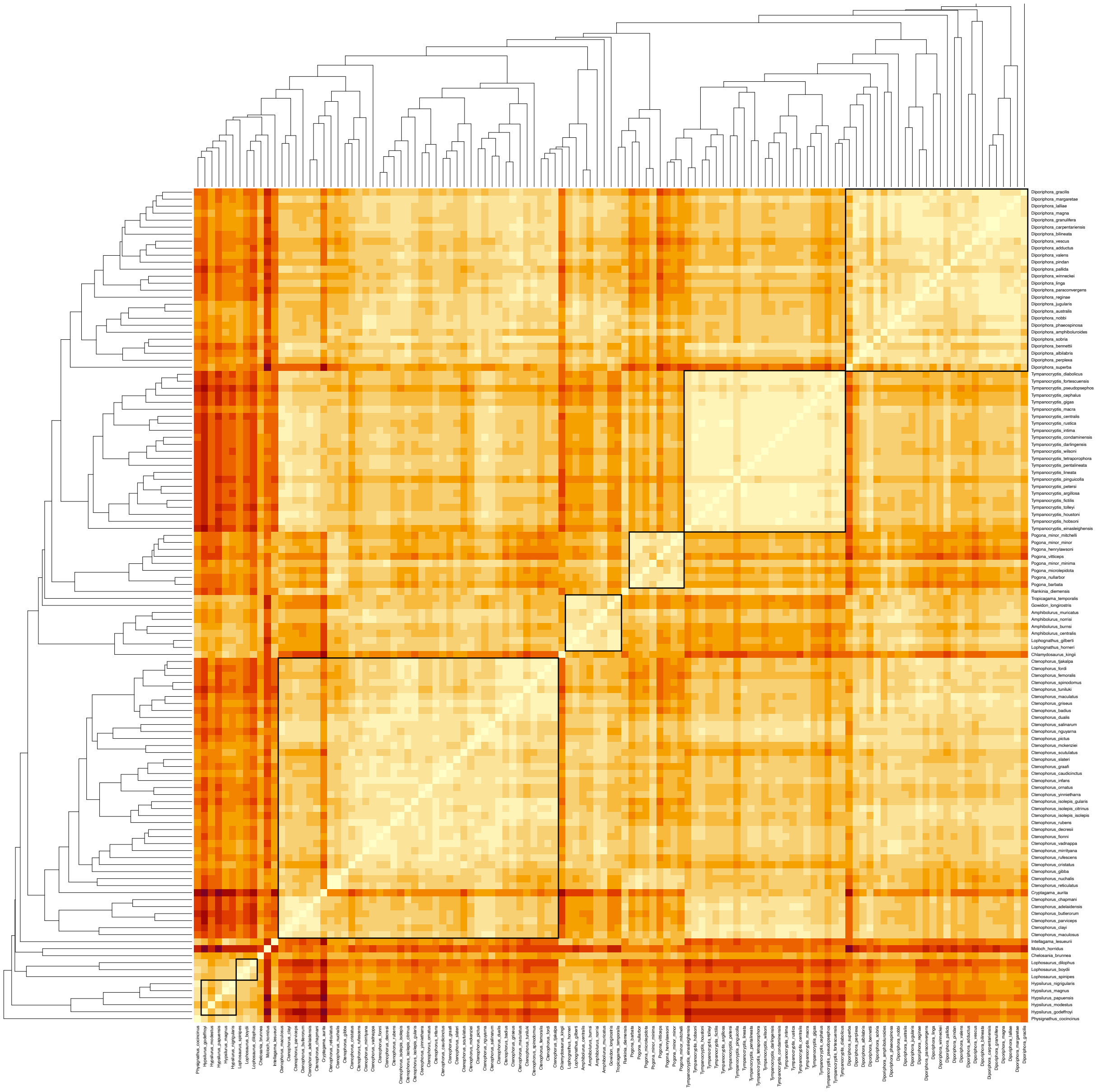

**Figure S11.** Heatmap of multivariate morphological euclidean distance among all pairs of amphibolurines. This presentation highlights the conservative evolution of *Tympanocryptis* along with the relative dissimilarity of novel species like *Chlamydosaurus* (more similar to *Lophosaurus*/*Hypsilurus*), *Ctenophorus aurita* (more similar to *Tympanocryptis*), and *Moloch.* Black boxes along the diagonal delineate genera or clades of morphologically similar taxa.

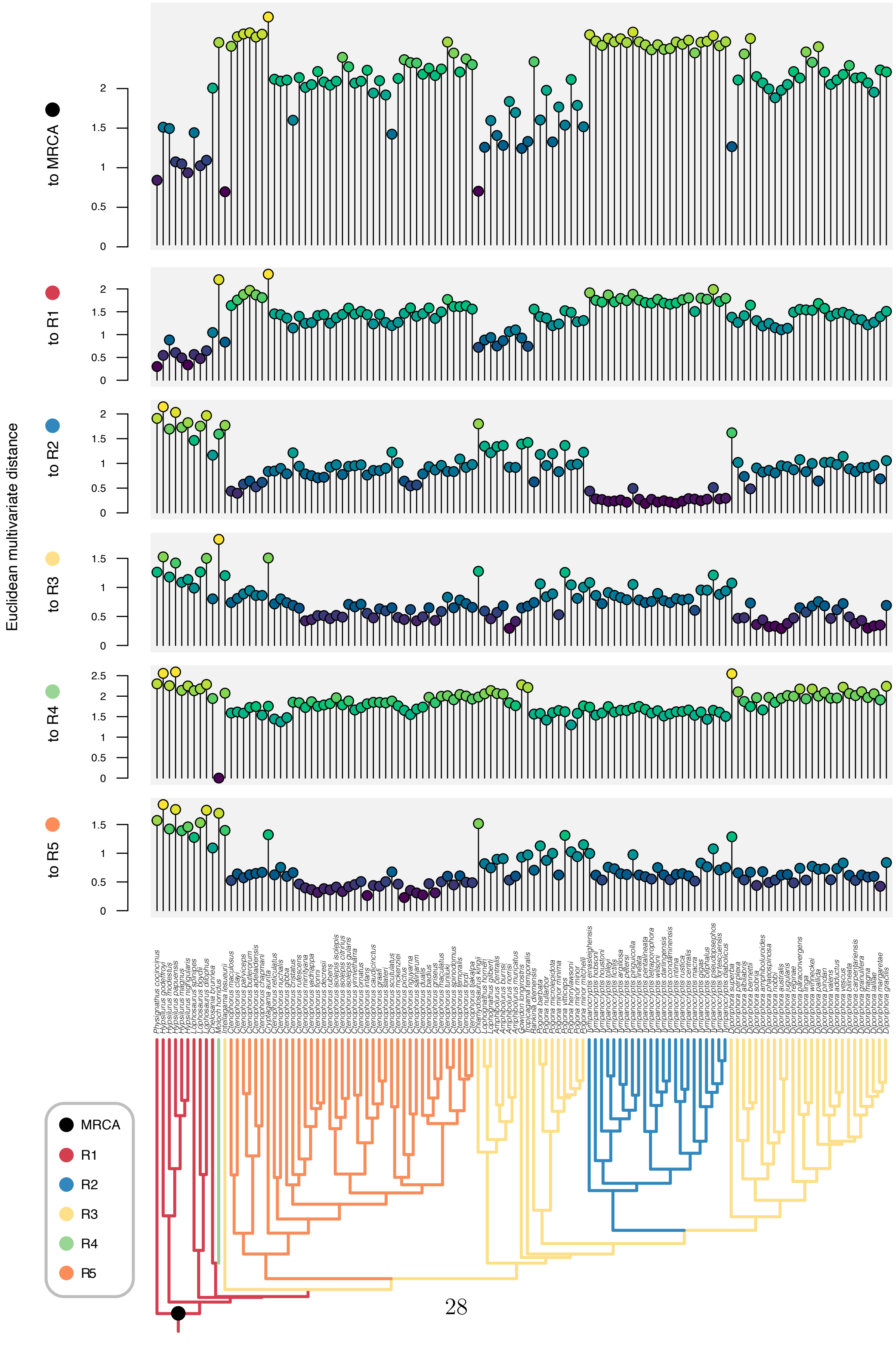

Euclidean multivariate distance
to MRCA
to R1
to R2
to R3
to R4
to R5
0
0.5
1
1.5
2
2.5
MRCA
R1
R2
R3
R4
R5

**Figure S12.** Multivariate Euclidean distance estimated between each extant species and estimated optima, as well as to the amphibolurine MRCA. Tree at bottom is colored according to the k = 5 model of morphological optima. Lollipop plots above show morphological distance between each extant species and the amphibolurine MRCA (black), primarily arboreal regime 1 (red), regime 2 (blue; *Tympanocryptis*), the primarily generalist regime 3 (yellow), regime 4 (green; *Moloch*), and regime 5 (orange; *Ctenophorus*). Based on this metric and our measured traits, the amphibolurine MRCA was most morphologically similar to *Intellagama*, *Physignathus*, and *Chlamydosaurus*. Terrestrial forms like *Ctenophorus aurita*, the *Ctenophorus adelaidensis* group, and *Tympanocryptis* are likely highly derived and morphologically distinct from the MRCA.

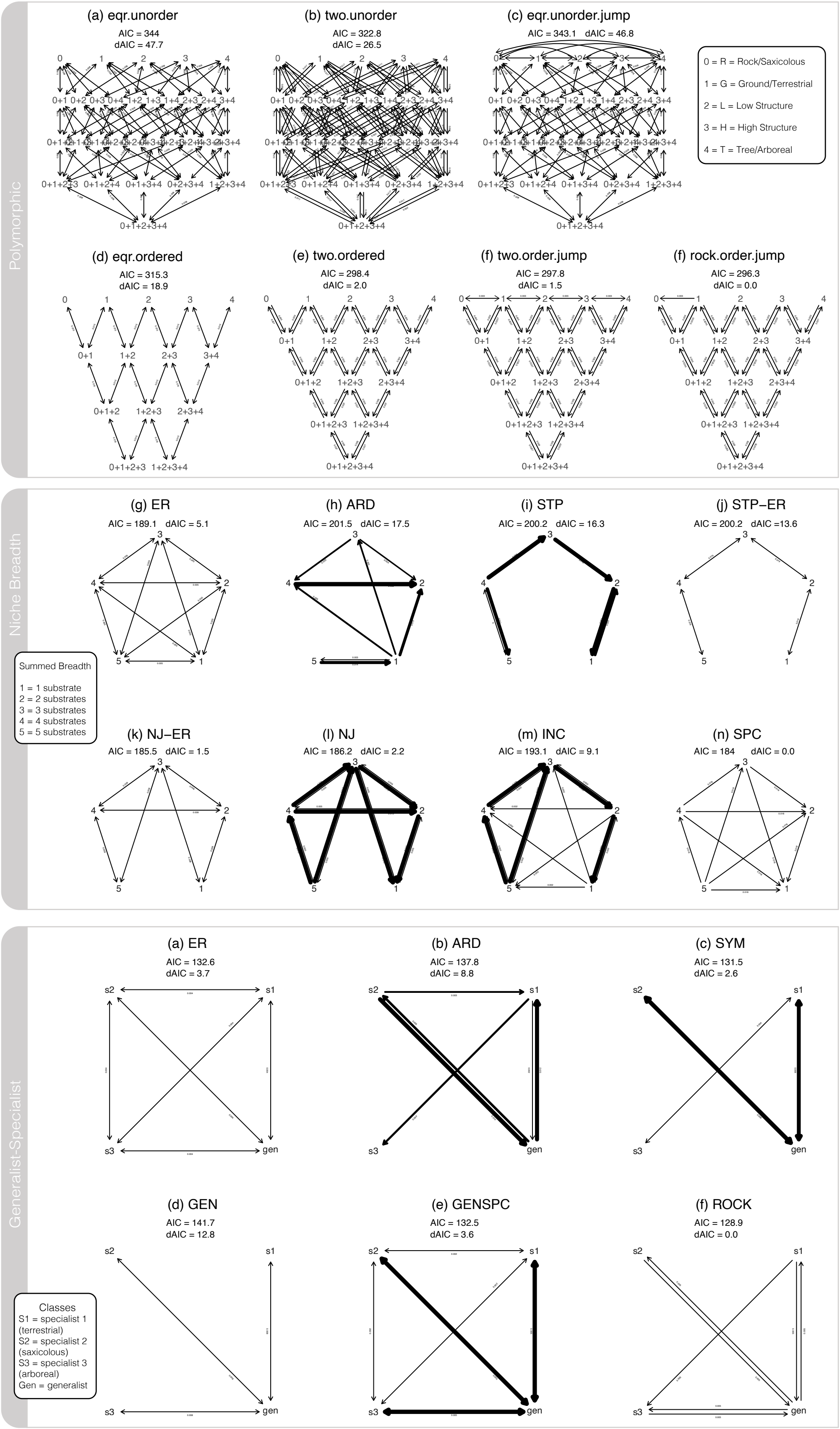

**Figure S13.** See caption below.

**Figure S13.** Visualizations of models fit to substrate usage, niche breadth, and niche transitions along the generalist to specialist continuum. Complete sets of Markov models compared, resulting in favored models (grey highlight) presented in Figure 2. Line width indicates transition rate. Upper panel depicts models used to understand niche generalisation and specialisation by modelling the evolution of character states using polymorphic Markov models. Middle panel depicts evolutionary models for understanding transitions in niche breadth (measured as the sum of used substrates 1-5). Bottom panel depicts models fit to understand evolution between three specialist classes (S1, S2, S3 with niche breadth = 1) and generalists (niche breadth >= 3). Ancestral state estimations for each model set were weight averaged across all models (see Fig.S16-S18).

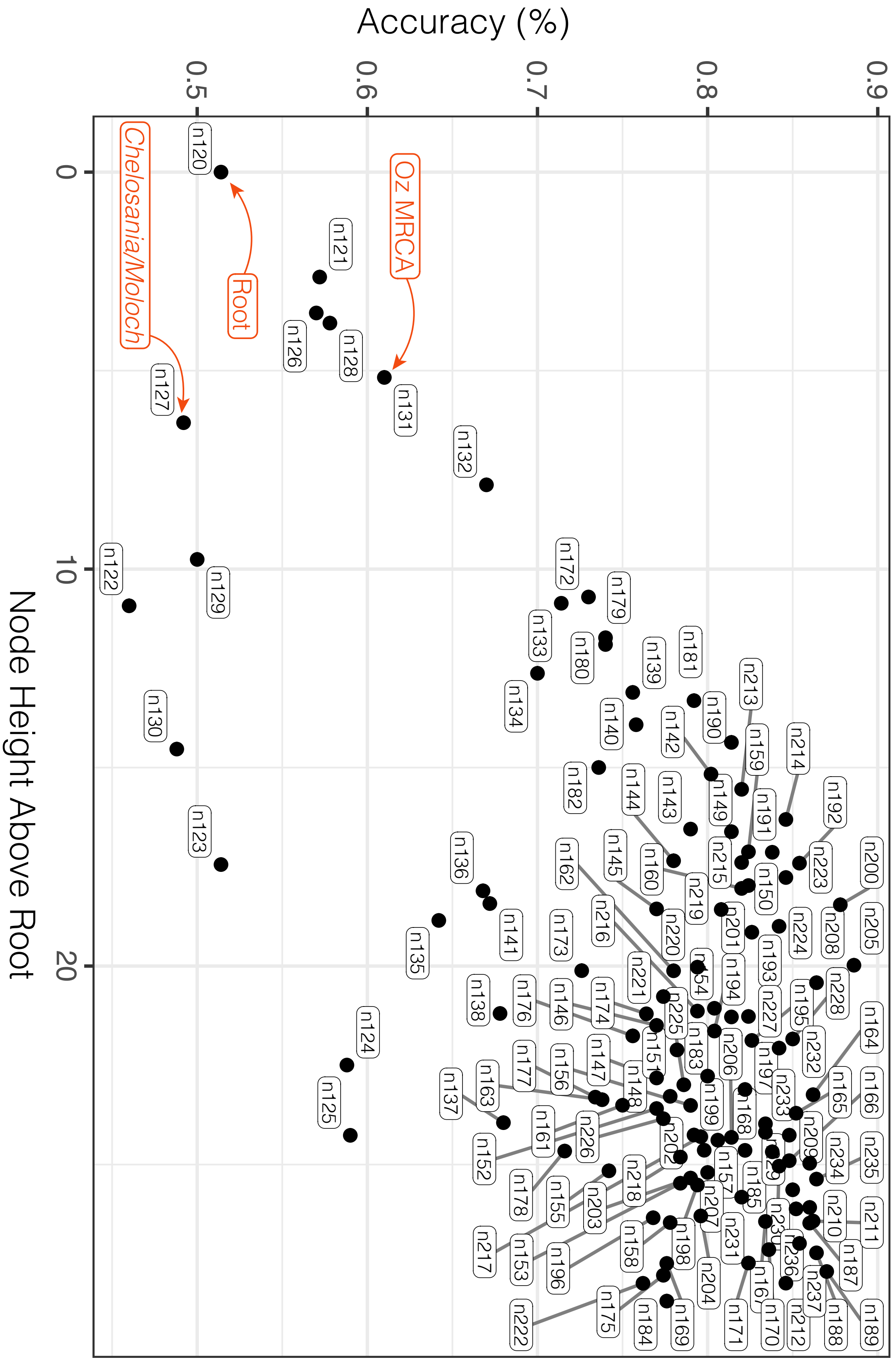

**Figure S14.** See caption below.

**Figure S14.** Summary of the RandomForest simulation exercise to explore bias and sensitivity. Results show the median accuracy of recovering the known trait state from 500 simulations using RandomForests. Accuracy is typically high (>70%) but there is some decrease in accuracy as we look at branches deeper in the phylogeny. We highlight three nodes of interest: the amphioblurine root, the Australian MRCA and the split between Chelosania and Moloch as reference points that identify the consistent difficulty in estimating some nodes (root, Chelosania/Moloch) and increased confidence in others (Oz MRCA).

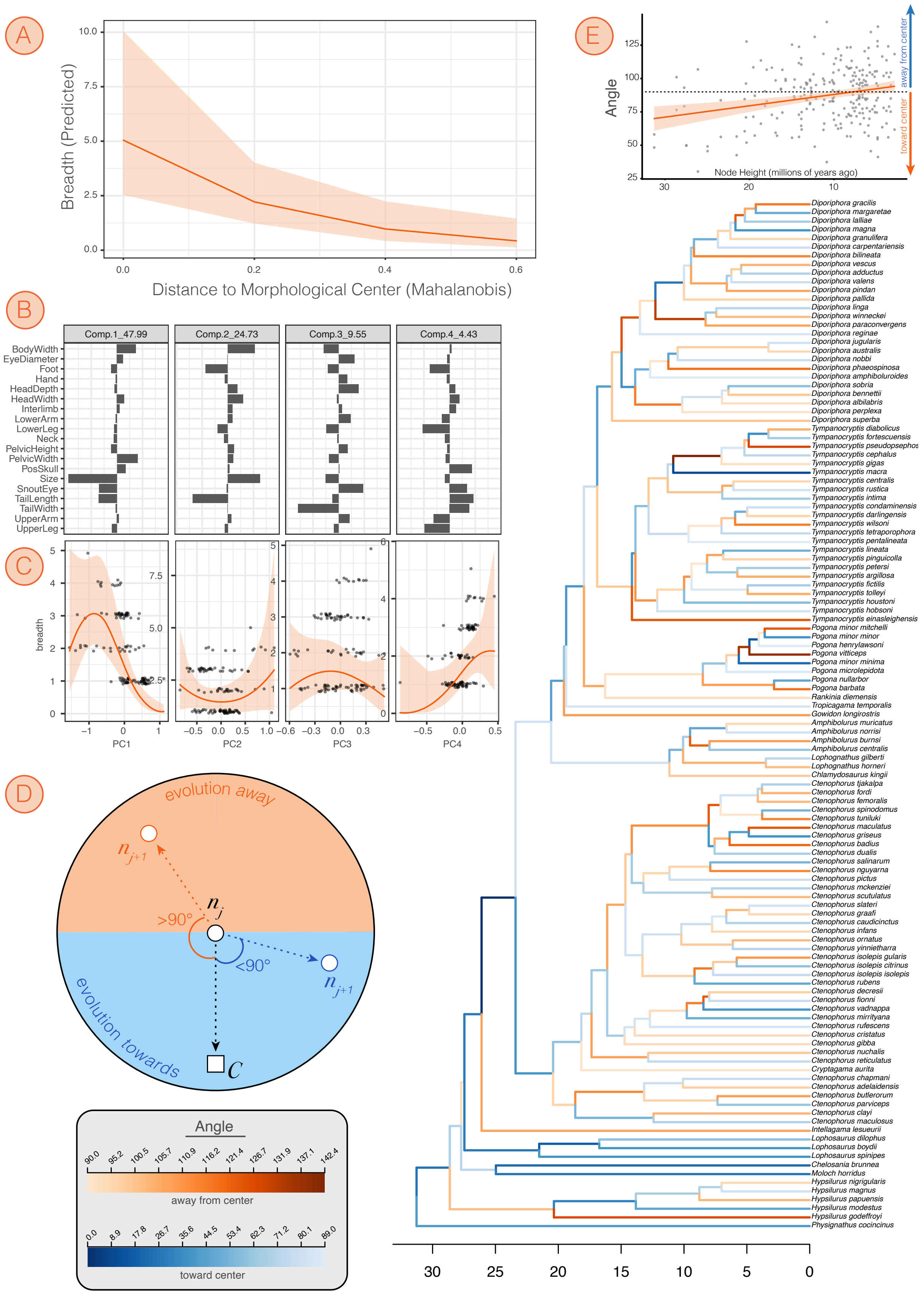

**Figure S15.** See caption below..

**Figure S15.** (A) Plot of habitat usage breadth as a function of Mahalanobis distance to the morphological center. Breadth scores are integers and jitter around values is solely to improve visualization of concentration of points. Orange line indicates best fit prediction resulting from the phylogenetic generalized linear mixed model. (B) Four panels indicate trait loadings on the first four Principal Component axes that account for >90% of trait variation. (C) Corresponding plots of habitat usage breadth against PC scores, with best fit GLMM lines and confidence intervals overlaid in orange. (D) Demonstration of how the angle of morphological change was calculated between an ancestral node (nj), descendant node (nj+1), and the morphological center/midpoint (C). Angles $<90°$ were classified as movement towards the center, angles $\geq 90°$ were classified as movement away from the center. Color and saturation of each branch indicates the direction of movement and value of the angle. (E) Plot of the relationship between angle of change and node age, showing biased movement towards the morphological center early in the evolution of the group. Fit of linear model overlaid.

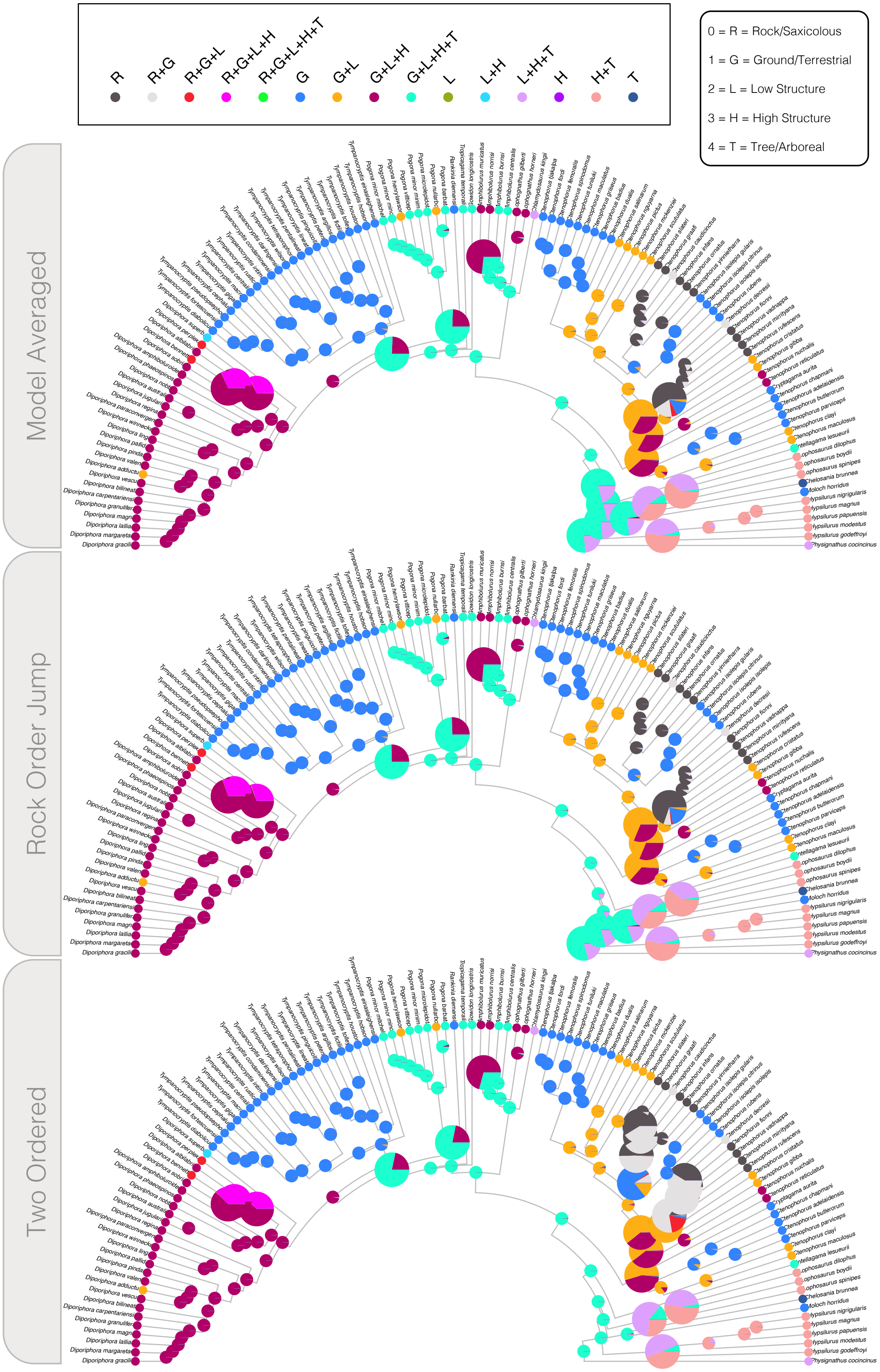

**Figure S16.** Ancestral state estimates for the **Polymorphism** models under (top) weighted model averages, (middle) the top model 'Rock Order Jump', and (bottom) the second best model 'Two Ordered'.

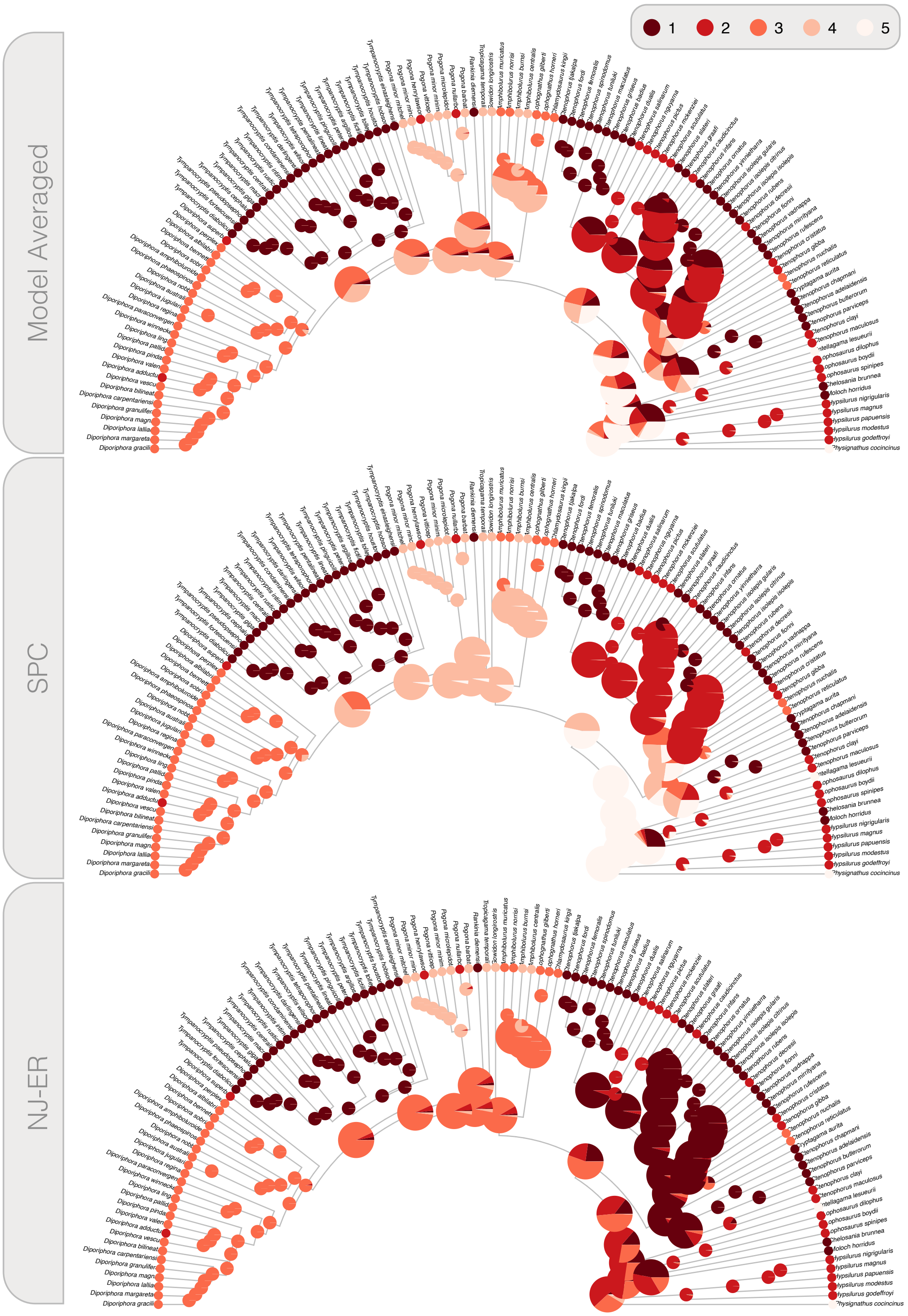

**Figure S17.** Ancestral state estimates for the **Niche Breadth** models under (top) weighted model averages, (middle) the top model 'SPC', and (bottom) the second best model 'NJ-ER'.

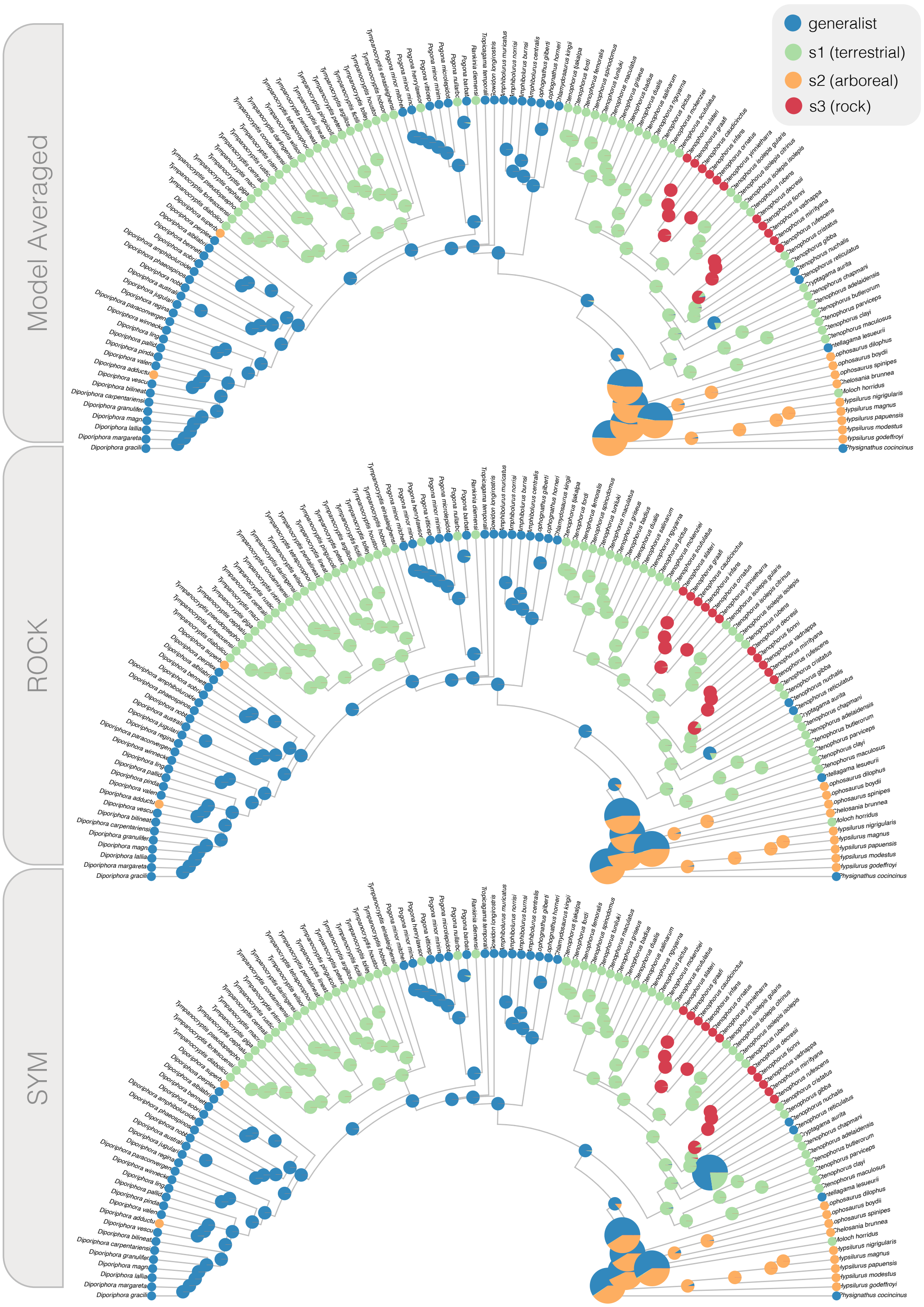

**Figure S18.** Ancestral state estimates for the **Generalist-Specialist** models under (top) weighted model averages, (middle) the top model 'ROCK', and (bottom) the second best model 'SYM'.

### Software

All software and versions used for *pipesnake* V.12 assembly and analysis of molecular data.

```
ASSEMBLY_POSTPROCESSING:
  python: 3.8.3
ASTER:
  astral-hybrid: 1.16.3.4
BBMAP_DEDUPE: BBMAP_FILTER: BBMAP_REFORMAT:
  BBMAP - clumpify.sh: '39.01'
  BBMAP - reformat.sh: '39.01'
  BBMAP - bbmap.sh: ''
BLAT:
  blat: '36'
CUSTOM_DUMPSOFTWAREVERSIONS:
  python: 3.10.6
  yaml: '6.0'
GBLOCKS:
  gblocks: 0.91b
IQTREE:
  iqtree: 2.2.6
MAFFT:
  mafft: '7.221'
MAKE_PRG:
  python: 3.8.3
MERGE_TREES:
  BusyBox: v1.22.1
PARSE_BLAT_RESULTS:
  python: 3.8.3
PEAR:
  pear: 0.9.6)
PHYLOGENY_MAKE_ALIGNMENTS:
  python: 3.8.3
PREPARE_ADAPTOR:
  python: 3.8.3
QUALITY_2_ASSEMBLY:
  python: 3.8.3
SED:
  sed: This is not GNU sed version 4.0
SPADES:
  spades: 3.15.5
TRIMMOMATIC:
  trimmomatic: '0.39'
TRIMMOMATIC_CLEAN_PE:
  trimmomatic: '0.39'
Workflow:
  Nextflow: 23.04.4
  ausarg/pipesnake: '1.2'
```

### Supplementary Tables

#### Table S1. Molecular Sampling

| Family | Subfamily | Genus | Species | RegNo | AHE | gene | uce | total_loci | Source |
|---|---|---|---|---|---|---|---|---|---|
| outgroup | — | Gallus | gallus | SAMN15960293 | 368 | 38 | 653 | 1059 | NCBI |
| Sphenodontidae | — | Sphenodon | punctatus | SAMN08038466 | 385 | 39 | 4992 | 5416 | NCBI |
| Sphenodontidae | — | Sphenodon | punctatus | SAMN08038466 | 385 | 39 | 4992 | 5416 | NCBI |
| Varanidae | — | Varanus | komodoensis | SAMN10967258 | 388 | 41 | 4997 | 5426 | NCBI |
| Dibamidae | — | Dibamus | novaeguineae | ABTC062295 | 338 | 0 | 2 | 340 | Burbrink et al. 2021 (AHE_T212) |
| Gekkonidae | — | Heteronotia | binoei | CCM8104 | 388 | 40 | 4984 | 5412 | NCBI |
| Agamidae | Agaminae | Acanthocercus | guentherpetersi | TJC1385 | 360 | 35 | 4791 | 5186 | Title & Singhal et al. (2024) |
| Agamidae | Agaminae | Agama | lanzai | CAS227496 | 351 | 33 | 4823 | 5207 | Title & Singhal et al. (2024) |
| Agamidae | Agaminae | Laudakia | agrorensis | CAS232150 | 362 | 34 | 4755 | 5151 | Title & Singhal et al. (2024) |
| Agamidae | Agaminae | Laudakia | melanura | MVZHerp248402 | 360 | 34 | 4785 | 5179 | Title & Singhal et al. (2024) |
| Agamidae | Agaminae | Phrynocephalus | luteoguttatus | CAS232112 | 366 | 34 | 4804 | 5204 | Title & Singhal et al. (2024) |
| Agamidae | Agaminae | Phrynocephalus | putjatai | SAMN38108945 | 385 | 40 | 4891 | 5316 | Title & Singhal et al. (2024) |
| Agamidae | Agaminae | Pseudotrapelus | dhofarensis | CAS227583 | 361 | 34 | 4803 | 5198 | Title & Singhal et al. (2024) |
| Agamidae | Agaminae | Pseudotrapelus | jensvindumi | CAS225340 | 355 | 36 | 4782 | 5173 | Title & Singhal et al. (2024) |
| Agamidae | Agaminae | Trapelus | agilis | CAS228565 | 25 | 25 | 3948 | 3998 | Title & Singhal et al. (2024) |
| Agamidae | Agaminae | Xenagama | zonura | TJC1334 | 356 | 34 | 4807 | 5197 | Title & Singhal et al. (2024) |
| Agamidae | Amphibolurinae | Amphibolurus | burnsi | AM9736 | 345 | 28 | 4027 | 4400 | AusARG_Agamidae |
| Agamidae | Amphibolurinae | Amphibolurus | burnsi | NMVD74137 | 356 | 33 | 4674 | 5063 | AusARG_Agamidae |
| Agamidae | Amphibolurinae | Amphibolurus | centralis | NMVD72710 | 292 | 18 | 2698 | 3008 | AusARG_Agamidae |
| Agamidae | Amphibolurinae | Amphibolurus | centralis | NMVD74288 | 359 | 34 | 4572 | 4965 | AusARG_Agamidae |
| Agamidae | Amphibolurinae | Amphibolurus | muricatus | AM391L | 345 | 29 | 4452 | 4826 | AusARG_Agamidae |
| Agamidae | Amphibolurinae | Amphibolurus | muricatus | AMSR.147368 | 364 | 35 | 4752 | 5151 | AusARG_Agamidae |
| Agamidae | Amphibolurinae | Amphibolurus | muricatus | AMSR.151559 | 343 | 26 | 4088 | 4457 | AusARG_Agamidae |
| Agamidae | Amphibolurinae | Amphibolurus | muricatus | AMSR.156047 | 363 | 34 | 4763 | 5160 | AusARG_Agamidae |
| Agamidae | Amphibolurinae | Amphibolurus | norrisi | WAMR140534 | 362 | 34 | 4711 | 5107 | AusARG_Agamidae |
| Agamidae | Amphibolurinae | Amphibolurus | norrisi | WAMR77934 | 362 | 36 | 4733 | 5131 | AusARG_Agamidae |
| Agamidae | Amphibolurinae | Chelosania | brunnea | ABTC80828 | 189 | 9 | 2970 | 3168 | Title & Singhal et al. (2024) |
| Agamidae | Amphibolurinae | Chelosania | brunnea | WAMR163197 | 357 | 34 | 4698 | 5089 | AusARG_Agamidae |
| Agamidae | Amphibolurinae | Chelosania | brunnea | WAMR163197 | 364 | 35 | 4718 | 5117 | AusARG_Agamidae |
| Agamidae | Amphibolurinae | Chlamydosaurus | kingii | AMSR.140277 | 366 | 34 | 4725 | 5125 | AusARG_Agamidae |
| Agamidae | Amphibolurinae | Chlamydosaurus | kingii | NMVD74308 | 368 | 33 | 4737 | 5138 | AusARG_Agamidae |
| Agamidae | Amphibolurinae | Chlamydosaurus | kingii | UMMZ236528 | 369 | 35 | 4808 | 5212 | Title & Singhal et al. (2024) |
| Agamidae | Amphibolurinae | Cryptagama | aurita | TR1103 | 359 | 31 | 4607 | 4997 | AusARG_Agamidae |
| Agamidae | Amphibolurinae | Cryptagama | aurita | TR1107 | 364 | 36 | 4744 | 5144 | AusARG_Agamidae |
| Agamidae | Amphibolurinae | Ctenophorus | adelaidensis | WAMR151705 | 361 | 35 | 4781 | 5177 | AusARG_Agamidae |
| Agamidae | Amphibolurinae | Ctenophorus | adelaidensis | WAMR154000 | 363 | 34 | 4754 | 5151 | AusARG_Agamidae |
| Agamidae | Amphibolurinae | Ctenophorus | badius | SAMAR29301 | 356 | 35 | 4770 | 5161 | AusARG_Agamidae |
| Agamidae | Amphibolurinae | Ctenophorus | badius | WAMR121913 | 241 | 12 | 1991 | 2244 | AusARG_Agamidae |
| Agamidae | Amphibolurinae | Ctenophorus | butlerorum | WAMR113629 | 313 | 22 | 3339 | 3674 | AusARG_Agamidae |
| Agamidae | Amphibolurinae | Ctenophorus | butlerorum | WAMR121917 | 368 | 36 | 4803 | 5207 | AusARG_Agamidae |
| Agamidae | Amphibolurinae | Ctenophorus | caudicinctus | NMVD74396 | 357 | 35 | 4794 | 5186 | AusARG_Agamidae |
| Agamidae | Amphibolurinae | Ctenophorus | caudicinctus | WAMR154520 | 326 | 23 | 3834 | 4183 | AusARG_Agamidae |
| Agamidae | Amphibolurinae | Ctenophorus | caudicinctus.caudicinctus.A | NMVD74376 | 251 | 11 | 1680 | 1942 | AusARG_Agamidae |
| Agamidae | Amphibolurinae | Ctenophorus | caudicinctus.caudicinctus.A | NMVD74376 | 251 | 11 | 1680 | 1942 | Title & Singhal et al. (2024) |
| Agamidae | Amphibolurinae | Ctenophorus | caudicinctus.caudicinctus.B | NMVD74373 | 363 | 33 | 4715 | 5111 | AusARG_Agamidae |
| Agamidae | Amphibolurinae | Ctenophorus | caudicinctus.mensarum | NMVD74398 | 265 | 16 | 2483 | 2764 | AusARG_Agamidae |
| Agamidae | Amphibolurinae | Ctenophorus | caudicinctus.mensarum | WAMR116892 | 367 | 34 | 4786 | 5187 | AusARG_Agamidae |
| Agamidae | Amphibolurinae | Ctenophorus | chapmani | WAMR144221 | 361 | 35 | 4810 | 5206 | AusARG_Agamidae |
| Agamidae | Amphibolurinae | Ctenophorus | chapmani | WAMR154321 | 364 | 35 | 4809 | 5208 | AusARG_Agamidae |

*(continued)*

| Family | Subfamily | Genus | Species | RegNo | AHE | gene | uce | total_loci | Source |
|---|---|---|---|---|---|---|---|---|---|
| Agamidae | Amphibolurinae | Ctenophorus | clayi | NMVZ54524 | 362 | 36 | 4725 | 5123 | AusARG_Agamidae |
| Agamidae | Amphibolurinae | Ctenophorus | clayi | NMVZ75788 | 365 | 36 | 4757 | 5158 | AusARG_Agamidae |
| Agamidae | Amphibolurinae | Ctenophorus | cristatus | NMVZ75787 | 366 | 36 | 4786 | 5188 | AusARG_Agamidae |
| Agamidae | Amphibolurinae | Ctenophorus | cristatus | SAMAR31851 | 363 | 35 | 4692 | 5090 | AusARG_Agamidae |
| Agamidae | Amphibolurinae | Ctenophorus | decresii | SAMAR49234 | 365 | 34 | 4681 | 5080 | AusARG_Agamidae |
| Agamidae | Amphibolurinae | Ctenophorus | decresii | SAMAR53785 | 361 | 33 | 4729 | 5123 | AusARG_Agamidae |
| Agamidae | Amphibolurinae | Ctenophorus | dualis | SAMAR59605 | 364 | 32 | 4655 | 5051 | AusARG_Agamidae |
| Agamidae | Amphibolurinae | Ctenophorus | dualis | WAMR142341 | 366 | 34 | 4753 | 5153 | AusARG_Agamidae |
| Agamidae | Amphibolurinae | Ctenophorus | femoralis | SAMAR22804 | 355 | 32 | 4163 | 4550 | AusARG_Agamidae |
| Agamidae | Amphibolurinae | Ctenophorus | femoralis | WAMR120527 | 359 | 33 | 4421 | 4813 | AusARG_Agamidae |
| Agamidae | Amphibolurinae | Ctenophorus | fionni | SAMAR30170 | 365 | 35 | 4726 | 5126 | AusARG_Agamidae |
| Agamidae | Amphibolurinae | Ctenophorus | fionni | SAMAR49107 | 360 | 35 | 4751 | 5146 | AusARG_Agamidae |
| Agamidae | Amphibolurinae | Ctenophorus | fordi | NMVD65877 | 365 | 36 | 4724 | 5125 | AusARG_Agamidae |
| Agamidae | Amphibolurinae | Ctenophorus | fordi | WAMR141036 | 356 | 31 | 4513 | 4900 | AusARG_Agamidae |
| Agamidae | Amphibolurinae | Ctenophorus | gibba | NMVZ53191 | 357 | 35 | 4703 | 5095 | AusARG_Agamidae |
| Agamidae | Amphibolurinae | Ctenophorus | gibba | SAMAR45990 | 194 | 11 | 1216 | 1421 | AusARG_Agamidae |
| Agamidae | Amphibolurinae | Ctenophorus | graafi | ABTC91400 | 367 | 34 | 4727 | 5128 | AusARG_Agamidae |
| Agamidae | Amphibolurinae | Ctenophorus | graafi | ABTC91543 | 351 | 29 | 4439 | 4819 | AusARG_Agamidae |
| Agamidae | Amphibolurinae | Ctenophorus | griseus | WAMR142324 | 354 | 26 | 3985 | 4365 | AusARG_Agamidae |
| Agamidae | Amphibolurinae | Ctenophorus | griseus | WAMR142353 | 351 | 29 | 4182 | 4562 | AusARG_Agamidae |
| Agamidae | Amphibolurinae | Ctenophorus | ibiri | SAMAR36504 | 266 | 20 | 2420 | 2706 | AusARG_Agamidae |
| Agamidae | Amphibolurinae | Ctenophorus | ibiri | SAMAR57428 | 362 | 32 | 4477 | 4871 | AusARG_Agamidae |
| Agamidae | Amphibolurinae | Ctenophorus | infans | WAMR119039 | 362 | 35 | 4746 | 5143 | AusARG_Agamidae |
| Agamidae | Amphibolurinae | Ctenophorus | infans | WAMR119041 | 364 | 34 | 4687 | 5085 | AusARG_Agamidae |
| Agamidae | Amphibolurinae | Ctenophorus | isolepis | UMMZ244206 | 360 | 31 | 4817 | 5208 | Title & Singhal et al. (2024) |
| Agamidae | Amphibolurinae | Ctenophorus | isolepis.citrinus | NMVD74451 | 362 | 34 | 4702 | 5098 | AusARG_Agamidae |
| Agamidae | Amphibolurinae | Ctenophorus | isolepis.gularis | NMVD74425 | 368 | 32 | 4698 | 5098 | AusARG_Agamidae |
| Agamidae | Amphibolurinae | Ctenophorus | isolepis.gularis | SAMAR26194 | 363 | 34 | 4642 | 5039 | AusARG_Agamidae |
| Agamidae | Amphibolurinae | Ctenophorus | isolepis.gularis | WAMR164249 | 368 | 34 | 4779 | 5181 | AusARG_Agamidae |
| Agamidae | Amphibolurinae | Ctenophorus | isolepis.isolepis | WAMR110090 | 364 | 34 | 4781 | 5179 | AusARG_Agamidae |
| Agamidae | Amphibolurinae | Ctenophorus | isolepis.isolepis | WAMR159813 | 364 | 34 | 4779 | 5177 | AusARG_Agamidae |
| Agamidae | Amphibolurinae | Ctenophorus | kartiwarru | SAMAR30254 | 367 | 32 | 4642 | 5041 | AusARG_Agamidae |
| Agamidae | Amphibolurinae | Ctenophorus | kartiwarru | SAMAR49254 | 355 | 29 | 4365 | 4749 | AusARG_Agamidae |
| Agamidae | Amphibolurinae | Ctenophorus | maculatus | AMSR.140523 | 360 | 33 | 4701 | 5094 | AusARG_Agamidae |
| Agamidae | Amphibolurinae | Ctenophorus | maculatus | WAMR142114 | 355 | 31 | 4349 | 4735 | AusARG_Agamidae |
| Agamidae | Amphibolurinae | Ctenophorus | maculatus | WAMR142114 | 355 | 31 | 4349 | 4735 | AusARG_Agamidae |
| Agamidae | Amphibolurinae | Ctenophorus | maculosus | SAMAR25909 | 329 | 24 | 3387 | 3740 | AusARG_Agamidae |
| Agamidae | Amphibolurinae | Ctenophorus | maculosus | SAMAR62749 | 364 | 33 | 4578 | 4975 | AusARG_Agamidae |
| Agamidae | Amphibolurinae | Ctenophorus | mckenziei | SAMAR32266 | 366 | 33 | 4724 | 5123 | AusARG_Agamidae |
| Agamidae | Amphibolurinae | Ctenophorus | mckenziei | WAMR91852 | 363 | 31 | 4663 | 5057 | AusARG_Agamidae |
| Agamidae | Amphibolurinae | Ctenophorus | mirrityana | NMV | 371 | 35 | 4695 | 5101 | AusARG_Agamidae |
| Agamidae | Amphibolurinae | Ctenophorus | mirrityana | SAMAR31655 | 364 | 35 | 4745 | 5144 | AusARG_Agamidae |
| Agamidae | Amphibolurinae | Ctenophorus | modestus | SAMAR40708 | 365 | 34 | 4727 | 5126 | AusARG_Agamidae |
| Agamidae | Amphibolurinae | Ctenophorus | modestus | SAMAR51748 | 366 | 35 | 4760 | 5161 | AusARG_Agamidae |
| Agamidae | Amphibolurinae | Ctenophorus | modestus | SAMAR52935 | 364 | 36 | 4702 | 5102 | AusARG_Agamidae |
| Agamidae | Amphibolurinae | Ctenophorus | modestus | SAMAR60922 | 368 | 35 | 4821 | 5224 | AusARG_Agamidae |
| Agamidae | Amphibolurinae | Ctenophorus | nguyarna | WAMR157972 | 364 | 34 | 4784 | 5182 | AusARG_Agamidae |
| Agamidae | Amphibolurinae | Ctenophorus | nguyarna | WAMR157973 | 363 | 34 | 4697 | 5094 | AusARG_Agamidae |
| Agamidae | Amphibolurinae | Ctenophorus | nuchalis | CTN53 | 366 | 36 | 4756 | 5158 | AusARG_Agamidae |
| Agamidae | Amphibolurinae | Ctenophorus | nuchalis | SAMAR42726 | 365 | 33 | 4732 | 5130 | AusARG_Agamidae |
| Agamidae | Amphibolurinae | Ctenophorus | nuchalis | UMMZ244210 | 320 | 23 | 3998 | 4341 | Title & Singhal et al. (2024) |
| Agamidae | Amphibolurinae | Ctenophorus | ornatus | WAMR104404 | 362 | 33 | 4769 | 5164 | AusARG_Agamidae |
| Agamidae | Amphibolurinae | Ctenophorus | ornatus | WAMR117279 | 366 | 34 | 4766 | 5166 | AusARG_Agamidae |
| Agamidae | Amphibolurinae | Ctenophorus | ornatus | WAMR154798 | 367 | 33 | 4784 | 5184 | AusARG_Agamidae |
| Agamidae | Amphibolurinae | Ctenophorus | ornatus | WAMR84713 | 359 | 34 | 4737 | 5130 | AusARG_Agamidae |

*(continued)*

| Family | Subfamily | Genus | Species | RegNo | AHE | gene | uce | total_loci | Source |
|---|---|---|---|---|---|---|---|---|---|
| Agamidae | Amphibolurinae | Ctenophorus | parviceps | WAMR120357 | 365 | 34 | 4738 | 5137 | AusARG_Agamidae |
| Agamidae | Amphibolurinae | Ctenophorus | parviceps | WAMR120360 | 364 | 36 | 4714 | 5114 | AusARG_Agamidae |
| Agamidae | Amphibolurinae | Ctenophorus | pictus | NMVD71337 | 366 | 33 | 4765 | 5164 | AusARG_Agamidae |
| Agamidae | Amphibolurinae | Ctenophorus | pictus | SAMAR28208 | 307 | 18 | 2351 | 2676 | AusARG_Agamidae |
| Agamidae | Amphibolurinae | Ctenophorus | reticulatus | CTR20 | 361 | 35 | 4754 | 5150 | AusARG_Agamidae |
| Agamidae | Amphibolurinae | Ctenophorus | reticulatus | CTR45 | 364 | 35 | 4731 | 5130 | AusARG_Agamidae |
| Agamidae | Amphibolurinae | Ctenophorus | reticulatus | UMMZ244212 | 355 | 28 | 4802 | 5185 | Title & Singhal et al. (2024) |
| Agamidae | Amphibolurinae | Ctenophorus | rubens | WAMR127501 | 365 | 34 | 4768 | 5167 | AusARG_Agamidae |
| Agamidae | Amphibolurinae | Ctenophorus | rubens | WAMR157271 | 364 | 34 | 4719 | 5117 | AusARG_Agamidae |
| Agamidae | Amphibolurinae | Ctenophorus | rufescens | SAMAR28276 | 354 | 35 | 4680 | 5069 | AusARG_Agamidae |
| Agamidae | Amphibolurinae | Ctenophorus | rufescens | WAMR53417 | 366 | 35 | 4757 | 5158 | AusARG_Agamidae |
| Agamidae | Amphibolurinae | Ctenophorus | salinarum | NMVD74401 | 337 | 26 | 4206 | 4569 | AusARG_Agamidae |
| Agamidae | Amphibolurinae | Ctenophorus | salinarum | NMVZ54444 | 362 | 33 | 4723 | 5118 | AusARG_Agamidae |
| Agamidae | Amphibolurinae | Ctenophorus | scutulatus | NMVD74467 | 365 | 33 | 4718 | 5116 | AusARG_Agamidae |
| Agamidae | Amphibolurinae | Ctenophorus | scutulatus | NMVZ54544 | 354 | 29 | 4587 | 4970 | AusARG_Agamidae |
| Agamidae | Amphibolurinae | Ctenophorus | scutulatus | UMMZ244218 | 357 | 30 | 4823 | 5210 | Title & Singhal et al. (2024) |
| Agamidae | Amphibolurinae | Ctenophorus | slateri | NACCM2282 | 371 | 34 | 4810 | 5215 | Title & Singhal et al. (2024) |
| Agamidae | Amphibolurinae | Ctenophorus | slateri | NMVD72581 | 364 | 34 | 4739 | 5137 | AusARG_Agamidae |
| Agamidae | Amphibolurinae | Ctenophorus | slateri | NMVD72687 | 349 | 30 | 4418 | 4797 | AusARG_Agamidae |
| Agamidae | Amphibolurinae | Ctenophorus | slateri | NMVD72688 | 339 | 27 | 4331 | 4697 | AusARG_Agamidae |
| Agamidae | Amphibolurinae | Ctenophorus | slateri | NMVD74055 | 355 | 29 | 4426 | 4810 | AusARG_Agamidae |
| Agamidae | Amphibolurinae | Ctenophorus | slateri | NMVD74268 | 364 | 33 | 4722 | 5119 | AusARG_Agamidae |
| Agamidae | Amphibolurinae | Ctenophorus | slateri | NMVD74284 | 364 | 34 | 4761 | 5159 | AusARG_Agamidae |
| Agamidae | Amphibolurinae | Ctenophorus | spinodomus | AMSR.156634 | 368 | 34 | 4783 | 5185 | AusARG_Agamidae |
| Agamidae | Amphibolurinae | Ctenophorus | spinodomus | SAMAR36978 | 367 | 36 | 4734 | 5137 | AusARG_Agamidae |
| Agamidae | Amphibolurinae | Ctenophorus | tjakalpa | SAMAR31888 | 351 | 29 | 4405 | 4785 | AusARG_Agamidae |
| Agamidae | Amphibolurinae | Ctenophorus | tjakalpa | WAMR140408 | 367 | 33 | 4726 | 5126 | AusARG_Agamidae |
| Agamidae | Amphibolurinae | Ctenophorus | tjantjalka | ABTC33798 | 367 | 35 | 4730 | 5132 | AusARG_Agamidae |
| Agamidae | Amphibolurinae | Ctenophorus | tjantjalka | SAMAR46211 | 357 | 30 | 4176 | 4563 | AusARG_Agamidae |
| Agamidae | Amphibolurinae | Ctenophorus | tuniluki | SAMAR22756 | 302 | 20 | 2681 | 3003 | AusARG_Agamidae |
| Agamidae | Amphibolurinae | Ctenophorus | tuniluki | SAMAR22950 | 347 | 30 | 3959 | 4336 | AusARG_Agamidae |
| Agamidae | Amphibolurinae | Ctenophorus | vadnappa | SAMAR31656 | 364 | 36 | 4712 | 5112 | AusARG_Agamidae |
| Agamidae | Amphibolurinae | Ctenophorus | vadnappa | SAMAR36388 | 361 | 36 | 4738 | 5135 | AusARG_Agamidae |
| Agamidae | Amphibolurinae | Ctenophorus | yinnietharra | WAMR165963 | 292 | 18 | 3383 | 3693 | AusARG_Agamidae |
| Agamidae | Amphibolurinae | Ctenophorus | yinnietharra | WAMR165964 | 354 | 28 | 4537 | 4919 | AusARG_Agamidae |
| Agamidae | Amphibolurinae | Diporiphora | adductus | WAMR120511 | 357 | 31 | 4721 | 5109 | AusARG_Agamidae |
| Agamidae | Amphibolurinae | Diporiphora | adductus | WAMR157264 | 359 | 36 | 4769 | 5164 | AusARG_Agamidae |
| Agamidae | Amphibolurinae | Diporiphora | albilabris | NMVD73840 | 359 | 35 | 4701 | 5095 | AusARG_Agamidae |
| Agamidae | Amphibolurinae | Diporiphora | albilabris | NMVD73858 | 234 | 17 | 909 | 1160 | AusARG_Agamidae |
| Agamidae | Amphibolurinae | Diporiphora | ameliae | QMJ88274 | 358 | 33 | 4678 | 5069 | AusARG_Agamidae |
| Agamidae | Amphibolurinae | Diporiphora | ameliae | QMJ88275 | 366 | 36 | 4720 | 5122 | AusARG_Agamidae |
| Agamidae | Amphibolurinae | Diporiphora | amphiboluroides | NMVZ75792 | 369 | 33 | 4724 | 5126 | AusARG_Agamidae |
| Agamidae | Amphibolurinae | Diporiphora | amphiboluroides | WAMR104419 | 339 | 32 | 4417 | 4788 | AusARG_Agamidae |
| Agamidae | Amphibolurinae | Diporiphora | australis | NMVD74097 | 366 | 34 | 4795 | 5195 | AusARG_Agamidae |
| Agamidae | Amphibolurinae | Diporiphora | australis | NMVD74102 | 359 | 35 | 4730 | 5124 | AusARG_Agamidae |
| Agamidae | Amphibolurinae | Diporiphora | bennettii | ANWC | 359 | 36 | 4681 | 5076 | AusARG_Agamidae |
| Agamidae | Amphibolurinae | Diporiphora | bennettii | WAMR168157 | 363 | 35 | 4800 | 5198 | AusARG_Agamidae |
| Agamidae | Amphibolurinae | Diporiphora | bilineata | NMVD72632 | 361 | 36 | 4798 | 5195 | AusARG_Agamidae |
| Agamidae | Amphibolurinae | Diporiphora | bilineata | NMVD74266 | 358 | 30 | 4679 | 5067 | AusARG_Agamidae |
| Agamidae | Amphibolurinae | Diporiphora | carpentariaensis | NMVD74068 | 363 | 34 | 4718 | 5115 | AusARG_Agamidae |
| Agamidae | Amphibolurinae | Diporiphora | carpentariaensis | NMVD74079 | 362 | 35 | 4776 | 5173 | AusARG_Agamidae |
| Agamidae | Amphibolurinae | Diporiphora | gracilis | ANU | 364 | 32 | 4747 | 5143 | AusARG_Agamidae |
| Agamidae | Amphibolurinae | Diporiphora | gracilis | NMVD74328 | 360 | 34 | 4764 | 5158 | AusARG_Agamidae |
| Agamidae | Amphibolurinae | Diporiphora | granulifera | ANU | 363 | 34 | 4779 | 5176 | AusARG_Agamidae |
| Agamidae | Amphibolurinae | Diporiphora | granulifera | NMVD74047 | 362 | 33 | 4749 | 5144 | AusARG_Agamidae |

(continued)

| Family | Subfamily | Genus | Species | RegNo | AHE | gene | uce | total_loci | Source |
|---|---|---|---|---|---|---|---|---|---|
| Agamidae | Amphibolurinae | Diporiphora | jugularis | NMVD74101 | 338 | 28 | 4440 | 4806 | AusARG_Agamidae |
| Agamidae | Amphibolurinae | Diporiphora | jugularis | QMJ95583 | 360 | 35 | 4757 | 5152 | AusARG_Agamidae |
| Agamidae | Amphibolurinae | Diporiphora | lalliae | NMVD72718 | 352 | 34 | 4724 | 5110 | AusARG_Agamidae |
| Agamidae | Amphibolurinae | Diporiphora | lalliae | NMVD73907 | 359 | 32 | 4756 | 5147 | AusARG_Agamidae |
| Agamidae | Amphibolurinae | Diporiphora | lalliae | UMMZ242571 | 360 | 34 | 4779 | 5173 | Title & Singhal et al. (2024) |
| Agamidae | Amphibolurinae | Diporiphora | linga | NMVZ53103 | 344 | 29 | 4476 | 4849 | AusARG_Agamidae |
| Agamidae | Amphibolurinae | Diporiphora | linga | NMVZ53196 | 364 | 34 | 4729 | 5127 | AusARG_Agamidae |
| Agamidae | Amphibolurinae | Diporiphora | linga | WAMR145923 | 356 | 36 | 4489 | 4881 | AusARG_Agamidae |
| Agamidae | Amphibolurinae | Diporiphora | linga | WAMR145924 | 365 | 35 | 4747 | 5147 | AusARG_Agamidae |
| Agamidae | Amphibolurinae | Diporiphora | magna | ANU | 361 | 34 | 4734 | 5129 | AusARG_Agamidae |
| Agamidae | Amphibolurinae | Diporiphora | magna | MAGNT | 358 | 32 | 4610 | 5000 | AusARG_Agamidae |
| Agamidae | Amphibolurinae | Diporiphora | magna | SAMAR10135 | 360 | 35 | 4780 | 5175 | AusARG_Agamidae |
| Agamidae | Amphibolurinae | Diporiphora | margaretae | NMVD73834 | 360 | 35 | 4744 | 5139 | AusARG_Agamidae |
| Agamidae | Amphibolurinae | Diporiphora | margaretae | NMVD73849 | 358 | 34 | 4748 | 5140 | AusARG_Agamidae |
| Agamidae | Amphibolurinae | Diporiphora | nobbi | NMVD71319 | 363 | 36 | 4718 | 5117 | AusARG_Agamidae |
| Agamidae | Amphibolurinae | Diporiphora | pallida | WAMR177292 | 358 | 34 | 4760 | 5152 | AusARG_Agamidae |
| Agamidae | Amphibolurinae | Diporiphora | paraconvergens | WAMR102709 | 360 | 34 | 4708 | 5102 | AusARG_Agamidae |
| Agamidae | Amphibolurinae | Diporiphora | paraconvergens | WAMR131073 | 361 | 35 | 4758 | 5154 | AusARG_Agamidae |
| Agamidae | Amphibolurinae | Diporiphora | paraconvergens | WAMR145070 | 360 | 36 | 4752 | 5148 | AusARG_Agamidae |
| Agamidae | Amphibolurinae | Diporiphora | paraconvergens | WAMR145457 | 363 | 35 | 4754 | 5152 | AusARG_Agamidae |
| Agamidae | Amphibolurinae | Diporiphora | paraconvergens | WAMR157943 | 360 | 35 | 4800 | 5195 | AusARG_Agamidae |
| Agamidae | Amphibolurinae | Diporiphora | paraconvergens | WAMR163948 | 357 | 32 | 4723 | 5112 | AusARG_Agamidae |
| Agamidae | Amphibolurinae | Diporiphora | perplexa | ANU | 363 | 34 | 4782 | 5179 | AusARG_Agamidae |
| Agamidae | Amphibolurinae | Diporiphora | perplexa | ANWC | 358 | 33 | 4694 | 5085 | AusARG_Agamidae |
| Agamidae | Amphibolurinae | Diporiphora | perplexa | NMVD73803 | 336 | 29 | 2227 | 2592 | AusARG_Agamidae |
| Agamidae | Amphibolurinae | Diporiphora | perplexa | NMVD73978 | 362 | 35 | 4747 | 5144 | AusARG_Agamidae |
| Agamidae | Amphibolurinae | Diporiphora | phaeospinosa | AMSR.151842 | 203 | 11 | 560 | 774 | AusARG_Agamidae |
| Agamidae | Amphibolurinae | Diporiphora | phaeospinosa | NMVD74128 | 336 | 27 | 2443 | 2806 | AusARG_Agamidae |
| Agamidae | Amphibolurinae | Diporiphora | pindan | NMVD74316 | 360 | 35 | 4784 | 5179 | AusARG_Agamidae |
| Agamidae | Amphibolurinae | Diporiphora | pindan | NMVD74348 | 357 | 35 | 4806 | 5198 | AusARG_Agamidae |
| Agamidae | Amphibolurinae | Diporiphora | reginae | NMVZ53285 | 351 | 31 | 4617 | 4999 | AusARG_Agamidae |
| Agamidae | Amphibolurinae | Diporiphora | reginae | WAMR112653 | 356 | 36 | 4752 | 5144 | AusARG_Agamidae |
| Agamidae | Amphibolurinae | Diporiphora | sobria | NMVD72706 | 350 | 33 | 4290 | 4673 | AusARG_Agamidae |
| Agamidae | Amphibolurinae | Diporiphora | sobria | NMVD73985 | 362 | 34 | 4735 | 5131 | AusARG_Agamidae |
| Agamidae | Amphibolurinae | Diporiphora | sobria | NMVD74297 | 360 | 34 | 4514 | 4908 | AusARG_Agamidae |
| Agamidae | Amphibolurinae | Diporiphora | sobria | NMVD74297 | 344 | 29 | 3917 | 4290 | AusARG_Agamidae |
| Agamidae | Amphibolurinae | Diporiphora | superba | NMVD73871 | 302 | 21 | 1448 | 1771 | AusARG_Agamidae |
| Agamidae | Amphibolurinae | Diporiphora | superba | NMVZ19163 | 206 | 11 | 447 | 664 | AusARG_Agamidae |
| Agamidae | Amphibolurinae | Diporiphora | valens | NMVD74365 | 359 | 35 | 4767 | 5161 | AusARG_Agamidae |
| Agamidae | Amphibolurinae | Diporiphora | valens | NMVD74375 | 357 | 32 | 4704 | 5093 | AusARG_Agamidae |
| Agamidae | Amphibolurinae | Diporiphora | vescus | NMVZ54459 | 347 | 30 | 4543 | 4920 | AusARG_Agamidae |
| Agamidae | Amphibolurinae | Diporiphora | vescus | WAMR100763 | 355 | 34 | 4466 | 4855 | AusARG_Agamidae |
| Agamidae | Amphibolurinae | Diporiphora | winneckei | CUMV14268 | 362 | 33 | 4816 | 5211 | Title & Singhal et al. (2024) |
| Agamidae | Amphibolurinae | Diporiphora | winneckei | NMVD72750 | 362 | 34 | 4704 | 5100 | AusARG_Agamidae |
| Agamidae | Amphibolurinae | Diporiphora | winneckei | QMJ88273 | 362 | 35 | 4783 | 5180 | AusARG_Agamidae |
| Agamidae | Amphibolurinae | Gowidon | longirostris | NMVD72733 | 354 | 28 | 4391 | 4773 | AusARG_Agamidae |
| Agamidae | Amphibolurinae | Gowidon | longirostris | NMVD74368 | 361 | 36 | 4789 | 5186 | AusARG_Agamidae |
| Agamidae | Amphibolurinae | Gowidon | longirostris | NMVD74390 | 335 | 30 | 4315 | 4680 | AusARG_Agamidae |
| Agamidae | Amphibolurinae | Gowidon | longirostris | UMFS20601 | 368 | 35 | 4814 | 5217 | Title & Singhal et al. (2024) |
| Agamidae | Amphibolurinae | Gowidon | longirostris | WAMR110768 | 360 | 35 | 4759 | 5154 | AusARG_Agamidae |
| Agamidae | Amphibolurinae | Hypsilurus | auritus | ABTC90139 | 370 | 33 | 4804 | 5207 | Title & Singhal et al. (2024) |
| Agamidae | Amphibolurinae | Hypsilurus | magnus | ABTC128837 | 363 | 35 | 4765 | 5163 | AusARG_Agamidae |
| Agamidae | Amphibolurinae | Hypsilurus | magnus | AMSR122474 | 367 | 35 | 4859 | 5261 | Singhal et al. 2021 |
| Agamidae | Amphibolurinae | Hypsilurus | magnus | NMV | 361 | 36 | 4776 | 5173 | AusARG_Agamidae |
| Agamidae | Amphibolurinae | Hypsilurus | magnus | NMV | 366 | 35 | 4790 | 5191 | AusARG_Agamidae |

*(continued)*

| Family | Subfamily | Genus | Species | RegNo | AHE | gene | uce | total_loci | Source |
|---|---|---|---|---|---|---|---|---|---|
| Agamidae | Amphibolurinae | Hypsilurus | modestus | AMSR122434 | 367 | 34 | 4588 | 4989 | Title & Singhal et al. (2024) |
| Agamidae | Amphibolurinae | Hypsilurus | modestus | NMV | 349 | 29 | 4386 | 4764 | AusARG_Agamidae |
| Agamidae | Amphibolurinae | Hypsilurus | modestus | NMV | 367 | 34 | 4761 | 5162 | AusARG_Agamidae |
| Agamidae | Amphibolurinae | Hypsilurus | schoedei | ABTC136665 | 365 | 34 | 4825 | 5224 | Title & Singhal et al. (2024) |
| Agamidae | Amphibolurinae | Intellagama | lesueurii | ABTC32057 | 373 | 34 | 4863 | 5270 | Title & Singhal et al. (2024) |
| Agamidae | Amphibolurinae | Intellagama | lesueurii | NMVD75727 | 361 | 34 | 4747 | 5142 | AusARG_Agamidae |
| Agamidae | Amphibolurinae | Intellagama | lesueurii | NMVD76213 | 336 | 26 | 4146 | 4508 | AusARG_Agamidae |
| Agamidae | Amphibolurinae | Intellagama | lesueurii | SAMAR55588 | 352 | 33 | 4499 | 4884 | AusARG_Agamidae |
| Agamidae | Amphibolurinae | Intellagama | lesueurii | WAMR33417 | 341 | 28 | 4237 | 4606 | AusARG_Agamidae |
| Agamidae | Amphibolurinae | Lophognathus | gilberti | CCM6449 | 369 | 36 | 4832 | 5237 | Title & Singhal et al. (2024) |
| Agamidae | Amphibolurinae | Lophognathus | horneri | NMVD72652 | 361 | 35 | 4697 | 5093 | AusARG_Agamidae |
| Agamidae | Amphibolurinae | Lophognathus | horneri | WAMR132850 | 363 | 34 | 4749 | 5146 | AusARG_Agamidae |
| Agamidae | Amphibolurinae | Lophognathus | horneri | WAMR139481 | 365 | 35 | 4746 | 5146 | AusARG_Agamidae |
| Agamidae | Amphibolurinae | Lophosaurus | boydii | QMJ60630 | 368 | 33 | 4461 | 4862 | Title & Singhal et al. (2024) |
| Agamidae | Amphibolurinae | Lophosaurus | boydii | QMJ60641 | 364 | 35 | 4766 | 5165 | AusARG_Agamidae |
| Agamidae | Amphibolurinae | Lophosaurus | boydii | SAMAR55835 | 364 | 35 | 4772 | 5171 | AusARG_Agamidae |
| Agamidae | Amphibolurinae | Lophosaurus | dilophus | AMSR122449 | 369 | 34 | 4772 | 5175 | Title & Singhal et al. (2024) |
| Agamidae | Amphibolurinae | Lophosaurus | diolophis | ABTC114684 | 322 | 25 | 3845 | 4192 | AusARG_Agamidae |
| Agamidae | Amphibolurinae | Lophosaurus | diolophis | ABTC50280 | 365 | 35 | 4755 | 5155 | AusARG_Agamidae |
| Agamidae | Amphibolurinae | Lophosaurus | diolophis | ABTC90265 | 363 | 36 | 4761 | 5160 | AusARG_Agamidae |
| Agamidae | Amphibolurinae | Lophosaurus | diolophis | ABTC98703 | 364 | 36 | 4741 | 5141 | AusARG_Agamidae |
| Agamidae | Amphibolurinae | Lophosaurus | diolophis | NMV | 365 | 35 | 4767 | 5167 | AusARG_Agamidae |
| Agamidae | Amphibolurinae | Lophosaurus | spinipes | ABTC100198 | 363 | 36 | 4770 | 5169 | AusARG_Agamidae |
| Agamidae | Amphibolurinae | Lophosaurus | spinipes | AMSR130058 | 368 | 33 | 4497 | 4898 | Title & Singhal et al. (2024) |
| Agamidae | Amphibolurinae | Lophosaurus | spinipes | SAMAR37801 | 361 | 35 | 4753 | 5149 | AusARG_Agamidae |
| Agamidae | Amphibolurinae | Moloch | horridus | NMVZ11011 | 186 | 8 | 1488 | 1682 | AusARG_Agamidae |
| Agamidae | Amphibolurinae | Moloch | horridus | NMVZ54529 | 297 | 20 | 3470 | 3787 | AusARG_Agamidae |
| Agamidae | Amphibolurinae | Moloch | horridus | UMMZ244221 | 336 | 26 | 4678 | 5040 | Title & Singhal et al. (2024) |
| Agamidae | Amphibolurinae | Physignathus | coccincinus | ABTC82049 | 365 | 36 | 4741 | 5142 | AusARG_Agamidae |
| Agamidae | Amphibolurinae | Physignathus | coccincinus | SAMAR54798 | 118 | 3 | 1067 | 1188 | AusARG_Agamidae |
| Agamidae | Amphibolurinae | Physignathus | cocincinus | CTMZ04041 | 337 | 0 | 1 | 338 | Burbrink et al. 2021 (AHE_T212) |
| Agamidae | Amphibolurinae | Physignathus | cocincinus | MVZ226496 | 371 | 34 | 4799 | 5204 | Title & Singhal et al. (2024) |
| Agamidae | Amphibolurinae | Pogona | barbata | AMSR.148400 | 360 | 36 | 4722 | 5118 | AusARG_Agamidae |
| Agamidae | Amphibolurinae | Pogona | henrylawsoni | QMJ63887 | 360 | 36 | 4725 | 5121 | AusARG_Agamidae |
| Agamidae | Amphibolurinae | Pogona | henrylawsoni | QMJ84036 | 338 | 30 | 4319 | 4687 | AusARG_Agamidae |
| Agamidae | Amphibolurinae | Pogona | microlepidota | NMVD73864 | 361 | 36 | 4783 | 5180 | AusARG_Agamidae |
| Agamidae | Amphibolurinae | Pogona | microlepidota | WAMR174031 | 275 | 15 | 3199 | 3489 | AusARG_Agamidae |
| Agamidae | Amphibolurinae | Pogona | minor.minima | WAMR112792 | 363 | 35 | 4783 | 5181 | AusARG_Agamidae |
| Agamidae | Amphibolurinae | Pogona | minor.minima | WAMR112800 | 352 | 32 | 4638 | 5022 | AusARG_Agamidae |
| Agamidae | Amphibolurinae | Pogona | minor.minor | UMMZ244225 | 294 | 17 | 4387 | 4698 | Title & Singhal et al. (2024) |
| Agamidae | Amphibolurinae | Pogona | minor.minor | WAMR131084 | 363 | 36 | 4804 | 5203 | AusARG_Agamidae |
| Agamidae | Amphibolurinae | Pogona | minor.minor | WAMR145276 | 364 | 33 | 4730 | 5127 | AusARG_Agamidae |
| Agamidae | Amphibolurinae | Pogona | minor.mitchelli | WAMR113000 | 370 | 35 | 4776 | 5181 | AusARG_Agamidae |
| Agamidae | Amphibolurinae | Pogona | minor.mitchelli | WAMR154527 | 363 | 36 | 4773 | 5172 | AusARG_Agamidae |
| Agamidae | Amphibolurinae | Pogona | nullarbor | SAMAR63054 | 356 | 35 | 4715 | 5106 | AusARG_Agamidae |
| Agamidae | Amphibolurinae | Pogona | nullarbor | WAMR91832 | 364 | 35 | 4741 | 5140 | AusARG_Agamidae |
| Agamidae | Amphibolurinae | Pogona | vitticeps | AMSR.151036 | 360 | 36 | 4755 | 5151 | AusARG_Agamidae |
| Agamidae | Amphibolurinae | Pogona | vitticeps | CTMZ04051 | 358 | 0 | 2 | 360 | Burbrink et al. 2021 (AHE_T212) |
| Agamidae | Amphibolurinae | Pogona | vitticeps | NMVD71349 | 365 | 36 | 4781 | 5182 | AusARG_Agamidae |
| Agamidae | Amphibolurinae | Pogona | vitticeps | SAMEA2300447 | 388 | 41 | 4999 | 5428 | NCBI |
| Agamidae | Amphibolurinae | Pogona | vitticeps | SAMEA2300447 | 388 | 41 | 4999 | 5428 | NCBI |
| Agamidae | Amphibolurinae | Rankinia | diemensis | AMSR.149569 | 358 | 33 | 4706 | 5097 | AusARG_Agamidae |
| Agamidae | Amphibolurinae | Rankinia | diemensis | AMSR.151560 | 362 | 36 | 4718 | 5116 | AusARG_Agamidae |
| Agamidae | Amphibolurinae | Rankinia | diemensis | NMVD71910 | 363 | 35 | 4775 | 5173 | AusARG_Agamidae |
| Agamidae | Amphibolurinae | Rankinia | diemensis | NMVD74157 | 360 | 35 | 4739 | 5134 | AusARG_Agamidae |

*(continued)*

| Family | Subfamily | Genus | Species | RegNo | AHE | gene | uce | total_loci | Source |
|---|---|---|---|---|---|---|---|---|---|
| Agamidae | Amphibolurinae | Rankinia | diemensis | NMVD74165 | 363 | 34 | 4796 | 5193 | AusARG_Agamidae |
| Agamidae | Amphibolurinae | Rankinia | diemensis | UTAS | 364 | 34 | 4710 | 5108 | AusARG_Agamidae |
| Agamidae | Amphibolurinae | Tropicagama | temporalis | AMSR.121164 | 144 | 5 | 674 | 823 | AusARG_Agamidae |
| Agamidae | Amphibolurinae | Tropicagama | temporalis | WAMR112261 | 367 | 35 | 4778 | 5180 | AusARG_Agamidae |
| Agamidae | Amphibolurinae | Tympanocryptis | argillosa | SAMAR49360 | 360 | 35 | 4464 | 4859 | AusARG_Agamidae |
| Agamidae | Amphibolurinae | Tympanocryptis | argillosa | SAMAR54247 | 364 | 32 | 3961 | 4357 | AusARG_Agamidae |
| Agamidae | Amphibolurinae | Tympanocryptis | centralis | NMVZ75749 | 359 | 33 | 4749 | 5141 | AusARG_Agamidae |
| Agamidae | Amphibolurinae | Tympanocryptis | centralis | SAMAR46084 | 348 | 30 | 2648 | 3026 | AusARG_Agamidae |
| Agamidae | Amphibolurinae | Tympanocryptis | centralis | WAMR47369 | 360 | 33 | 3944 | 4337 | AusARG_Agamidae |
| Agamidae | Amphibolurinae | Tympanocryptis | centralis | WAMR50133 | 295 | 21 | 1221 | 1537 | AusARG_Agamidae |
| Agamidae | Amphibolurinae | Tympanocryptis | cephalus | NMVD74070 | 183 | 13 | 304 | 500 | AusARG_Agamidae |
| Agamidae | Amphibolurinae | Tympanocryptis | cephalus | WAMR161060 | 366 | 33 | 4751 | 5150 | AusARG_Agamidae |
| Agamidae | Amphibolurinae | Tympanocryptis | condaminensis | QMJ81871 | 359 | 34 | 4734 | 5127 | AusARG_Agamidae |
| Agamidae | Amphibolurinae | Tympanocryptis | condaminensis | QMJ82087 | 360 | 34 | 4714 | 5108 | AusARG_Agamidae |
| Agamidae | Amphibolurinae | Tympanocryptis | darlingensis | NMVD77147 | 359 | 34 | 4755 | 5148 | AusARG_Agamidae |
| Agamidae | Amphibolurinae | Tympanocryptis | darlingensis | NMVD77158 | 361 | 34 | 4779 | 5174 | AusARG_Agamidae |
| Agamidae | Amphibolurinae | Tympanocryptis | diabolicus | WAMR135412 | 367 | 35 | 4760 | 5162 | AusARG_Agamidae |
| Agamidae | Amphibolurinae | Tympanocryptis | diabolicus | WAMR135413 | 362 | 32 | 4770 | 5164 | AusARG_Agamidae |
| Agamidae | Amphibolurinae | Tympanocryptis | einasleighensis | NMVZ34075 | 360 | 34 | 4774 | 5168 | AusARG_Agamidae |
| Agamidae | Amphibolurinae | Tympanocryptis | einasleighensis | NMVZ34093 | 359 | 36 | 4690 | 5085 | AusARG_Agamidae |
| Agamidae | Amphibolurinae | Tympanocryptis | fictilis | SAMAR46178 | 361 | 36 | 4749 | 5146 | AusARG_Agamidae |
| Agamidae | Amphibolurinae | Tympanocryptis | fictilis | SAMAR58134 | 365 | 36 | 4729 | 5130 | AusARG_Agamidae |
| Agamidae | Amphibolurinae | Tympanocryptis | fortescuensis | WAMR110291 | 360 | 33 | 4760 | 5153 | AusARG_Agamidae |
| Agamidae | Amphibolurinae | Tympanocryptis | fortescuensis | WAMR158076 | 367 | 35 | 4781 | 5183 | AusARG_Agamidae |
| Agamidae | Amphibolurinae | Tympanocryptis | gigas | WAMR175943 | 359 | 32 | 4631 | 5022 | AusARG_Agamidae |
| Agamidae | Amphibolurinae | Tympanocryptis | hobsoni | NMVZ33994 | 361 | 35 | 4745 | 5141 | AusARG_Agamidae |
| Agamidae | Amphibolurinae | Tympanocryptis | hobsoni | NMVZ34020 | 359 | 35 | 4736 | 5130 | AusARG_Agamidae |
| Agamidae | Amphibolurinae | Tympanocryptis | houstoni | SAMAR61480 | 348 | 29 | 4333 | 4710 | AusARG_Agamidae |
| Agamidae | Amphibolurinae | Tympanocryptis | intima | SAMAR42854 | 360 | 35 | 4755 | 5150 | AusARG_Agamidae |
| Agamidae | Amphibolurinae | Tympanocryptis | intima | SAMAR44954 | 281 | 16 | 1086 | 1383 | AusARG_Agamidae |
| Agamidae | Amphibolurinae | Tympanocryptis | intima | SAMAR46815 | 231 | 14 | 690 | 935 | AusARG_Agamidae |
| Agamidae | Amphibolurinae | Tympanocryptis | lineata | NMVZ75948 | 360 | 35 | 4654 | 5049 | AusARG_Agamidae |
| Agamidae | Amphibolurinae | Tympanocryptis | lineata | NMVZ75960 | 366 | 34 | 4740 | 5140 | AusARG_Agamidae |
| Agamidae | Amphibolurinae | Tympanocryptis | lineata | SAMAR38823 | 371 | 33 | 4698 | 5102 | Title & Singhal et al. (2024) |
| Agamidae | Amphibolurinae | Tympanocryptis | macra | NMVD72634 | 361 | 36 | 4740 | 5137 | AusARG_Agamidae |
| Agamidae | Amphibolurinae | Tympanocryptis | macra | NMVD72689 | 364 | 34 | 4717 | 5115 | AusARG_Agamidae |
| Agamidae | Amphibolurinae | Tympanocryptis | macra | NMVD73895 | 358 | 35 | 4704 | 5097 | AusARG_Agamidae |
| Agamidae | Amphibolurinae | Tympanocryptis | macra | NMVD73903 | 344 | 28 | 4541 | 4913 | AusARG_Agamidae |
| Agamidae | Amphibolurinae | Tympanocryptis | mccartneyi | AMSR.186297 | 339 | 31 | 4601 | 4971 | AusARG_Agamidae |
| Agamidae | Amphibolurinae | Tympanocryptis | pentalineata | NMVD74074 | 338 | 24 | 2884 | 3246 | AusARG_Agamidae |
| Agamidae | Amphibolurinae | Tympanocryptis | pentalineata | NMVD74075 | 230 | 16 | 572 | 818 | AusARG_Agamidae |
| Agamidae | Amphibolurinae | Tympanocryptis | petersi | NMVD74164 | 361 | 35 | 4747 | 5143 | AusARG_Agamidae |
| Agamidae | Amphibolurinae | Tympanocryptis | petersi | NMVD74168 | 354 | 30 | 4115 | 4499 | AusARG_Agamidae |
| Agamidae | Amphibolurinae | Tympanocryptis | pinguicolla | NMVZ75954 | 361 | 34 | 4707 | 5102 | AusARG_Agamidae |
| Agamidae | Amphibolurinae | Tympanocryptis | pseudopsephos | WAMR106201 | 333 | 26 | 2323 | 2682 | AusARG_Agamidae |
| Agamidae | Amphibolurinae | Tympanocryptis | pseudopsephos | WAMR146827 | 316 | 24 | 3945 | 4285 | AusARG_Agamidae |
| Agamidae | Amphibolurinae | Tympanocryptis | rustica | NMVD74671 | 364 | 33 | 4743 | 5140 | AusARG_Agamidae |
| Agamidae | Amphibolurinae | Tympanocryptis | rustica | SAMAR38823 | 337 | 29 | 4048 | 4414 | AusARG_Agamidae |
| Agamidae | Amphibolurinae | Tympanocryptis | tetraporophora | ANWC | 359 | 34 | 4708 | 5101 | AusARG_Agamidae |
| Agamidae | Amphibolurinae | Tympanocryptis | tetraporophora | NMVD74046 | 357 | 34 | 4753 | 5144 | AusARG_Agamidae |
| Agamidae | Amphibolurinae | Tympanocryptis | tolleyi | SAMAR26711 | 364 | 36 | 4681 | 5081 | AusARG_Agamidae |
| Agamidae | Amphibolurinae | Tympanocryptis | wilsoni | QMJ87307 | 357 | 35 | 4741 | 5133 | AusARG_Agamidae |
| Agamidae | Amphibolurinae | Tympanocryptis | wilsoni | QMJ89119 | 353 | 31 | 4564 | 4948 | AusARG_Agamidae |
| Agamidae | Draconinae | Acanthosaura | armata | lsuhc9351 | 16 | 0 | 4128 | 4144 | Title & Singhal et al. (2024) |
| Agamidae | Draconinae | Acanthosaura | coronata | FMNH262964 | 359 | 35 | 4798 | 5192 | Title & Singhal et al. (2024) |

(continued)

| Family | Subfamily | Genus | Species | RegNo | AHE | gene | uce | total_loci | Source |
|---|---|---|---|---|---|---|---|---|---|
| Agamidae | Draconinae | Acanthosaura | crucigera | LSUHC9913 | 354 | 0 | 1 | 355 | Burbrink et al. 2021 (AHE_T212) |
| Agamidae | Draconinae | Acanthosaura | lepidogaster | cwl818 | 14 | 0 | 3873 | 3887 | Title & Singhal et al. (2024) |
| Agamidae | Draconinae | Acanthosaura | nataliae | MVZHerp258141 | 349 | 30 | 4659 | 5038 | Title & Singhal et al. (2024) |
| Agamidae | Draconinae | Aphaniotis | acutirostris | MVZHerp239462 | 351 | 35 | 4820 | 5206 | Title & Singhal et al. (2024) |
| Agamidae | Draconinae | Bronchocela | cristatella | acd1547 | 15 | 0 | 3961 | 3976 | Title & Singhal et al. (2024) |
| Agamidae | Draconinae | Bronchocela | jubata | MVZHerp253996 | 359 | 35 | 4824 | 5218 | Title & Singhal et al. (2024) |
| Agamidae | Draconinae | Bronchocela | smaragdina | FMNH262309 | 355 | 35 | 4735 | 5125 | Title & Singhal et al. (2024) |
| Agamidae | Draconinae | Calotes | emma | CAS230941 | 27 | 27 | 4425 | 4479 | Title & Singhal et al. (2024) |
| Agamidae | Draconinae | Calotes | emma | CTMZ04032 | 348 | 0 | 2 | 350 | Burbrink et al. 2021 (AHE_T212) |
| Agamidae | Draconinae | Calotes | mystaceus | dsm869 | 15 | 0 | 3758 | 3773 | Title & Singhal et al. (2024) |
| Agamidae | Draconinae | Calotes | versicolor | lsuhc10327 | 16 | 0 | 4093 | 4109 | Title & Singhal et al. (2024) |
| Agamidae | Draconinae | Calotes | versicolor | SAMN07676179 | 360 | 36 | 3848 | 4244 | NCBI |
| Agamidae | Draconinae | Ceratophora | aspera | AF128491 | 15 | 0 | 3303 | 3318 | Title & Singhal et al. (2024) |
| Agamidae | Draconinae | Cophotis | ceylanica | AF128493 | 15 | 0 | 3739 | 3754 | Title & Singhal et al. (2024) |
| Agamidae | Draconinae | Cristidorsa | planidorsata | CAS219935 | 358 | 34 | 4778 | 5170 | Title & Singhal et al. (2024) |
| Agamidae | Draconinae | Diploderma | yunnanense | CAS242183 | 360 | 34 | 4806 | 5200 | Title & Singhal et al. (2024) |
| Agamidae | Draconinae | Draco | blanfordii | lsuhc9427 | 15 | 0 | 3838 | 3853 | Title & Singhal et al. (2024) |
| Agamidae | Draconinae | Draco | maculatus | kufs320 | 15 | 0 | 3906 | 3921 | Title & Singhal et al. (2024) |
| Agamidae | Draconinae | Draco | spilopterus | elr1338 | 13 | 0 | 3896 | 3909 | Title & Singhal et al. (2024) |
| Agamidae | Draconinae | Draco | walkeri | MVZHerp239464 | 342 | 32 | 4691 | 5065 | Title & Singhal et al. (2024) |
| Agamidae | Draconinae | Gonocephalus | abbotti | FMNH230175 | 368 | 34 | 4806 | 5208 | Title & Singhal et al. (2024) |
| Agamidae | Draconinae | Gonocephalus | doriae | FMNH230175 | 368 | 34 | 4790 | 5192 | Singhal et al. 2021 |
| Agamidae | Draconinae | Gonocephalus | interruptus | KU314925 | 341 | 0 | 1 | 342 | Burbrink et al. 2021 (AHE_T212) |
| Agamidae | Draconinae | Gonocephalus | interruptus | rmb9384 | 15 | 0 | 4041 | 4056 | Title & Singhal et al. (2024) |
| Agamidae | Draconinae | Gonocephalus | sophiae | rmb8061 | 14 | 0 | 3975 | 3989 | Title & Singhal et al. (2024) |
| Agamidae | Draconinae | Lyriocephalus | scutatus | AF128494 | 16 | 0 | 3586 | 3602 | Title & Singhal et al. (2024) |
| Agamidae | Draconinae | Mantheyus | phuwuanensis | fmnh262580 | 15 | 0 | 4072 | 4087 | Title & Singhal et al. (2024) |
| Agamidae | Draconinae | Pseudocalotes | kingdonwardi | CAS241965 | 364 | 33 | 4786 | 5183 | Title & Singhal et al. (2024) |
| Agamidae | Draconinae | Pseudocalotes | larutensis | lsuhc10285 | 15 | 0 | 3842 | 3857 | Title & Singhal et al. (2024) |
| Agamidae | Draconinae | Pseudocalotes | microlepis | FMNH258710 | 357 | 32 | 4843 | 5232 | Title & Singhal et al. (2024) |
| Agamidae | Draconinae | Ptyctolaemus | collicristatus | cas220561 | 17 | 0 | 3882 | 3899 | Title & Singhal et al. (2024) |
| Agamidae | Draconinae | Ptyctolaemus | gularis | cas221296 | 15 | 0 | 4041 | 4056 | Title & Singhal et al. (2024) |
| Agamidae | Draconinae | Salea | horsfieldii | AF128491 | 16 | 0 | 3979 | 3995 | Title & Singhal et al. (2024) |
| Agamidae | Leiolepinae | Leiolepis | belliana | CAS210725 | 42 | 28 | 4456 | 4526 | Singhal et al. 2021 |
| Agamidae | Leiolepinae | Leiolepis | belliana | LSUHC6842 | 350 | 0 | 1 | 351 | Burbrink et al. 2021 (AHE_T212) |
| Agamidae | Amphibolurinae | Lophosaurus | dilophus | WAMR109630 | 365 | 35 | 4665 | 5065 | AusARG_Agamidae |
| Agamidae | Amphibolurinae | Tympanocryptis | osbornei | AA087670 | 365 | 34 | 4693 | 5092 | AusARG_Agamidae |
| Agamidae | Amphibolurinae | Tympanocryptis | osbornei | AA61541 | 360 | 34 | 4703 | 5097 | AusARG_Agamidae |
| Agamidae | Uromastycinae | Uromastyx | sp | CTMZ12648 | 167 | 0 | 0 | 167 | Burbrink et al. 2021 (AHE_T212) |
| Teiidae | — | Salvator | merianae | SAMN09273531 | 388 | 41 | 5000 | 5429 | NCBI |
| Scincidae | Lygosominae_Eugongylini | Cryptoblepharus | egeriae | SAMN32772511 | 388 | 40 | 5003 | 5431 | NCBI |
| Elapidae | — | Hydrophis | elegans | SAMN35787919 | 385 | 39 | 4859 | 5283 | NCBI |
| Agamidae | Hydrosaurinae | Hydrosaurus | sp | JAM848 | — | — | 2718 | 2718 | Streicher_Iguania |

**Table S2. Fossil Calibrations**

| Fossil Information | Calibration | Split/Position | Source |
|---|---|---|---|
| Uniform—Sophineta | ‘B(2.38,2.55)’ | Lepidosauria (Sphenodon + Squamata) | see below |
| Uniform—Secondary | ‘B(0.50,0.80)’ | Amphibolurinae + (Agaminae + Draconinae) | Burbrink et al. 2020; Title et al. 2024 |
| Uniform—Riversleigh ‘Physignathus’ | ‘B(0.17,0.50)’ | Hypsilurus + Remaining Australian Amphibolurinae | Covacevich 1990 |

*Note.* The root divergence between lepidosaurs (Sphenodon + squamates) is based on the fossil taxa Sophineta cracoviensis (Evans & Bialynicka, 2009), Megachirella wachtleri (Renesto & Posenato, 2003), and the Vellberg Jaw (Jones et al., 2013). These taxa represent stem squamates and rhynchocephalians, and so provide a soft lower bound on the crown divergence of Lepidosauria, with a soft upper bound provided by Protosaurus speneri (von Meyer, 1832).

**Table S3. Morphological Measurements**

| No. | Measurement | Abbreviation | Method |
|---|---|---|---|
| 1 | Snout-vent length | SVL | From the tip of the snout to the vent. |
| 2 | Snout-axilla length | SAL | From the tip of the snout to the midpoint of the crease between the fore-limb and the body on the ventral surface. |
| 3 | Inter-limb length | ILL | Midpoint of the crease on the ventral surface where the fore-limb connects to the body, to the midpoint of the crease on the ventral surface where the hind-limb connects to the body. |
| 4 | Body width | BW | From one lateral side of the body to the other, where possible at the midpoint of the ILL. |
| 5 | Pelvic width | PW | From the midpoint of the crease on the ventral surface where the left hind limb connects to the body, to the midpoint of the crease on the ventral surface where the right hind limb connects to the body. |
| 6 | Pelvic height | PH | From the top of the dorsal surface were the PW was measured, to the bottom of the ventral surface where the PW was measured. |
| 7 | Head width | HW | Widest part of the head from one dorsal-lateral edge to the other edge. |
| 8 | Head length | HL | From the nose tip, to the anterior of the ear. |
| 9 | Snout length | SN | From the nasal opening to the anterior of the eye. |
| 10 | Eye diameter | ED | From one side of the eye to the other. |
| 11 | Head depth | HD | From the top of the tallest part of the head on the dorsal surface, to the bottom of the ventral surface under the jaw. |
| 12 | Tail width | TW | Measured at the vent, from one dorsal-lateral edge to the other edge. |
| 13 | Tail length | TL | Measured from the vent to the tip of the tail. |
| 14 | Fore-limb length | FLL | Measured fully extended from the midpoint of the crease on the ventral surface where the front limb connects to the body, to the end of longest toe (claw included). |
| 15 | Front foot | FFOOT | From the base of the foot to the end of the longest toe (claw included). |
| 16 | Lower front limb | LFL | Measured from the base of the lower fore limb to the juncture where the limb meets the front foot. |
| 17 | Upper front limb | UFL | Measured from the crease on the ventral surface where the fore limb connects to the body, to the end of the lower front limb. |
| 18 | Hind-limb length | HLL | From the midpoint of the crease on the ventral surface where the hind limb connects |

| No. | Measurement | Abbreviation | Method |
| --- | --- | --- | --- |
| | | | to the body, to the end of longest toe (claw included) |
| 19 | Hind foot | HFOOT | From the base of the hind foot to the end of the longest toe (claw included) |
| 20 | Lower hind limb | LHL | Measured from the top of the knee joint to the heel juncture where the limb meets the front foot. |
| 21 | Upper hind limb | UHL | Measured from the crease on the ventral surface where the hind limb connects to the body, to the end of the knee. |

**Table S3. Morphological measurements collected (21).**

| No. | Measurement | Shorthand | Method |
|---|---|---|---|
| 3 | Interlimb length | Interlimb | see Table S3 |
| 4 | Body width | Body__Width | see Table S3 |
| 5 | Pelvic width | Pelvic__Width | see Table S3 |
| 6 | Pelvic height | Pelvic__Height | see Table S3 |
| 7 | Head width | Head__Width | see Table S3 |
| 9 | Snout length | Snout__Eye | see Table S3 |
| 10 | Eye diameter | Eye__Diameter | see Table S3 |
| 11 | Head depth | Head__Depth | see Table S3 |
| 12 | Tail width | Tail__Width | see Table S3 |
| 13 | Tail length | Tail__Length | see Table S3 |
| 17 | Upper front limb | Upper__Arm | see Table S3 |
| 16 | Lower front limb | Lower__Arm | see Table S3 |
| 15 | Front foot | Hand | see Table S3 |
| 21 | Upper hind limb | Upper__Leg | see Table S3 |
| 20 | Lower hind limb | Lower__Leg | see Table S3 |
| 19 | Hind foot | Foot | see Table S3 |
| 22 | Neck length | Neck | Snout__Axilla - Head__Length |
| 23 | Posterior skull | Pos__Skull | Head__Length - (Snout__Eye + Eye__Diameter) |
| 24 | Pelvic gap | Pelvic__Gap | Snout__Vent - (Interlimb + Snout__Axilla) |

**Table S3. Final morphological traits used for phenotypic analyses (19).**

#### Table S4. BayesTraits Model Fitting Results

| trait | fabric | VarRates | fabric_noV | fabric_noB | BM | best_model | variance | disparity | func_div | func_even | BM_rate_constant | delta_fabric_BM |
|---|---|---|---|---|---|---|---|---|---|---|---|---|
| Size | 4.347283 | -10.49492 | -8.786684 | -10.34354 | -12.12269 | fabric_full | 1.8677873 | 0.3648166 | 0.6508999 | 0.5574674 | 0.0036639 | 16.469968 |
| TailLength | 21.370924 | 11.69656 | 11.032386 | 10.25499 | 14.21709 | fabric_full | 1.6292760 | 0.1652216 | 0.6571144 | 0.4853009 | 0.0023428 | 7.153839 |
| PelvicWidth | 84.614148 | 74.17618 | 74.079081 | 74.42456 | 64.93564 | fabric_full | 1.1947444 | 0.0736472 | 0.5838028 | 0.4392344 | 0.0009712 | 19.678503 |
| BodyWidth | 34.199465 | 26.17027 | 23.228444 | 26.83045 | 30.12197 | fabric_full | 1.0949751 | 0.1191263 | 0.6717122 | 0.5590405 | 0.0017699 | 4.077491 |
| SnoutEye | 60.264876 | 53.71249 | 51.692876 | 54.61451 | 58.84224 | fabric_full | 1.0117128 | 0.0788389 | 0.6385742 | 0.5250549 | 0.0010726 | 1.422641 |
| Foot | 87.587220 | 84.08598 | 78.497124 | 83.88982 | 83.88980 | fabric_full | 0.9995301 | 0.0537720 | 0.6418632 | 0.4967055 | 0.0007005 | 3.697424 |
| PosSkull | 96.608958 | 93.10331 | 85.893212 | 91.57044 | 90.83893 | fabric_full | 0.9450553 | 0.0360397 | 0.6072598 | 0.4167057 | 0.0006203 | 5.770026 |
| TailWidth | 53.013935 | 50.10927 | 43.404429 | 50.73475 | 52.92838 | fabric_full | 0.8857337 | 0.0672043 | 0.6161991 | 0.5656558 | 0.0011863 | 0.085553 |
| HeadWidth | 91.758160 | 88.38046 | 79.674046 | 89.04225 | 89.51111 | fabric_full | 0.7587556 | 0.0350892 | 0.6719401 | 0.4819992 | 0.0006389 | 2.247051 |
| HeadDepth | 87.440963 | 85.72778 | 76.768466 | 86.10532 | 88.04195 | BM | 0.7012197 | 0.0278678 | 0.6198712 | 0.5107771 | 0.0006508 | -0.600983 |
| Neck | 73.415897 | 70.07112 | 63.424944 | 67.14744 | 71.54452 | fabric_full | 0.6968378 | 0.0238528 | 0.6054567 | 0.4806514 | 0.0008705 | 1.871379 |
| PelvicHeight | 105.392370 | 104.55769 | 96.044915 | 102.05399 | 108.33293 | BM | 0.6404908 | 0.0213563 | 0.5840434 | 0.4554133 | 0.0004596 | -2.940564 |
| EyeDiameter | 99.429377 | 97.79872 | 90.032312 | 98.48655 | 102.25141 | BM | 0.6132040 | 0.0221819 | 0.6495101 | 0.5091384 | 0.0005100 | -2.822032 |
| UpperArm | 125.932379 | 123.22714 | 115.607148 | 123.15233 | 126.81728 | BM | 0.5553347 | 0.0158573 | 0.6035020 | 0.4625320 | 0.0003338 | -0.884904 |
| LowerLeg | 128.305167 | 127.30274 | 118.832284 | 125.79280 | 130.54499 | BM | 0.5338807 | 0.0247560 | 0.6182549 | 0.5323700 | 0.0003138 | -2.239819 |
| Hand | 130.519791 | 129.97166 | 120.419049 | 130.71713 | 130.87166 | BM | 0.5040150 | 0.0095631 | 0.6086064 | 0.4474384 | 0.0003099 | -0.351867 |
| LowerArm | 150.707693 | 150.03209 | 141.357799 | 148.94942 | 152.24296 | BM | 0.4366660 | 0.0123221 | 0.6208814 | 0.5195238 | 0.0002146 | -1.535267 |
| UpperLeg | 128.500791 | 128.39444 | 119.263162 | 128.65201 | 132.68701 | BM | 0.4265069 | 0.0168591 | 0.6497425 | 0.6060006 | 0.0002977 | -4.186218 |
| Interlimb | 120.023773 | 116.03700 | 110.894019 | 119.12322 | 123.54082 | BM | 0.3945153 | 0.0145229 | 0.6099829 | 0.5718187 | 0.0003523 | -3.517043 |
